# The Landscape of Stop Codon-Free Regions in Primates: A Reservoir of Proto-Genes

**DOI:** 10.64898/2026.02.27.708503

**Authors:** Aswin S Soman, G Shreyasree, Aishwarya Dwivedi, Gayathri S Pramod, Chirayu Sakarkar, Dipannita Bhattacharya, Nagarjun Vijay

**Affiliations:** Department of Biological Sciences, IISER Bhopal, Bhauri, Madhya Pradesh, India; Department of Biotechnology, Dayananda Sagar College of Engineering, Bangalore, Karnataka, India; Department of Biotechnology, Bennett University, Greater Noida, Uttar Pradesh, India; Department of Microbiology, Maulana Azad College, University of Calcutta, Kolkata, West Bengal, India

**Keywords:** stop-codon-free regions, primates, *de novo* gene birth, Proto-gene, exon-shadow, exitron, codon periodicity, GC content

## Abstract

Gene duplication has long been viewed as the primary source of new genes, yet growing evidence suggests that de novo emergence from non-coding DNA may be more common than previously assumed, requiring unbiased genome-wide strategies to identify its structural precursors. New protein-coding genes can arise from non-coding DNA, but the sequence features enabling this transition remain unclear. Here, we systematically identify and characterise stop-codon-free regions (SCFRs) across telomere-to-telomere assemblies of human and six other primates. Short SCFRs are abundant and widely distributed, whereas long SCFRs are rare and increasingly associated with coding overlap, moderate GC enrichment, and structured exon–intron contexts. We define exon shadows as in-frame SCFR extensions beyond annotated exon boundaries that lack stop codons, revealing latent coding-compatible sequence adjacent to established exons. We also detect introns fully spanned by single SCFRs, consistent with exitron-like architectures. Repeat composition, codon usage, and Fourier spectral analyses show that length filtering enriches for gene-like features and identifies a subset of long SCFRs with codon-scale periodicity. Together, these findings provide a framework for identifying extended ORF-like regions that may serve as substrates for de novo gene emergence in primates.

## 1. Introduction

The emergence of new protein-coding genes has been a key topic in genome biology and molecular evolution. Traditionally, it was believed that most new genes arose through the process of gene duplication followed by functional divergence [1]. This ensured that even though the formation of new functional genes took place, the ancestral gene’s function was retained. However, recent theories suggest a new method by which these new protein-coding genes arise: *de novo* gene emergence. These theories also suggest that *de novo* gene birth from non-coding DNA is quite a common and ongoing process across various lineages, including humans [2–4]. These *de novo* genes originate when non-coding sequences begin to be transcribed and translated, forming proto genes that may be shaped by natural selection and sometimes preserved for their beneficial functions [5]. Most studies which focus on *de novo* genes discuss young genes which lack a specific identifiable homolog. *De novo* gene birth, once described as a highly unlikely phenomenon, now has evidence of its existence across various species. It has also been studied that *de novo* gene birth plays a very important role in evolutionary modifications that occur across species [6].

For a non-coding region present in a genome to convert to a protein-coding gene, it must first acquire the ability to be transcribed into RNA and then subsequently into a functional protein. Acquisition of a promoter, or the translocation of the region to a locus downstream of a pre-existing promoter, can facilitate transcription [7]. Once transcribed, translation into protein requires the presence of an open reading frame (ORF) with a start codon (typically AUG in eukaryotes) and a sufficiently long downstream sequence that is free of in-frame stop codons. These stop codon-free regions allow the ribosome to continue elongation, potentially generating novel peptides [3]. Analysing cases of *de novo* gene emergence shows that singular mutations can enable the production of a stable RNA transcript from genomic regions that were formerly silent, lacking both transcription and translation activities [8]. In this context, long stop-codon-free regions (SCFRs) within intergenic and intronic DNA are of particular interest. These regions are considered a vast and largely unexplored collection of “proto-genes” which are transient ORFs that may be non-functional initially but can, through mutation and selection, evolve into fully functional protein-coding genes [9].

Proto-genes serve as a valuable source of evolutionary innovation, particularly when their translation results in peptides that are either not deleterious or occasionally beneficial. Recent studies using ribosome profiling and comparative genomics have revealed widespread translation of non-coding regions, further supporting the idea that *de novo* gene birth is an important ongoing process in genomes across various taxa [10,11]. These findings suggest that SCFRs, specifically the ones present in regions with accessible chromatin or low levels of purifying selection, may be hotspots for the emergence of new protein-coding genes. A major challenge in the formation of extended ORFs from non-coding sequences is the high frequency of stop codons, which often truncates translation prematurely. However, the uneven distribution of nucleotides, especially in regions with biased base composition, can result in extended stretches lacking in-frame stop codons in one or more reading frames [12]. For example, regions with high adenine-thymine (AT) content tend to have shorter SCFRs, as the probability of forming stop codons (TAA, TAG, TGA) is increased. However, regions which are richer in GC content tend to have longer stretches of SCFRs, thus showing that these regions can arise at random and persist over evolutionary timescales, providing a fertile ground for translation and potential functionalisation.

De novo gene birth has been investigated across a wide range of species to acquire a better understanding of the molecular and evolutionary mechanisms by which non-coding sequences give rise to novel protein-coding genes [2,13]. These studies show that many genomes have a large number of lineage-specific genes with no discernible homologs, many of which appear to have originated from non-coding DNA. However, challenges have been encountered while trying to conduct comprehensive genome analyses due to fragmented and error-prone assemblies, which often lead to ambiguities in results [14,15]. The recent advent of telomere-to-telomere (T2T) assemblies of human and several non-human primates marks a turning point in our ability to explore the whole genomic landscape and study SCFRs in greater detail [13,14,16]. Because these assemblies are gapless, they provide a remarkable view of genomic regions which were previously inaccessible, allowing researchers to study and understand proto-genes better. We leverage these T2T assemblies in humans and various other primates to identify and evaluate SCFRs throughout the genome to examine their sequence composition, overlap with known and putative coding exons, and the potential to encode novel proteins. We also examine the potential reasons behind the occurrence of these long SCFRs, including the involvement of various transposons in these genomic regions. Together, this work sheds light on the raw genomic substrate from which new genes can emerge and offers new insights into the early stages of de novo gene evolution in primate lineages.

## 2. Materials and methods

### 2.1. Identification of Stop-Codon-Free Regions (SCFRs)

To comprehensively identify stop-codon-free regions (SCFRs) in the genome, we developed and employed a custom Python script (*find_stop_codon_free_regions_with_reverse.py*, **see Supplementary data-scripts**) that systematically scans both DNA strands of a given genome assembly in all six possible reading frames. The script accepts a genome FASTA file as input and parses each chromosome or scaffold using Biopython [17]. For every sequence, it scans the forward strand in all three reading frames (+1, +2, +3) and the reverse complement strand in the corresponding reverse frames (−1, -2, -3). The script iteratively examines every triplet (codon) in each frame and identifies contiguous regions that do not contain any in-frame stop codons (TAA, TAG, or TGA). Each such region is defined as an SCFR and recorded along with its genomic coordinates and reading frame. For reverse-strand regions, the script internally computes the reverse complement of the sequence and then remaps coordinates to match the original forward-strand orientation. All identified regions are reported in BED format with four columns: chromosome name, start position (0-based), end position (exclusive), and frame (positive for forward strand and negative for reverse strand). Gaps in the genome represented as N’s are excluded from the analysis and reported in the gap file in BED format.

Genome assemblies and annotation of seven primate species were downloaded from the NCBI genome database [18]. The genome assemblies used are human (T2T-CHM13v2.0 [GCA_009914755.4]), chimpanzee (NHGRI_mPanTro3-v2.1_pri [GCA_028858775.3]), bonobo (NHGRI_mPanPan1-v2.1_pri [GCA_029289425.3]), gibbon (NHGRI_mSymSyn1-v2.1_pri [GCA_028878055.3]), gorilla (NHGRI_mGorGor1-v2.1_pri [GCA_029281585.3]), Bornean orangutan (NHGRI_mPonPyg2-v2.1_pri [GCA_028885625.3]), and Sumatran orangutan (NHGRI_mPonAbe1-v2.1_pri[GCA_028885655.3]). The SCFR finder script was run on each genome to ensure complete and unbiased coverage of genic and intergenic regions, including repetitive and previously unresolved sequences. This procedure yielded a comprehensive catalogue of SCFRs across each genome, facilitating downstream analyses of potential proto-genes.

### 2.2. Filtering Annotated Coding Exons from SCFRs

We removed all regions overlapping known coding exons using the RefSeq annotation to ensure that the identified SCFRs were not simply annotated protein-coding exons. The coding exon coordinates were extracted from the RefSeq GTF file using a custom Python script (*gtf_to_bed.py*, see **supplementary data-scripts**), which selectively parsed and converted all CDS (coding sequence) entries to BED format [19]. To exclude all SCFRs that overlap known coding exons, we used bedtools [20] intersect with the -v flag, which reports only the entries in the first file that have no overlap with the second. This filtering step produced a curated list of SCFRs entirely outside known protein-coding exons, enabling downstream analyses focused on unannotated or potentially novel translated regions, such as proto-genes or *de novo* open reading frames.

### 2.3. Identification of gene deserts

To systematically identify large intergenic regions, commonly referred to as gene deserts [21], we developed a custom Python workflow designed to analyse BED files containing genomic coordinates of all annotated protein-coding exons (*gene_desert_finder.py*; see **Supplementary data-scripts**). In the first step, the pipeline merges all exons belonging to the same gene, thereby creating a single continuous interval representing the entire gene, including introns. In the second step, these gene-level intervals are further merged across genes whenever adjacent or overlapping exons are detected, ensuring that no closely spaced genes are erroneously classified as intergenic. The script then iteratively scans each chromosome and calculates the genomic gap between every pair of adjacent merged intervals. For all intergenic regions, length-based Z-scores are computed, and a threshold of Z ≥ 2 is applied. This threshold selectively retains only those regions that are at least two standard deviations longer than the genome-wide mean, representing statistically unusually long gene-free intervals. The pipeline produces two output tables: one listing all intergenic regions, and another containing only those that satisfy the Z-score criterion and are therefore defined as putative gene deserts.

### 2.4. Quantification of codon usage patterns and PCA

To investigate compositional biases, including GC content and codon usage patterns within each species, we analysed SCFRs from all seven genomes. Because SCFRs collectively cover most of each genome and show substantial variation in length, we applied length-based filtering at 2.5kb, 5 kb, 7.5 kb, and 10 kb to enable more consistent comparisons. We then further classified SCFRs by their overlap with protein-coding genes, producing three subsets: (1) SCFRs overlapping coding regions and sharing the same frame, (2) SCFRs overlapping coding regions with a different frame, and (3) SCFRs that show no overlap with any annotated coding region. For each species, filtered SCFR sequences were extracted chromosome-wise and processed using a custom Python script (*corrected_rscu_calc.py*, **Supplementary data-scripts**). The script parses each FASTA file, validates ORFs, and computes three classes of metrics: global metrics, regional metrics, and relative synonymous codon usage (RSCU), which are written to separate .tsv files. For the global metrics, it first checks whether each sequence represents a valid ORF by confirming that it begins with a canonical start codon, ends with a valid stop codon, and contains a clean reading frame without premature stops. It then calculates whole-sequence GC content, enumerates extended GC stretches (runs of ≥5 G or C nucleotides), and counts every codon across the entire ORF, allowing us to characterise the codon composition of each SCFR. To capture regional variation, the script applies a 60 bp sliding window with 30 bp overlap and quantifies the GC content and GC-stretch frequency within each window, enabling detection of local compositional biases that may be obscured at the global scale. Finally, the script computes Relative Synonymous Codon Usage (RSCU) by comparing the observed codon count of each synonymous family to its expected frequency, providing a normalised measure of codon bias suitable for downstream multivariate analyses such as PCA. Together, these metrics allow detailed quantification of sequence composition and codon-usage patterns across all SCFRs in each species.

The Principal Component Analysis (PCA) and K-means clustering were performed using R scripts (*pca_script.R*, *kmeans_script.*R; **Supplementary data-scripts**) to visualise codon-usage variation among species based on their RSCU profiles. The script first reshaped the RSCU values into a chromosome-by-SCFR matrix and removed codons exhibiting zero variance across all SCFRs, ensuring that only informative features contributed to the analysis. The remaining codon variables were then standardised so that each codon contributed equally to the PCA regardless of its absolute magnitude. For each pairwise combination of the first five principal components (PC1 to PC5), the script generated high-resolution PCA plots, colouring points by chromosome and displaying SCFR labels to facilitate group-level comparisons. To support biological interpretation, the script also computed codon loadings and highlighted the top five codons influencing each PC pair by projecting their vectors onto the PCA space. These visualisations enabled clear identification of species clusters based on similarities and differences in codon usage. Relative codon contributions to PC1, PC2, and PC1+PC2 were calculated from the squared loadings. Amino acid properties, including GC3 content, hydrogen bonding, polarity, charge, chemical class, volume, and hydropathy, were assigned using the standard genetic code and reference property tables in R for visualisation (*plot_codon_contribution_heatmap.R*, **Supplementary data-scripts**).

To identify potential proto-genes within SCFRs, open reading frames (≥600 bp) were extracted, clustered at 70% sequence identity using CD-HIT [22] to obtain non-redundant representatives, and queried against the NCBI nr database using BLASTP (version 2.9.0; e-value ≤ 1e−4; query coverage ≥ 70%) to assess homology [23]. To assess the contribution of repetitive elements to long SCFRs, we analysed SCFR sequences ≥1000 bp using RepeatMasker v4.1.7 (https://www.repeatmasker.org). For each species, SCFR FASTA files were identified from the length-filtered datasets and processed individually. RepeatMasker was executed with the following parameters: -pa 8 (8 parallel threads), -a (alignment output), -s (sensitive mode), -u (unaligned output), -gff (GFF annotation output), -html (HTML summary output), and -xsmall (soft-masked output). Species-specific repeat libraries were used for chimpanzee, gorilla, and human, while the Hominidae repeat library was applied to the remaining primate genomes.

### 2.5. Discrete Fourier Transform Analysis

To identify periodic patterns embedded within nucleotide sequences, we developed a Python script (*scfr_parallel_fft_motif_report_grouped.py*, see **Supplementary data-scripts**), which applies the Fourier transformation to translate linear sequence information into its corresponding frequency spectrum. Fourier analysis is appropriate for this purpose because repeating biological patterns of any kind manifest as distinct frequencies whose strengths can be quantitatively evaluated. Each sequence was first converted into four binary NumPy arrays marking the positions of A, C, G and T, and these arrays provided the numerical basis for subsequent frequency analysis [24]. For each binary array, the discrete Fourier transform was computed using NumPy’s FFT implementation, and magnitudes were summed across nucleotides to yield a combined spectrum that strengthens shared periodic signals, such as codon-related patterns that influence all bases rather than individual nucleotides alone. In the resulting spectrum, frequency denotes how often a specific pattern recurs along the sequence, whereas amplitude measures the intensity of that repeating signal within the nucleotide arrangement. The script then plots magnitude versus frequency and identifies spectral peaks that correspond to prominent repeating patterns, with each peak’s position translated into a characteristic period (e.g., a period of 3 for codons). Based on these peaks, sequences are classified into three groups: a single dominant peak near 0.33, indicating strong codon-level periodicity, a single peak at another frequency suggesting alternative periodic features, and multiple peaks reflecting mixed or complex repeating structures. The script then used the frequency peak with the highest amplitude to infer the underlying period and extract the most frequent motifs associated with it, capturing the dominant repeating structural patterns encoded in each sequence. All results, including spectra, classifications, motif summaries and sequence metadata, were compiled into a unified PDF report generated using LaTeX.

Fourier-derived frequency spectra were analysed using kernel density estimation (KDE) to characterise their distribution (*plot_fourier_frequencies.R*, **Supplementary data-scripts**). For each spectrum, the two dominant local maxima and the intervening minimum were identified, and the separation between peaks was quantified. This procedure was applied across all species using both the complete SCFR dataset and length-filtered subsets (≥5 kb, ≥7.5 kb, and ≥10 kb), with coding-overlapping and non-coding SCFRs analysed separately.

### 2.6. Genome -wide Identification, Analysis, and Visualisation of Exon Shadows and Exitron Candidates

Exon shadows were identified by intersecting SCFRs with protein-coding exons that shared the same reading frame using a custom shell script (*exon_shadow_analysis.sh*, **Supplementary data-scripts**). Gene annotations were obtained from RefSeq GTF files downloaded from the NCBI database. Only exons that were fully contained within an SCFR or that shared a boundary with an SCFR in the same frame were retained for shadow length calculation. Upstream and downstream shadow lengths were defined as the distance between the exon boundary and the nearest SCFR boundary in the corresponding direction. For each exon-SCFR overlap, additional exon features were recorded, including exon count (number of exons overlapping an SCFR), splicing status (constitutive, alternative, or unique), exon order within the transcript (first, middle, or last), and transcript count (number of transcripts containing the exon). In cases where multiple exons overlapped a single SCFR in the same frame, the intervening intronic sequence became part of the SCFR. Such intronic segments, lacking in-frame stop codons and sharing the exon frame, were classified as putative exitron candidates.

Strand asymmetry and directional shadow asymmetry were quantified using the ratios (Forward-Reverse)/(Forward+Reverse) and (Upstream-Downstream)/(Upstream+Downstream), respectively. Sequence composition metrics, including GC content, GC skew, GC3, and the average lengths of GC and AT stretches, were computed using a custom shell script (*get_composition.sh*, **Supplementary data-scripts**) utilising the SeqKit tool [25]. All statistical analyses and visualisations were generated using R.

To assess whether exon-SCFR boundary offsets occur randomly, enrichment of shadow lengths was evaluated separately for upstream and downstream shadows using a custom R script (*enrichment_shadow_length.R*, **Supplementary data-scripts**). Shadow lengths were discretised at single-base resolution, and observed frequencies were compared against a uniform null model in which all tested lengths were assumed to be equally likely. Enrichment was assessed using binomial tests, and p-values were corrected for multiple testing using the Benjamini–Hochberg false discovery rate procedure.

### 2.7. Gene Set Enrichment Analysis

Gene set enrichment analysis was performed using ShinyGO [26]. For each species, five gene sets were analysed: genes with symmetric shadows (GWSS), genes lacking downstream shadows (GWOD), genes lacking upstream shadows (GWOU), genes without shadows (GWOS), and genes with shadows (GWS). All analyses used the default background of protein-coding genes for the corresponding species based on ENSEMBL annotations (STRING v12.0 annotations for Gorilla gorilla). For each species and gene set, the top 20 enriched pathways were selected based on FDR and ranked by fold enrichment. For these pathways, the number of overlapping genes (nGenes), total pathway size, and fold enrichment were extracted. An additional metric, termed enrichment quantity, was computed by combining nGenes, pathway size, and fold enrichment as follows:

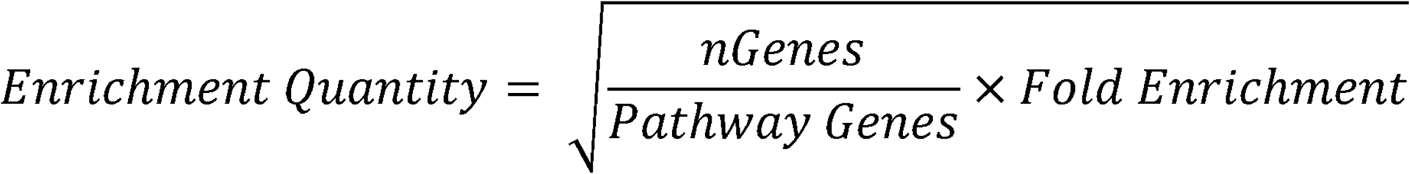

Custom Python scripts (pandas) (pathway_enrichment_data_manip.py, res_analyzer_with_common_gene_counts.py; see **Supplementary data-scripts**) were used to restructure enrichment outputs for downstream analysis, and R (plot_code_pathway_enrichment.R, see **Supplementary data-scripts**) was used to generate heatmaps summarising pathway enrichment patterns across gene sets and species.

## 3. Results

### 3.1. SCFRs Are Short, Abundant, and Evenly Distributed Across Reading Frames

To identify stop-codon-free regions (SCFRs) in primates, we systematically scanned all six reading frames of high-quality telomere-to-telomere assemblies for human, chimpanzee, gorilla, bonobo, gibbon, Bornean orangutan, and Sumatran orangutan **(Supplementary Table 1).** SCFRs were defined as uninterrupted stretches of sequence lacking in-frame stop codons (TAA, TAG, or TGA) in a given reading frame. Across species, we identified approximately 300 million SCFRs per genome, with totals ranging from 307.7 million in the Bornean orangutan to 342.5 million in the gorilla. Most SCFRs were extremely short, with median lengths consistently ∼39 bp (first quartile 15-18 bp and third quartile 75-78 bp) and mean lengths narrowly distributed between ∼56-59 bp, reflecting a shared primate-wide landscape dominated by compact ORF-like segments **(Figure 1a, Supplementary Tables 2 and 3)**. The shortest detectable SCFRs were three bp for all species, representing a single codon lacking a stop signal and constituting the minimal theoretical proto-ORF. Although the vast majority of SCFRs were compact, maximum lengths varied substantially among species, with most genomes containing SCFRs of up to ∼37-58 kb. Interestingly, the gorilla assembly contained notable length outliers, including an exceptionally large SCFR exceeding 808 kb, which contributed to its elevated length variance (standard deviation 123 bp).

**Figure 1:**
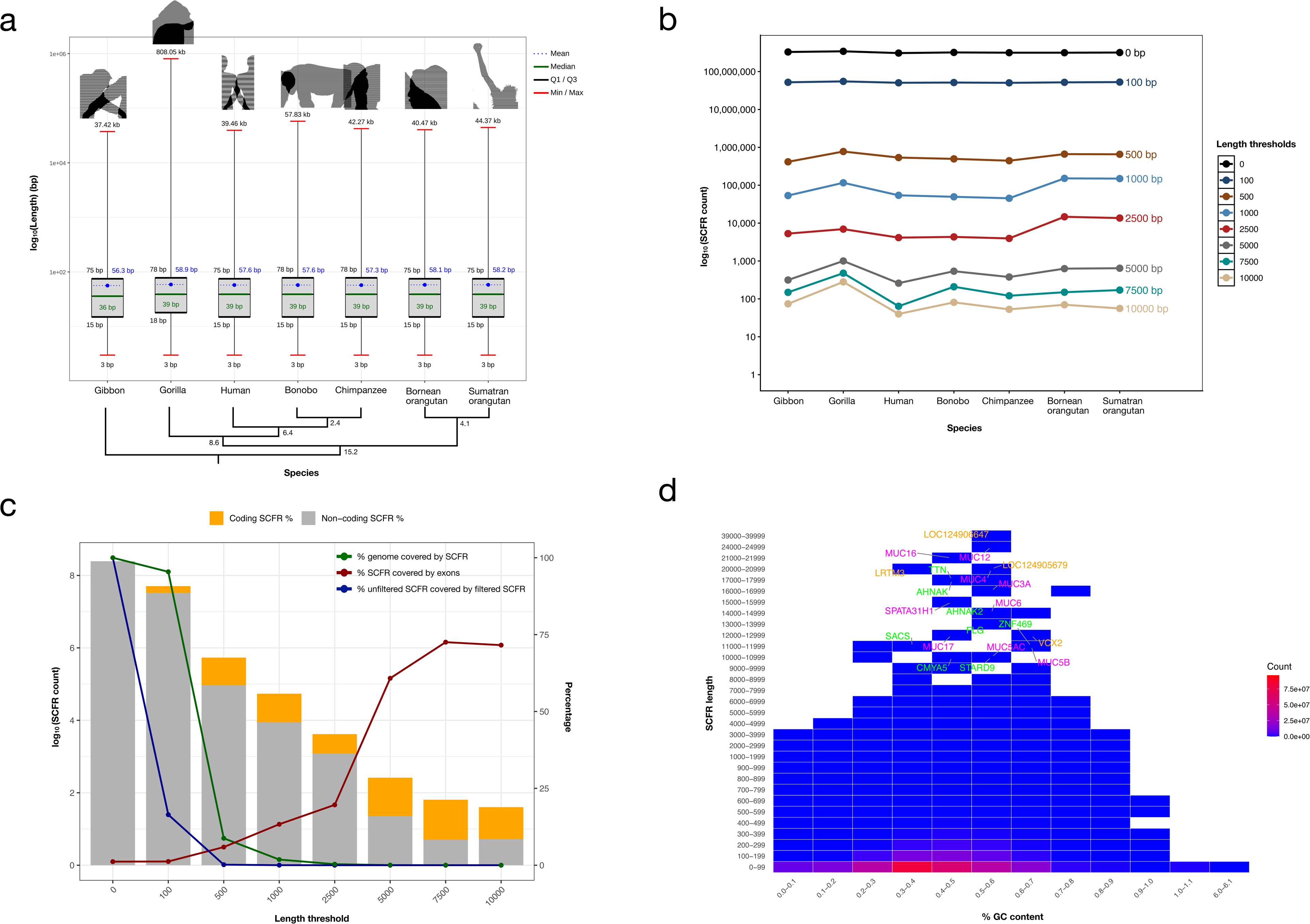
Length, Abundance, and Compositional Landscape of SCFRs Across Primate Genomes. **(a) Distribution of Stop-Codon-Free Region (SCFR) lengths across seven primate genomes:** Box plots summarise the genome-wide length distribution of SCFRs identified in the high-quality genome assembly of seven primate species. For each species, the inner box indicates the interquartile range (Q1-Q3), with the median represented by the central horizontal line. The blue point marker denotes the mean SCFR length, while whisker-like vertical lines extend to the minimum and maximum observed lengths. Lengths are shown on a log_10_ scale (bp), highlighting a predominant abundance of short SCFRs alongside rare, long SCFRs that span tens to hundreds of kilobases. Species included: bonobo, chimpanzee, gorilla, human, gibbon, Sumatran orangutan, and Bornean orangutan. **(b) Total SCFR count at different minimum-length thresholds:** The plot shows the genome-wide abundance of SCFRs in seven primate species across increasing minimum length thresholds (0 bp, 100 bp, 500 bp, 1 kb, 2.5 kb, 5 kb, 7.5 kb, and 10 kb). The X-axis lists the species compared, and the Y-axis displays SCFR counts on a log_10_ scale to accommodate the wide dynamic range in values. Each line represents a distinct minimum-length cutoff, illustrating a steep decline in SCFR abundance as length requirements increase, highlighting that long proto-gene-like ORFs constitute only a small subset of all potential SCFRs across primate genomes. **(c) Quantitative and genomic distribution properties of SCFRs in the human genome:** The plot summarises how SCFRs scale with increasing minimum length thresholds in the human genome. The X-axis represents minimum SCFR length cutoffs (bp), while the left Y-axis displays the total number of SCFRs on a log_10_ scale, and the right Y-axis shows percentage covered. The bars depict the SCFR count at each threshold, with stacked segments indicating the proportions intersecting coding exons (orange) and non-coding genomic regions (grey). The lines represent percentage covered in terms of length: green denotes the percentage of the genome covered by SCFRs, red indicates the percentage of SCFR bases overlapping coding exons, and blue shows the percentage of unfiltered SCFRs (0-bp threshold) retained after applying each length filter. Collectively, these data highlight that short SCFRs are highly prevalent and span almost the entire genome, whereas long SCFRs rapidly diminish in abundance. In contrast, the proportion of SCFR bases overlapping annotated coding exons steadily increased with higher length thresholds, indicating that longer SCFRs are more likely to contain coding regions. **(d) Genome-wide distribution of human SCFRs by length and GC content:** This heatmap displays the total number of SCFRs in the human genome across two dimensions: SCFR sequence length (y-axis; binned in 100 bp increments below 1 kb and 1 kb increments above 1 kb) and GC content (x-axis; binned in 0.1 fractional units). The colour scale represents SCFR frequency per bin, with warmer colours indicating higher abundance. Gene names labelled adjacent to long-SCFR bins denote protein-coding genes that intersect with SCFRs longer than 10 kb. The colours green, magenta, and orange indicate genes that appear in all species, at least in two species, and in a unique species, respectively.

We next examined whether SCFRs show preferential accumulation on either DNA strand by calculating strand asymmetry based on total SCFR count and length across forward and reverse strands. Strand asymmetry values range from -1 to +1, where 0 reflects a complete symmetry with balanced distribution, and positive or negative values indicate bias toward the forward or reverse strand, respectively. Across all primate genomes analysed, strand bias was negligible, with asymmetry values between -0.06 and +0.08 for SCFR counts and -0.01 to +0.01 for SCFR lengths **(Supplementary Figure 1, Supplementary Table. 4)**. Only a few unplaced scaffolds in bonobo showed moderate SCFR count asymmetry (−0.4 in NW_026946709.1, NW_026946710.1, NW_026946713.1; +0.4 in NW_026946711.1, NW_026946712.1), though this did not extend to SCFR lengths. This localised asymmetry is likely attributable to assembly or sequencing artefacts, as these scaffolds are relatively short, lack chromosomal anchoring, and therefore may contain unresolved strand annotation errors. The overall lack of strand preferential enrichment suggests that SCFRs arise largely through strand-neutral mutational and evolutionary processes rather than through directional selection pressures.

### 3.2. Length-Dependent Reduction in SCFR Frequency and Genomic Span

To evaluate how length thresholds affect SCFR abundance, genome coverage, and coding enrichment potential, we quantified SCFRs exceeding 100 bp, 500 bp, 1 kb, 2.5 kb, 5 kb, 7.5 kb, and 10 kb across seven primate genomes. While unfiltered SCFRs were widespread and spanned nearly the whole genome (99-100%), counts dropped dramatically with length, from ∼308–342 million unfiltered regions to ∼0.42–0.78 million at ≥500 bp and <300 at ≥10 kb, causing genome coverage to fall below 1% at thresholds ≥2.5 kb (**Figure 1b-c, Supplementary Figure 3, Supplementary Table 5**). Notably, >99.7% of all SCFRs identified were shorter than 500 bp in every species, underscoring how rare long uninterrupted ORF-like regions are in primate genomes. Despite the overarching trend, we detected species-specific differences at longer thresholds. Gorilla contained a disproportionately higher number of long SCFRs at ≥5 kb (CV = 47.2%), ≥7.5 kb (CV = 69.8%), and ≥10 kb (CV = 89.6%) relative to other primates (**Figure 1b, Supplementary Figure 2**). A similar enrichment was observed in Bornean and Sumatran orangutans at 1 kb (CV = 55.1%) and 2.5 kb (CV = 61.1%).

### 3.3. Longer SCFRs Tend to Contain a Higher Proportion of Coding Exons

A contrasting pattern emerged when evaluating the overlap between SCFRs and annotated coding exons. As the minimum length threshold increased, the proportion of SCFR bases intersecting coding regions rose markedly from ∼1% in unfiltered and ≥100 bp SCFRs to 16.8–60.7% at ≥5 kb, with a pronounced spike between the 2.5 kb and 5 kb thresholds (**Figure 1c, Supplementary Figure 3, Supplementary Table 5**). Beyond 5 kb, this proportion continued to increase in human, chimpanzee, and both orangutan species, plateaued in bonobo and gibbon, and decreased in gorilla. The overall trend suggests that long, uninterrupted ORF-like regions are more likely to fall within functional genes rather than non-coding DNA, indicating the underlying selection constraint. In contrast, short SCFRs are abundant in non-coding regions, where stop codons are frequently found.

### 3.4. GC-Dependent Constraints on SCFR Persistence

To evaluate the compositional biases that shape SCFR persistence, we quantified the percentage of GC and AT content for all SCFRs and examined their two-dimensional distributions across length and base-composition bins. Both the GC and AT composition profiles revealed the same fundamental trend: the majority of SCFRs clustered at short lengths (<1 kb), producing a dominant unimodal distribution characterised by high-density (warmer) bins (**Figure 1d, Supplementary Figure 4**). Shorter SCFRs additionally extended into both GC-poor/AT-rich and GC-rich/AT-poor regions, whereas long SCFRs (>10 kb) were rare and primarily confined to moderate GC/AT bins, indicating greater compositional constraints on extended uninterrupted ORFs. Despite the overall unimodal pattern, the distributions differ in the direction of skew: GC content shows a strong right skew, with a peak at 0.3-0.4, whereas AT content is left skewed, with a corresponding peak at 0.6-0.7. Notably, the gorilla deviated from the otherwise unimodal pattern, displaying a secondary peak in low-GC/high-AT bins (0.1-0.2 GC; 0.8-0.9 AT), suggesting a lineage-specific enrichment of GC-depleted genomic contexts that are permissive to SCFR formation (**Supplementary Figure 5**).

Long SCFRs are consistently localised near genes of exceptionally large or repetitive structure, including *TTN, AHNAK, AHNAK2, ZNF469, FLG, SACS, CMYA5, and STARD9*, which were shared across all examined species (**Figure 1d**). Members of the mucin (*MUC*) gene family were also prominently enriched, with at least seven MUC genes represented across species, reflecting their highly repetitive and expansion-prone genomic environments. We also detected species-specific gene associations; however, most corresponded to uncharacterised loci, indicating that long SCFRs may serve as reservoirs for discovering and annotating novel or previously unrecognised genes.

### 3.5. SCFR Localisation and Gene-like Signatures in Primate Gene Deserts

#### 3.5.1. Distribution of SCFRs Across Gene Deserts in Seven Primate Genomes

To determine whether gene deserts, defined as large intergenic regions devoid of annotated coding genes, harbour gene-like elements such as proto-genes, we examined the distribution of SCFRs within these regions across all seven primate genomes. To do this, we first identified gene deserts in the genomes of all seven primates using a Z-score-based approach applied to intergenic regions. Intergenic segments were broadly comparable among species, totalling ∼21,000 elements (ranging from 19,680 in human to 21,730 in gorilla) with a mean length of ∼99 kb (93.8 kb in bonobo to 104.4 kb in human) (**Figure 2a, Supplementary Figure 6a, Supplementary Table 6**). From these, we identified gene deserts, which varied from 331 in the Bornean orangutan to 528 in the chimpanzee, with mean lengths of ∼2.1 Mb (1.85 Mb in bonobo to 2.4 Mb in gorilla) (**Supplementary Figure 6b**). When intersecting SCFRs with gene deserts, we found that all SCFRs ≥5 kb in length were located entirely within gene deserts (**Figure 2b, Supplementary Figure 7, Supplementary Table 7**). In contrast, SCFRs shorter than 2.5 kb frequently exhibited only minimal overlap with gene deserts, sometimes as little as ∼3 bp. These patterns indicate that longer SCFRs preferentially reside within deep intergenic regions, whereas shorter elements tend to be positioned near gene-desert boundaries.

**Figure 2:**
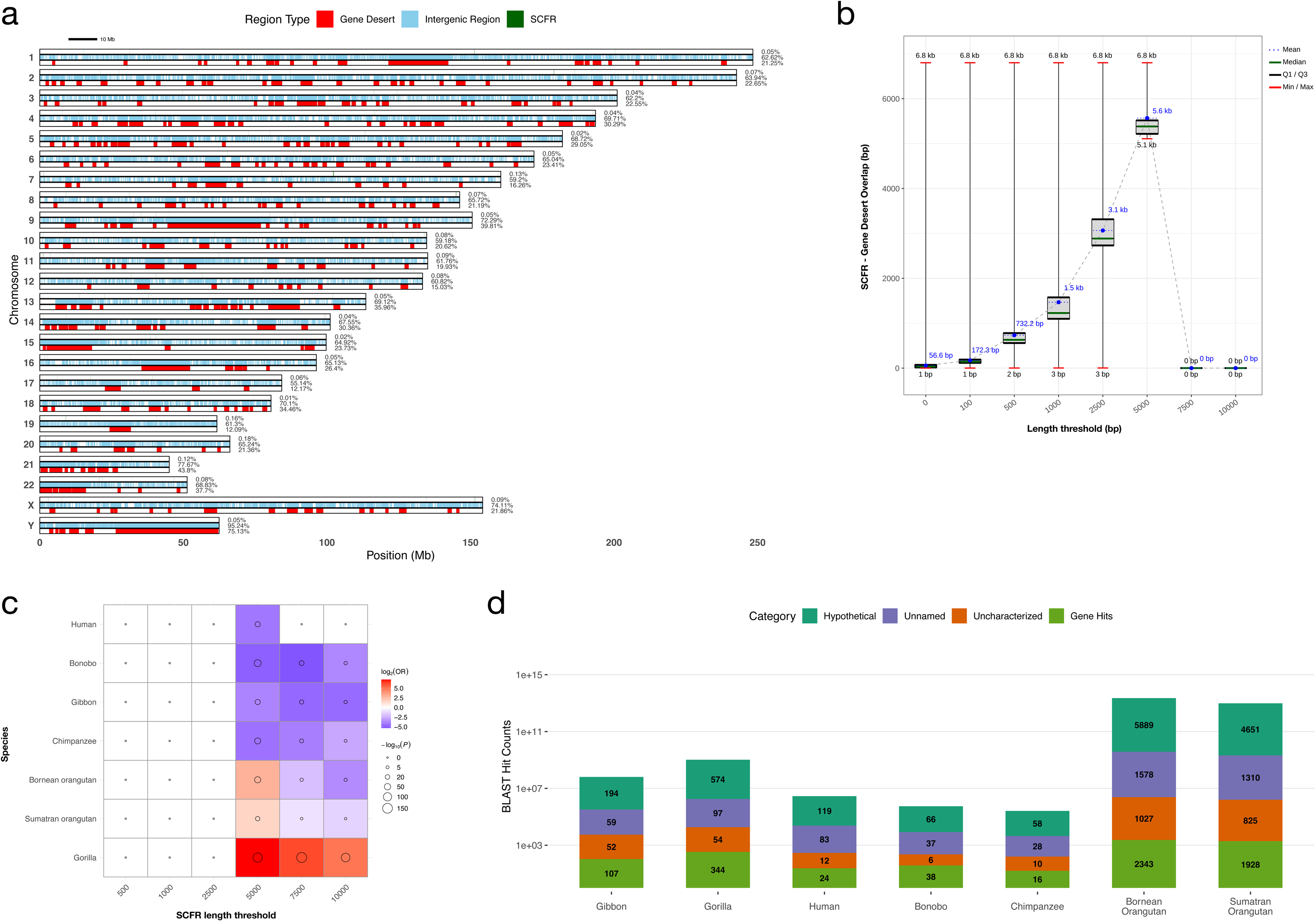
SCFR–Gene Desert Relationships: Distribution, Overlap, and Enrichment Across Primates. **(a) Distribution of Gene Deserts, Intergenic Regions, and SCFRs Across the Human Genome:** This figure maps the distribution of three genomic features, including Gene Deserts (GD, red), Intergenic Regions (IR, light blue), and SCFRs ≥5 kb (green) across the human chromosomes (Y-axis), visualised by genomic position in megabases (Mb, X-axis). The three parallel bars for each chromosome illustrate the extent and location of each feature type. At the end of each bar, the percentage of the genome covered by the features is labelled. Across the entire genome, IR constitute 65.91% of the space, GD constitute 25.56%, and the SCFRs (≥5 kb) constitute a small fraction at 0.06%. **(b) Length-Dependent Overlap Between SCFRs and Gene Deserts in the Human Genome:** This figure shows how the genomic overlap between SCFRs and gene deserts varies across increasing SCFR length thresholds. The x-axis represents minimum SCFR lengths, and the y-axis reports their overlap with gene deserts in base pairs, with the minimum, maximum, median, quartiles, and mean overlap indicated for each threshold. SCFRs ≥5 kb consistently show complete containment within gene deserts, with all statistics converging at full overlap values. In contrast, SCFRs at thresholds of 2.5 kb and below display minimal or negligible overlap, with ranges extending from 0 to only three bp. **(c) Enrichment and Depletion Patterns of SCFRs Within Gene Deserts Across Primate Genomes:** This figure illustrates the enrichment or depletion of SCFRs within gene deserts across primate species using Fisher’s exact tests at multiple SCFR length thresholds. Each point represents a species-threshold combination, with colour indicating the log_2_-transformed odds ratio (log_2_(OR)): red denotes enrichment (OR > 1) and blue denotes depletion (OR < 1). Point size reflects statistical significance, shown as -log_10_(P), where larger points correspond to more significant associations. A capped value is applied to the infinite OR observed for gorilla at the 5,000 bp threshold, plotted as one unit above the maximum log (OR) to visualise it as the strongest enrichment (deepest red). Together, the plot highlights marked species-specific and length-dependent patterns, with some primates showing depletion of large SCFRs in gene deserts and others, most notably gorilla, exhibiting strong enrichment. **(d) Distribution of BLASTP hits across species:** Stacked bar plot showing the composition of BLASTP hits identified from gene desert-derived ORF sequences in each species. The x-axis denotes species, and the y-axis represents log-transformed total hit counts. Stacked colours indicate annotation categories: hypothetical (green), unnamed (blue), uncharacterized (orange), and annotated gene hits (light green).

#### 3.5.2. Length-Dependent Enrichment of SCFRs Within Primate Gene Deserts

To assess whether SCFRs preferentially occur within gene-desert regions, we performed Fisher’s exact tests comparing SCFRs at multiple minimum length thresholds against gene deserts. For shorter SCFRs (500 bp, 1 kb, and 2.5 kb), the test yielded non-significant results (right p-value=1, Odds Ratio (OR) = NaN), indicating a random or ubiquitous distribution that precludes a meaningful assessment of enrichment or depletion (**Figure 2c, Supplementary Table 8**). In contrast, for the longer SCFRs (5 kb, 7.5 kb, and 10 kb), the test returned lower two-tailed p-values (P<10^-5^ in nearly all cases), indicating a non-random association. While human, chimpanzee, bonobo, and gibbon all showed strong depletion of large SCFRs within gene deserts (OR < 1), the gorilla genome stood out as a major exception, displaying robust enrichment across all longer thresholds. Specifically, gorilla 7.5 kb SCFRs showed an OR of 74.9 (P=8.14×10^-158^), and the 5 kb threshold resulted in an infinitely large Odds Ratio, pointing to a co-localisation. The orangutan species also exhibited significant enrichment, although only for the 5 kb threshold (OR = 7.357 for Bornean orangutan and OR = 3.043 for Sumatran orangutan), with longer SCFRs in these species showing depletion. The significant, length-dependent enrichment of longer SCFRs within gene deserts in certain primate species challenges the traditional view of these intervals as entirely non-coding and suggests their potential role as evolutionary grounds for *de novo* gene birth.

#### 3.5.3. Homology and Coding-Signature Analyses Reveal Proto-Gene Potential in SCFRs

To identify potential proto-genes within SCFRs, we extracted unique ORFs of a minimum of 600bp based on a 70% identity threshold in CD-hit. This yielded 18 ORFs in bonobo, up to 1324 ORFs in Bornean orangutan. A homology search using BLASTP (e-value = 1e-4, query coverage = 70%) with these ORFs as query against the nr database detected several proteins with significant e-values (**Supplementary Table 9**). Among the seven primates examined, the Bornean orangutan showed the highest number of total BLASTP hits (10,837), followed by the Sumatran orangutan (8,714) and the gorilla (1,069), whereas the chimpanzee (112) and the bonobo (147) exhibited comparatively fewer matches. Intermediate values were observed in the gibbon (412) and the human (238) (**Figure 2d, Supplementary data-nr_blastp**). While the majority of top hits were unnamed proteins, or hypothetical proteins, several of the hits consisted of predicted proteins (XP accessions ranging from 18.5% in the chimpanzee to 31.2% in the Sumatran orangutan. Analysis of the specific predicted protein hits revealed recurrent homology to a limited set of protein families, many of which are known to be highly repetitive or rapidly evolving (**Supplementary Table 10**). Among these, some genes, such as the mucin-like and proline-rich families, showed homology to ORFs across all seven primate species, highlighting their recurrent or ancient origin, with the most abundant hits being mucin-2-like (17 total hits) and mucin-17-like (13 total hits). Other hits were shared between at least two primate species, including proteins like Keratin-associated protein-like, Plectin-like(11 total hits), Dynein heavy chain-like (7 total hits), Serine/arginine repetitive matrix protein 2-like, and Fap1 adhesin-like (6 total hits). Notable species-specific homology was also observed, such as the Endochitinase 2-like gene, which gave hits to 19 ORFs exclusively from the Sumatran orangutan genome, demonstrating potential lineage-specific proto-gene candidates. Collectively, these results indicate that long SCFR-derived ORFs in gene deserts frequently exhibit detectable similarity to repetitive, low-complexity, or structurally biased protein families, rather than to well-characterised conserved genes.

In addition to sequence homology, Discrete Fourier Analysis (DFA) was performed on these ORFs to detect intrinsic coding signatures. DFA showed distinct peaks at a frequency of 0.33 only in the Bornean and Sumatran orangutan ORFs (two canonical and nine non-canonical ORFs), corresponding to the expected codon triplet periodicity found in bona fide coding sequences (**Supplementary Table 11, 12**). Taken together, these results suggest that gene deserts contain numerous ORFs with homology to existing proteins, with some showing codon triplet-associated periodicity, indicating the potential for these regions to play a role as reservoirs for the emergence of proto-genes.

### 3.6. Exon Shadows and Exitrons Define Structured, Coding-like Landscapes at Exon Boundaries

#### 3.6.1. Exon Shadows Capture Boundary Offsets Between Exons and SCFRs

To investigate how existing genes accommodate SCFRs, we defined exon shadows as the distance between exon boundaries and SCFR boundaries. For each SCFR-exon pair, an upstream exon shadow was defined as the distance between SCFR and the exon start (strand-aware), while a downstream exon shadow was defined as the distance between the exon and SCFR end (**Figure 3a**). A shadow length of zero indicates exact coincidence between exon and SCFR boundaries. The distribution of exon shadow lengths showed that the majority of shadows are short across all species (**Figure 3b**). Upstream shadows were consistently longer than downstream shadows, with higher mean (∼87 bp) and median (∼55 bp) lengths compared with downstream shadows (mean ∼71 bp; median ∼41 bp). The minimum shadow length was 1 bp for both upstream and downstream shadows in all species. In contrast, maximum shadow lengths varied widely among species, ranging from ∼11.8 kb in human to ∼56.1 kb in bonobo for upstream shadows, and from ∼16.8 kb in gibbon to ∼37.6 kb in bonobo for downstream shadows.

**Figure 3:**
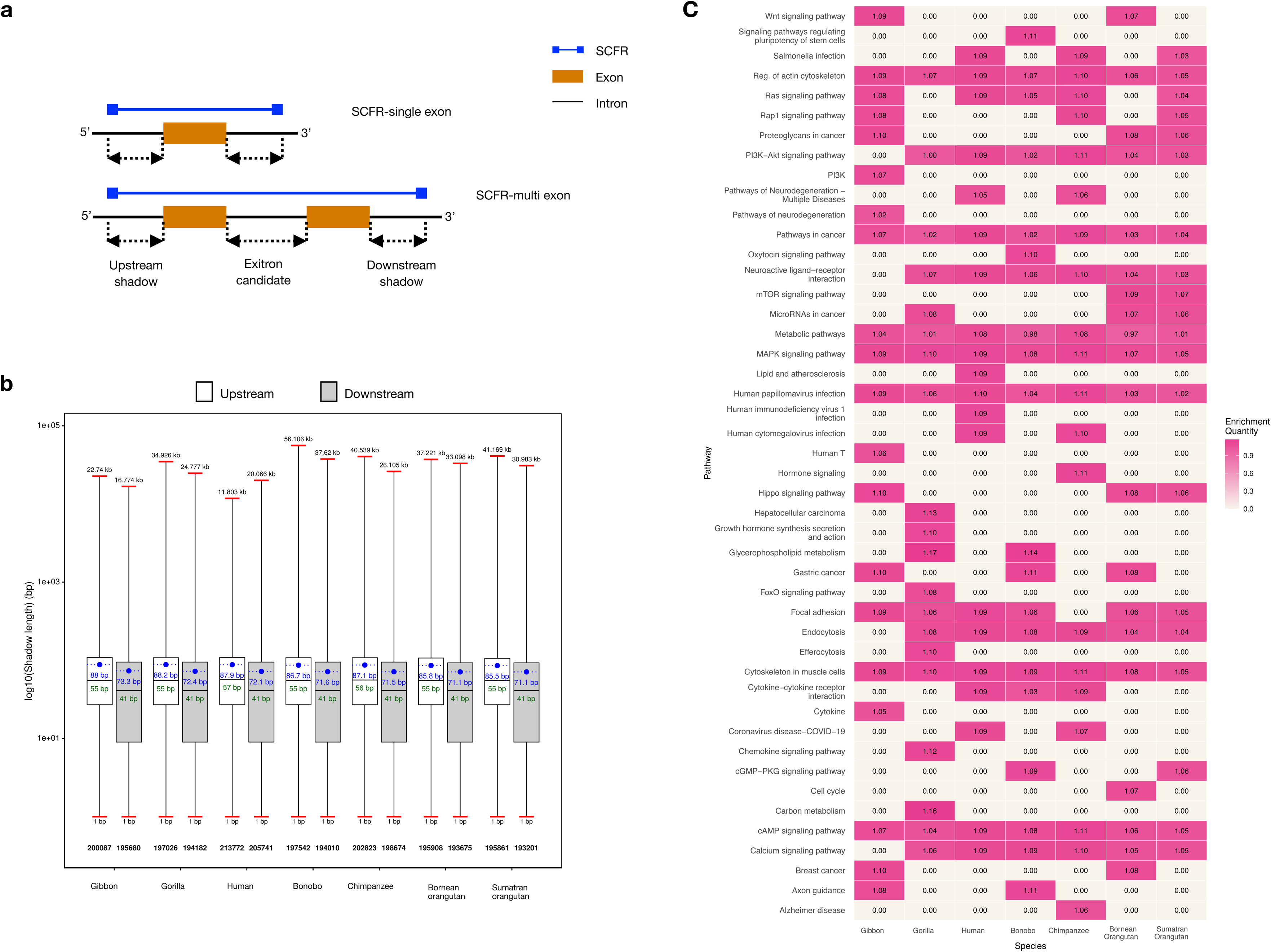
Exon Shadows: Structure, Length Distribution, and Functional Enrichment. **(a) Schematic representation of exon shadows and exitrons:** Diagram illustrating the definition of exon shadows and exitron candidates within exon–SCFR configurations. For a single exon–SCFR pair, an upstream exon shadow is defined (strand-aware) as the in-frame distance from the SCFR start to the annotated exon boundary, whereas a downstream exon shadow is defined as the in-frame distance from the exon boundary to the SCFR end. In multi-exon-SCFR configurations, introns that are completely spanned by a continuous SCFR extending across two adjacent exons are classified as exitron candidates. **(b) Length distribution of upstream and downstream exon shadows across primates:** Box plots summarising the log -transformed length distributions of upstream and downstream exon shadows in seven primate species. For each species, boxes represent the interquartile range (Q1-Q3), with the median shown as a black horizontal line. Blue points denote mean shadow lengths, and whiskers extend to the minimum and maximum observed values. Although the y-axis is log-transformed to facilitate comparison, axis tick labels indicate the corresponding untransformed lengths. **(c) Pathway enrichment of genes containing exon shadows across primates:** Heatmap showing biological pathways enriched among genes with exon shadows in seven primate species, using the default genomic background for each species. Species are displayed on the x-axis and enriched pathways on the y-axis. Colour intensity ranges from off-white (no significant enrichment) to progressively darker pink shades, indicating higher enrichment levels.

Across all species, ∼95% of genes contained at least one exon-SCFR pair with a non-zero shadow (**Supplementary Figure 8a**). This proportion decreased to ∼51% at the transcript level and ∼19% at the exon level. Thus, while SCFRs are widespread within genes, their association with individual transcripts and specific exon boundaries is more restricted. The majority of exon shadows were positive (∼88%), indicating that SCFRs typically extend beyond exon boundaries (**Supplementary Figure 8b**). Zero-length shadows accounted for ∼11% of cases, reflecting exact alignment between exon and SCFR boundaries. Negative shadow lengths were extremely rare (∼0.02%). These cases likely arise from splice-site geometry rather than true exon extension, occurring when an SCFR boundary defined by an amber stop codon (TAG) overlaps a splice acceptor (AG) and the exon has a non-zero phase, causing the exon to extend slightly beyond the SCFR boundary. (For the complete table of exon shadows identified, refer to **Supplementary data**.)

#### 3.6.2. Characterisation of Exon Shadows Across Transcript Structures

The vast majority of SCFR-transcript pairs with exon shadows contained a single exon (∼99%), whereas only ∼1% involved multiple exons, with a maximum of seven exons in bonobo and 24 in human transcripts (**Supplementary Figure 9a**). Neither the number nor the length of exon shadows showed any detectable strand bias, and no systematic asymmetry was observed between upstream and downstream shadows (**Supplementary Figure 10, Supplementary Table 13**). Together, these results indicate that exon shadows are distributed uniformly with respect to both transcriptional strand and relative exon position within SCFRs. Analysis of exon position revealed that first and last exons harboured comparable amounts of upstream shadow, whereas last exons lacked downstream shadows, consistent with termination at annotated stop codons (**Supplementary Figure 9b**). Examination of exon splicing categories showed that shadows predominantly occur in constitutively spliced exons across species (62-69%), with a smaller fraction associated with alternatively spliced exons (30-37%). Human transcripts represented a notable exception, where alternatively spliced exons slightly exceeded constitutive ones (53.7% vs 43.3%; **Supplementary Figure 9c**). Overall, these patterns suggest that exon shadows preferentially associate with exon architecture, while allowing species-specific modulation through alternative splicing.

#### 3.6.3. Enrichment and Codon-Structured Organisation of Exon Shadow Lengths Across Species

Shadow-length enrichment analysis revealed consistent over-representation of 21 to 30 bp upstream shadows and of 1-10 bp and 21-30 bp downstream shadows across species (**Supplementary Figure 11**). Among these, 1 bp downstream shadows showed particularly strong enrichment (∼32,043 occurrences; enrichment ratio ∼81.7) (**Supplementary Figure 12**). Beyond the shortest lengths, shadow counts exhibited a clear ∼3 bp periodicity in both upstream and downstream directions, consistent with codon structure and reading-frame constraints (**Supplementary Figure 13**). In addition, shadow frequencies declined progressively with increasing length, reflecting the reduced likelihood of maintaining long in-frame regions without encountering a stop codon. Together, these patterns indicate that exon shadow length distributions are structured and reproducible across species.

#### 3.6.4. Shadow-Dependent Differences in Functional Pathway Enrichment

Gene set enrichment analysis across five shadow-defined gene sets revealed clear differences in the extent and consistency of pathway enrichment. Genes with shadows (GWS) showed the strongest signal, with multiple pathways enriched across all seven species, whereas genes without shadows (GWOS) lacked any pathway consistently enriched across species (**Figure 1c**, **Supplementary Figure 14**). Genes with symmetric shadows (GWSS) showed the fewest enriched pathways, reflecting their limited representation. Several pathways were recurrently enriched across shadow-containing datasets. Cytoskeleton-related pathways were enriched in all datasets except GWOS. MAPK signalling, cancer-related pathways, and focal adhesion were enriched primarily in datasets retaining downstream shadows (GWOD and GWS) (**Supplementary Figure 15**). In contrast, pathways related to protein processing in the endoplasmic reticulum and nucleocytoplasmic transport were enriched only in genes lacking upstream shadows (GWOU). Although species-specific enrichments were widespread, the overall trends were conserved, with gorilla showing the highest number of enriched pathways and humans displaying comparatively fewer enrichments.

#### 3.6.5. Identification and Length Distribution of Exitrons Across Species

Exitrons are exon-like intronic regions whose retention preserves the protein-coding potential of the transcript. To characterise exitron architecture, we analysed the length and sequence properties of exitron candidates, defined as introns fully spanned by an SCFR that extends across two adjacent exons, thereby generating an intronic segment devoid of in-frame stop codons. Across species, we identified approximately 2,340 exitron candidates per genome. Exitron lengths were moderately conserved, with a mean of ∼155 bp and a median of ∼110 bp (**Figure 4a, b**). Minimum exitron lengths were comparable across species (27-39 bp), whereas maximum lengths varied substantially, ranging from ∼3.24 kb in human to ∼9.8 kb in bonobo. This broad but asymmetric length distribution indicates that while lower bounds of exitron size are constrained, upper bounds are more permissive and lineage-specific, consistent with variable tolerance for long coding-compatible intronic segments.

**Figure 4:**
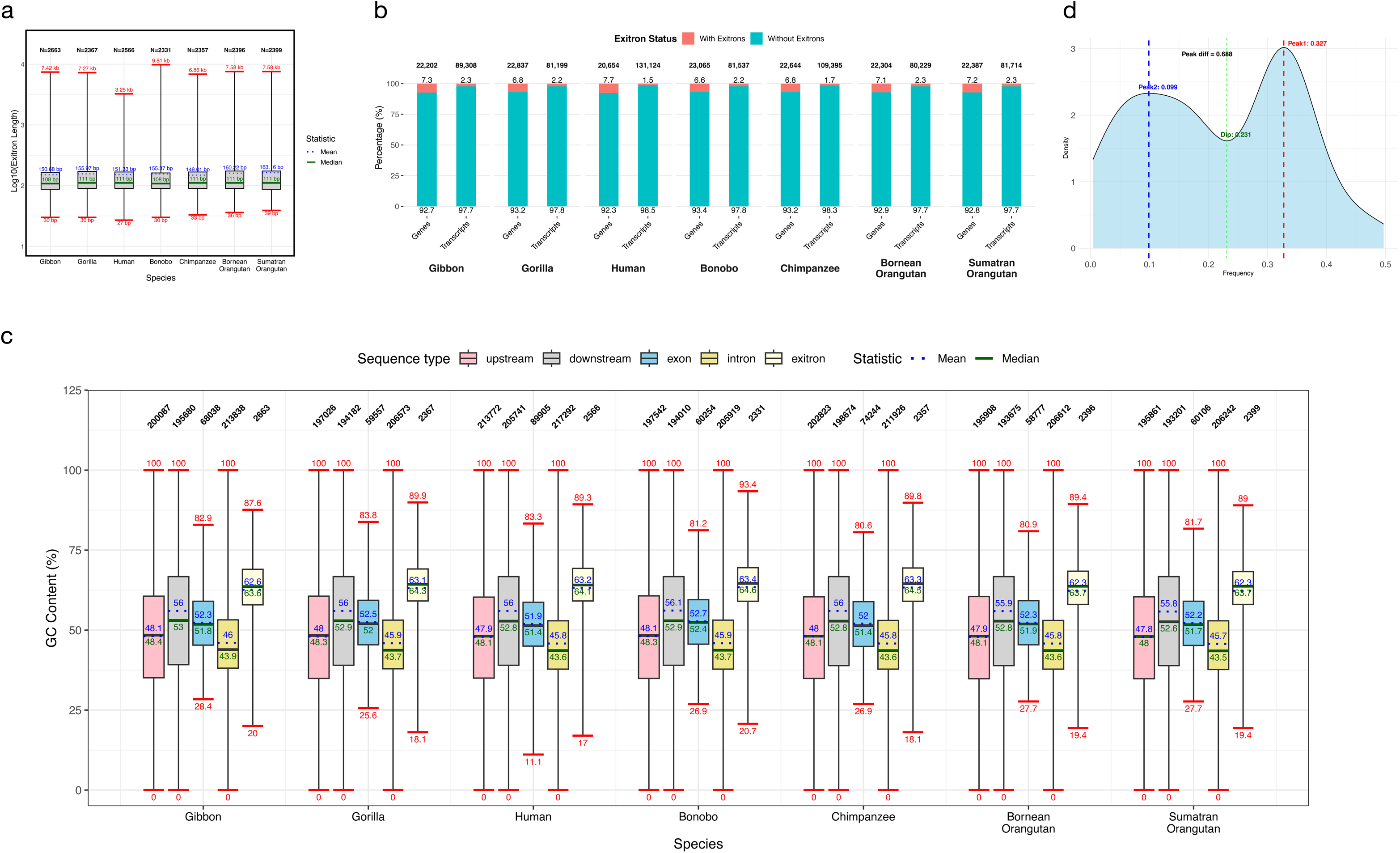
Comparative Genomic Analysis of Exitron Candidates in Primates. **(a) Length distribution of exitron candidates across primates:** Box plots showing the log -transformed length distributions of exitron candidates in seven primate species. For each species, boxes indicate the interquartile range (Q1-Q3), with the median represented by a black horizontal line. Blue dots mark the mean length, and whiskers extend to the minimum and maximum observed values. Although the y-axis is log-transformed to improve visualisation, tick labels correspond to the original (untransformed) lengths. **(b) Percentage of genes and transcripts with and without exitrons:** The x-axis is arranged in two tiers, with the lower tier showing species and the upper tier showing genomic regions (genes and transcripts). The y-axis indicates the percentage of the region with and without exitrons. The red stacks represent regions with exitrons, while the teal stacks represent regions without exitrons. The percentages are annotated above and below the respective bars. The total number of genes and transcripts for the respective species is annotated above the respective bars. **(c) GC Content Distribution Across Sequence Types in Primates:** The x-axis indicates species, and the y-axis indicates GC content (%). For each species, boxplots show the GC% distribution for five sequence types: upstream shadows (pink), downstream shadows (light grey), exons (blue), introns (yellow), and exitrons (cream). Whisker caps, indicated in red, mark the minimum and maximum values. Solid green and dotted blue lines denote the median and mean, respectively. Maximum and mean GC% values are annotated above the corresponding lines, and minimum and mean values below. Total sequence counts for each boxplot are indicated above the boxes. **(d) Fourier Frequency Distribution of Human Exitron Candidates:** Kernel density estimates (KDEs) of spectral frequency distributions derived from discrete Fourier transform (DFT) analysis of upstream exon shadows in human. The x-axis represents spectral frequency, and the y-axis denotes density. Red and blue dotted vertical lines indicate the two dominant spectral peaks, while the green vertical line marks the minimum (trough) between them. The annotated peak difference corresponds to the distance between the lowest and highest peak frequencies.

#### 3.6.6. Pathway Enrichment Highlights Shared and Divergent Roles of Exitrons

To assess the functional context of exitron-containing genes, we performed gene set enrichment analysis separately for each species. The extracellular matrix (ECM) pathway was consistently enriched across all seven species, indicating a shared functional association of exitrons with genes involved in structural and signalling interactions (**Supplementary Figure 15c**). Six additional pathways, including platelet activation, motor proteins, human papillomavirus infection, focal adhesion, cytoskeleton in muscle cells, and adrenergic signalling in cardiomyocytes, were enriched in all species except the gorilla. Alongside these conserved signals, distinct species-specific enrichments were observed. Neutrophil signalling and aldosterone synthesis and secretion were uniquely enriched in human and bonobo, respectively. Gibbons showed exclusive enrichment of Rap1 signalling, pancreatic secretion, and oxytocin signalling pathways, while Sumatran orangutans uniquely exhibited enrichment of insulin secretion and glutamatergic synapse pathways. Both orangutan species shared enrichment of AMPK signalling, MAPK signalling, endocrine resistance, insulin resistance, and glucagon signalling pathways. Notably, the gorilla exhibited the largest number of species-specific enrichments, with 17 pathways uniquely enriched. Together, these results indicate that while exitrons are associated with a conserved core of cellular and structural functions, their broader functional associations are shaped by lineage-specific regulatory and physiological contexts.

#### 3.6.7. Exon Shadows and Exitron Candidates Exhibit Conserved Coding-associated Nucleotide Signatures

Across all seven species, nucleotide composition showed highly conserved and feature-specific patterns that together define a graded transition from non-coding to coding sequence around exon–SCFR boundaries. Exitrons were consistently the most GC-rich and GC3-enriched elements, exceeding annotated exons and showing longer GC stretches but shorter AT stretches, consistent with strong coding-like constraints (**Figure 4c**, **Supplementary Figure 16-19**). Exons displayed intermediate values across all metrics, while introns were GC-poor, AT-rich, and compositionally symmetric. Exon shadows occupied an intermediate but asymmetric state: downstream shadows were markedly GC-enriched, positively GC-skewed, and depleted in AT stretches, approaching exitron-like composition, whereas upstream shadows were closer to introns, with lower GC content, mild negative GC skew, reduced GC3, and longer AT stretches. These directional differences indicate that compositional features change sharply and non-uniformly across exon boundaries, with downstream regions retaining stronger coding-associated signatures than upstream regions. Together, the concordant shifts in GC content, GC skew, GC3, and nucleotide stretch architecture argue that exon shadows are not compositionally neutral flanks but reflect boundary- and strand-dependent sequence constraints linked to exon structure and coding potential.

In all species, upstream and downstream shadows exhibit multiple conserved peaks (minimum 3). Upstream peaks occur at ∼0.33-0.34 and ∼0.02-0.03, with a dip at 0.2-0.3; density differences range from 0.455 (human) to 1.086 (Bornean orangutan) (**Supplementary Figure 20**). Downstream peaks are similarly conserved (∼0.33-0.34 and 0.034-0.053), with dips at 0.2-0.22 (except Bornean orangutan, 0.407) and peak differences of 0.843-3.145 (**Supplementary Figure 21**). Exitrons show at least two peaks, with a conserved ∼0.33 peak corresponding to codon triplet frequency, a secondary peak at 0.057-0.151, and a dip at ∼0.22-0.23; peak differences range from 0.062 (chimpanzee) to 2.986 (Sumatran orangutan) (**Supplementary Figure 22).** Together, these conserved spectral signatures, particularly the recurrent ∼0.33 codon-scale peak across shadows and exitrons, indicate that many SCFR-associated regions retain structured, frame-compatible periodic organisation rather than representing purely random sequence composition.

### 3.7. Evolutionary and Compositional Differentiation of SCFRs Across Length Scales in Primates

To investigate whether SCFRs exhibit gene-like compositional signatures, we quantified RSCU for SCFRs at increasing length thresholds (≥5 kb, ≥7.5 kb, and ≥10 kb), as longer SCFRs show a substantially higher proportion of coding overlap. For each threshold, PCA was performed using 61 sense codons (excluding the three stop codons), followed by k-means clustering. Clusters were annotated according to coding status (in-frame, out-of-frame, or non-coding), repeat content, and GC content to enable biological interpretation of compositional structure.

#### 3.7.1. Compositional Structuring of SCFRs Revealed by Codon Usage Analysis

Across thresholds, the proportion of variance explained by PC1 and PC2 generally increased with length in most species, most prominently in Gibbon, Human, and Chimpanzee, indicating more structured codon usage in longer SCFRs. Silhouette scores improved in parallel (peaking >0.4 in gibbon), and the Davies-Bouldin Index declined at higher thresholds, supporting enhanced cluster cohesion and separation (**Figure 5a**). The optimal number of clusters (k) remained stable (3 to 4) for most species, except in the chimpanzee at 10 kb (k=6).

**Figure 5:**
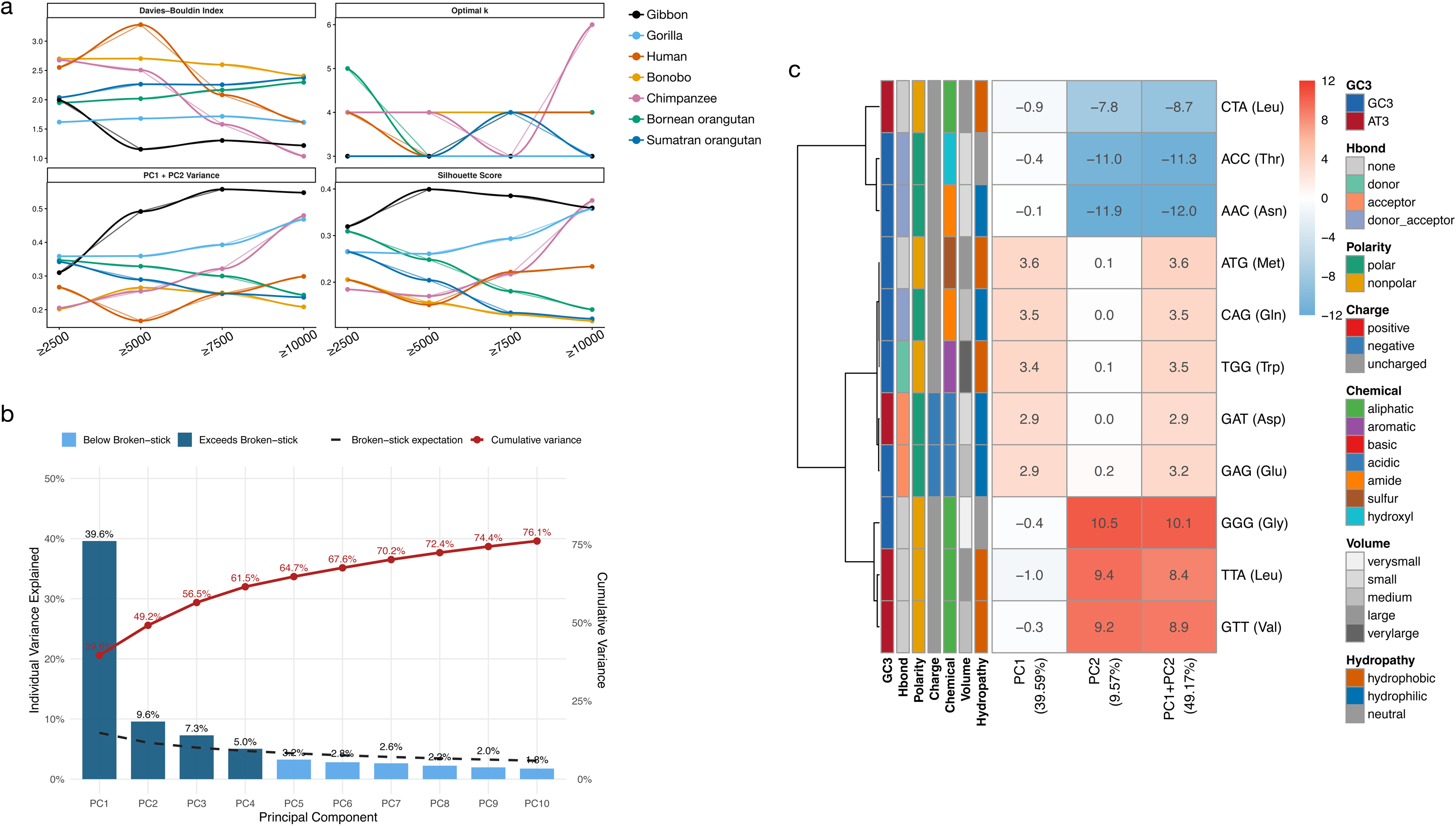
PCA-Based Characterisation of Codon Variation and Clustering Structure. **(a) Evaluation of PCA and Clustering Performance:** The four line plots assess the robustness and quality of principal component analysis (PCA) and clustering results of seven primates. Clustering performance is evaluated using the Davies-Bouldin Index (where lower values indicate better cluster separation) and Silhouette scores (where higher values reflect stronger within-cluster similarity and clearer separation between clusters). The optimal number of clusters (k) is determined based on clustering stability metrics. PCA performance is assessed by the proportion of variance explained by PC1 and PC2, indicating how effectively the first two principal components capture the overall variation in the dataset. Together, these metrics provide complementary measures of dimensionality reduction efficiency and clustering reliability. **(b) Variance explained by the Top 10 Principal Components in Gibbon:** The plots show the variance explained by the top 10 principal components (PCs) at a 5000bp length threshold. The X-axis denotes the PCs, while the Y-axis indicates the percentage of variance explained by each. Bars in dark blue exceed the expectations from the Broken-stick test, whereas light blue bars fall below it. The black dotted line represents the Broken-stick expectation for each PC, and the red line with circles shows the cumulative variance explained. The total number of retained PCs and the count of top 10 PCs exceeding the Broken-stick expectation are annotated above each plot. **(c) Relative Codon Contributions by Top 2 PCs in Gibbon:** The plots show the relative contribution of each codon to PC1, PC2, and the combined PC1+PC2 at a 5000bp length threshold. The X-axis represents the PCs and their explained variance, while the Y-axis lists codons with their corresponding amino acids. Adjacent bars indicate amino acid properties, including GC3 content, hydrogen bonding, polarity, charge, chemical class, volume, and hydropathy. Cell colours in the heatmap reflect contributions to variance, with cooler tones indicating negative contributions and warmer tones positive contributions.

Across all seven primates, SCFRs consistently resolved into two broad compositional regimes: (1) relatively homogeneous clusters with uniform GC content and strong enrichment of simple repeats, and (2) a large heterogeneous cluster characterised by broad GC variation and mixed repeat content (**Supplementary Figures 23-36**). Coding-overlapping SCFRs were rarely dispersed; instead, they were typically concentrated within a single heterogeneous cluster, whereas non-coding SCFRs were present in all clusters. At lower thresholds (≥2.5 kb), clusters were diffuse, GC variation was wide, and satellite as well as SINE repeats were more frequent. With increasing length thresholds, satellite and low-complexity repeats declined sharply across species, and SINE-associated substructures were progressively reduced, particularly beyond 5-7.5 kb.

GC bias, especially GC3, was the dominant axis of codon usage variation in most species (**Supplementary Figures 23-36**). PC1 frequently captured a GC-driven gradient separating low-GC, simple-repeat-rich SCFRs from higher-GC SCFRs enriched in coding overlap. In several genomes, GC segregation was evident within the heterogeneous cluster, with low-GC SCFRs forming compact subclusters. However, variance behaviour differed by lineage: variance explained by PC1 and PC2 increased with length in gibbon and chimpanzee, decreased in human and both orangutans, and showed non-linear dynamics in bonobo and gorilla (**Figure 5b**, **Supplementary Figures 37-40**). Notably, in human, cluster separability did not improve with length, and more principal components were required to explain comparable variance, indicating diffuse codon usage structure despite stringent length filtering. Together, these results indicate that increasing SCFR length reduces repeat-associated heterogeneity and enriches for coding-overlapping regions, but the degree of compositional consolidation is species dependent.

#### 3.7.2. Species-Specific Codon Usage Architecture and Repeat Associations

Despite a conserved homogeneous-heterogeneous cluster framework, marked lineage-specific differences were observed. In Gibbon, cluster architecture was stable from 5 kb onward, consistently resolving into two homogeneous clusters and one large heterogeneous cluster containing coding SCFRs (**Supplementary Figures 23, 24**). High-GC SCFRs were associated with SINE enrichment, whereas low-GC SCFRs were enriched in simple repeats. Codon loadings remained largely stable beyond 5 kb, indicating compositional robustness with increasing length (**Figure 5c, Supplementary Figures 41-44**). Gorilla exhibited a similar three-cluster configuration, but codon directionality shifted substantially across thresholds, transitioning from AT3-dominated signals at lower lengths to GC3-dominated signals at higher thresholds (**Supplementary Figures 25, 26**). Satellite repeats were abundant at 2.5 kb but absent by 10 kb. Coding SCFRs were confined to a heterogeneous cluster with persistent GC variation. Human displayed the most diffuse clustering. Even at higher thresholds, clusters remained heterogeneous with limited separability (**Supplementary Figures 27, 28**). Variance explained by the first two PCs decreased with increasing length, and at least six PCs were required to capture 50% of the total variance (**Supplementary Figure 38**). Low-GC SCFRs segregated within major heterogeneous clusters, but coding enrichment did not produce strongly cohesive groups. Simple repeats remained prevalent across all thresholds, whereas satellite and SINE repeats declined.

Among others, bonobo showed the strongest compositional structuring at 5 kb, where GC segregation along PC1 clearly separated homogeneous clusters from heterogeneous ones (**Supplementary Figures 29, 30**). Coding SCFRs tended to exhibit lower GC and were enriched in simple repeats. Beyond 7.5 kb, variance declined and repeat diversity sharply decreased.

Chimpanzee demonstrated improved separation at intermediate thresholds (5-7.5 kb), with low GC and simple repeats explaining cluster segregation; however, at 10 kb, cluster number increased despite reduced separability (**Supplementary Figures 31, 32**).

In the Bornean orangutan and Sumatran orangutan, overall separability was lower, and variance generally decreased with increasing length (**Supplementary Figures 33-36**). At intermediate thresholds, two homogeneous clusters (uniform GC and simple repeats) emerged alongside heterogeneous clusters containing mixed coding and non-coding SCFRs. Satellite-rich patches evident at 2.5 to 5 kb disappeared at higher thresholds. In the Sumatran orangutan, differences in top-loading codons further distinguished heterogeneous clusters at 7.5 kb despite limited overall cohesion. Collectively, these findings demonstrate that while GC bias, simple repeats, and reduction of satellite/SINE elements with increasing length represent conserved trends across primates, the extent of codon usage consolidation and cluster separability varies substantially among lineages. Progressive length filtering shifts SCFR composition away from repeat-dominated background structure toward coding-enriched heterogeneous clusters, but the strength and stability of this transition are lineage specific.

### 3.8. Comparative Fourier Spectral Profiling of Primate SCFRs

#### 3.8.1. Periodic Signatures and Codon-Level Periodicity

To detect latent periodic structure within SCFR sequences, we applied the discrete Fourier transformation to convert linear nucleotide arrangements into their corresponding frequency spectra. Fourier analysis enables quantitative detection of repeating biological patterns, as regularly spaced sequence features appear as distinct spectral peaks whose amplitudes reflect signal strength. In this framework, frequency represents the recurrence rate of a pattern along the sequence, whereas amplitude measures the magnitude of that periodic contribution.

Across all analysed primates, Fourier profiling revealed a clear dichotomy between SCFRs overlapping annotated coding exons and those located entirely within non-coding regions (**Supplementary Figures 45-51**). SCFRs with coding overlap consistently exhibited a dominant spectral peak near ∼0.33, corresponding to the canonical three-nucleotide periodicity imposed by the genetic code. In contrast, SCFRs lacking coding overlap frequently shifted their primary peak toward ∼0.17-0.18, indicating an alternative periodic regime. This shift suggests that, in the absence of translational constraints, long SCFRs are structured not by codon organisation but by intrinsic sequence architecture, such as repeat-associated or motif-driven periodicity. Together, these results demonstrate that Fourier spectra effectively discriminate coding-derived SCFRs from structurally organised non-coding regions.

#### 3.8.2. Species-Specific Variation and Spectral Stability

Although the bimodal structure of SCFR spectra was broadly conserved, the dominance and stability of specific frequency peaks varied across species and length thresholds. In human, SCFRs ≥5 kb displayed a pronounced and stable peak at ∼0.33 with large peak separations, indicating strong codon-level periodicity within long regions and reinforcing their coding enrichment (**Supplementary Figures 47**). In contrast, gorilla frequently retained a dominant ∼0.17 peak even among long SCFRs containing coding overlap, a pattern consistent with the influence of exceptionally large non-coding structural outliers identified in its assembly (**Supplementary Figure 4**). Most species maintained bimodal distributions across thresholds; however, the Sumatran orangutan exhibited a unimodal distribution at ≥10 kb in non-coding SCFRs, suggesting the presence of highly uniform, lineage-specific repetitive architectures that override codon-scale signals. Collectively, these findings indicate that while codon-level periodicity is a conserved hallmark of coding-enriched SCFRs, the spectral landscape of long SCFRs differs markedly among primates. This variation likely reflects lineage-specific balances between translational constraint and repeat-driven sequence organisation, underscoring distinct evolutionary routes through which extended ORF-like regions emerge and persist.

## 4. Discussion

In this study, we define stop-codon-free regions (SCFRs) as uninterrupted stretches of DNA lacking in-frame stop codons within a given reading frame and use them as a genome-wide proxy for latent coding potential. Gene duplication has traditionally been viewed as the primary mechanism underlying the origin of new genes, yet increasing evidence suggests that *de novo* emergence from non-coding DNA is more common than previously assumed [1–3,8]. Because newly emerged genes or proto-genes often lack detectable similarity to known proteins, their identification and systematic characterisation remain challenging [13,27]. To address this, we adopt an unbiased sequence-based approach: instead of relying on homology or annotation, we quantify SCFRs across entire genomes and examine their global distribution, compositional properties, and local genomic context to assess the structural preconditions for coding potential within non-coding DNA. By leveraging complete telomere-to-telomere assemblies from seven primates, we establish a comprehensive framework that links SCFR abundance, length constraints, compositional architecture, and coding-associated signatures. Rather than focusing solely on annotated coding regions, we systematically map the broader landscape of uninterrupted ORF-like sequences across whole genomes, identifying where extended SCFRs persist and how their structure and sequence features relate to non-coding compartments. This approach enables a genome-wide view of how coding-compatible sequence is distributed and constrained, providing a foundation for understanding the sequence-level substrate from which *de novo* genes may arise.

The investigation into the genomic landscape of seven primate species reveals that SCFRs are ubiquitous, numbering approximately 300 million per genome. While the vast majority of these regions are compact with a median length of roughly 39 bp, they represent a massive, strand-neutral reservoir of potential proto-genes. However, this reservoir narrows rapidly with increasing length. As minimum length thresholds increase, SCFR counts and genome coverage decline precipitously, with more than 99.7% of regions shorter than 500 bp and fewer than a few hundred exceeding 10 kb. In parallel, longer SCFRs are increasingly concentrated within coding exons, whereas short SCFRs remain predominantly in non-coding DNA. This shift indicates that extended uninterrupted reading frames are rarely sustained outside established genes. Base composition further constrains this persistence as long SCFRs are largely confined to moderate GC environments, whereas short SCFRs occur across a broader compositional spectrum. These observations refine the view of non-coding DNA as uniformly permissive for proto-gene formation. Although short ORF-like sequences are widespread, the emergence of long coding-compatible regions is tightly constrained by sequence length, nucleotide composition, and local genomic context. Understanding how these constraints operate in specific genomic compartments, therefore, becomes critical for identifying which SCFRs may realistically contribute to *de novo* gene evolution.

While gene deserts have traditionally been defined as extended gene-free intervals, accumulating evidence indicates that they are not functionally inert, as several studies have demonstrated their involvement in long-range regulatory interactions that modulate the expression of neighbouring genes [21,28–30]. Nevertheless, no protein-coding genes have been formally annotated within these regions, and their potential to generate proto-genes has remained largely unexplored. Here, we show that gene deserts harbour hundreds of long SCFRs (≥5 kb), all entirely embedded within these intervals, with clear length-dependent enrichment that underscores the presence of substantial ORF-like sequence space in deep intergenic regions. Many of these desert-derived ORFs exhibit similarity to repetitive, low-complexity, or structurally biased protein families rather than to well-characterised conserved genes; notably, mucin-like and proline-rich families show detectable homology across all seven primate species. In addition, a subset displays codon-scale periodicity consistent with bona fide coding architecture. Collectively, these findings extend the functional landscape of gene deserts beyond distal regulation and support the hypothesis that they may serve as evolutionary reservoirs for proto-gene emergence, offering concrete candidates for future functional and evolutionary investigation.

To examine how SCFRs operate within established genes and whether they might provide latent coding potential for gene extension, we defined the concept of an exon shadow. An exon shadow represents the uninterrupted SCFR sequence that extends beyond an annotated exon boundary, either upstream or downstream. In other words, when an exon ends but the reading frame continues without encountering a stop codon, the remaining in-frame stretch forms a potential extension “shadow” of that exon. This framework allows us to assess whether existing genes are embedded within longer ORF-compatible contexts that could, in principle, facilitate exon elongation, boundary shifts, or gradual coding expansion. Across species, most genes contained at least one exon associated with a non-zero shadow, and in most cases, the SCFR extended beyond the annotated exon boundary. Shadow lengths were typically short but showed clear enrichment at codon-compatible intervals and a consistent ∼3 bp periodicity, indicating that these extensions preserve reading-frame structure rather than arising randomly. Shadows were predominantly associated with constitutive exons and were enriched in genes involved in cytoskeletal organisation, signalling, and adhesion. These observations raise important questions. If many exons sit within uninterrupted, in-frame extensions, under what conditions might these latent segments be incorporated into transcripts or translated into functional protein sequence? Do alternative splicing, mutation, or boundary shifts occasionally convert shadows into bona fide coding regions? By defining exon shadows, we establish a framework to explore how existing genes might extend their coding potential and how gradual boundary remodelling could contribute to evolutionary innovation.

While examining exon shadows, we observed that in some cases a single SCFR spans an entire intron without encountering an in-frame stop codon. Such introns are structurally poised for complete retention while preserving coding potential, fitting the definition of exitrons described by Marquez et al. 2015 [31]. Across primates, we identified ∼2,340 such exitron candidates per genome, with lengths constrained at the lower end but variably extended across lineages. Functionally, exitron-containing genes were consistently enriched in extracellular matrix and cytoskeletal pathways, yet also showed lineage-specific associations, suggesting both conserved and adaptable roles. Notably, exitrons displayed the strongest coding-like nucleotide signatures, such as high GC, elevated GC3, and structured stretch architecture, surpassing even annotated exons, while exon shadows showed intermediate, direction-dependent profiles. These findings position SCFR-spanned introns as a transitional layer between canonical introns and coding exons, expanding the exon-shadow concept from boundary extensions to whole-intron coding compatibility. If shadows represent potential exon elongation, exitrons represent potential exon internal expansion through regulated intron retention. Taken together, this framework treats uninterrupted reading frames as latent extensions of gene structure and directs attention to concrete candidates for evaluating how splicing regulation and sequence bias interact during coding evolution.

To determine whether long SCFRs begin to resemble a genuine coding sequence, we examined codon usage patterns across increasing length thresholds. As shorter, repeat-rich regions were progressively excluded, long SCFRs showed clearer compositional structuring, with GC and particularly GC3 emerging as the dominant axis separating low-GC, simple-repeat–associated regions from higher-GC regions enriched for coding overlap. Coding-associated SCFRs were not randomly distributed but concentrated within specific heterogeneous clusters, while repeat-driven homogeneous clusters diminished with length. Notably, SINE elements were enriched in many SCFRs, especially at lower thresholds. Because the SINE/Alu family of repeat elements contain internal promoters, their retention within extended SCFRs may increase the likelihood of transcriptional activation, providing a potential route toward proto-gene emergence [32–34]. These patterns indicate that length filtering does more than reduce SCFR number, it preferentially enriches for regions with gene-like codon organisation while gradually excluding repeat-dominated background. At the same time, the uneven contribution of different repeat classes and the lineage-specific differences in codon architecture suggest that the shift toward coding-like composition is neither uniform nor purely stochastic.

We next assessed whether long SCFRs display intrinsic coding-like organisation by analysing their sequence periodicity with the Fourier transformation. SCFRs overlapping annotated exons showed a strong peak at ∼0.33, consistent with three-nucleotide codon periodicity [35], whereas non-coding SCFRs frequently exhibited a dominant peak around ∼0.17-0.18, reflecting repeat- or motif-driven structure rather than translational constraint. This bimodal spectral pattern was broadly conserved across primates, though its strength varied among lineages and length thresholds. This suggests SCFR length alone is insufficient to infer coding potential, as only a subset of long SCFRs carries the codon-scale periodic signal expected of translated regions. Fourier analysis, therefore, adds an orthogonal structural criterion, refining the identification of which long ORF-like regions warrant further evolutionary and functional evaluation.

Overall, this study reframes uninterrupted reading frames as a measurable genomic substrate rather than a rare anomaly. Across primates, SCFRs are abundant but highly filtered by length, composition, repeat context, and periodic structure, with only a small fraction exhibiting features consistent with genuine coding potential. By integrating genomic distribution, exon boundary architecture, intron retention potential, codon usage, repeat associations, and spectral periodicity, we move beyond simple ORF counting toward a multidimensional definition of coding compatibility. This framework does not claim that long SCFRs are genes, but it systematically narrows the landscape to those regions most plausibly positioned for evolutionary testing. Future work integrating transcriptional, translational, and population-level constraint data will be essential to determine which of these structurally primed regions transition from latent sequence to functional gene.

## Conclusions

In conclusion, our study systematically maps stop-codon-free regions (SCFRs) across seven primate telomere-to-telomere genomes and establishes a genome-wide framework for evaluating latent coding potential independent of annotation. We observe that SCFRs are abundant but sharply constrained by length, with longer regions becoming rare and increasingly associated with annotated coding overlap, moderate GC enrichment, and structured exon–intron contexts. Our findings reveal that exon shadows frequently extend beyond annotated exon boundaries, exposing coding-compatible sequence adjacent to established genes, and that some introns are fully spanned by single SCFRs, consistent with exitron-like architectures. We further observe that progressive length filtering enriches for gene-like codon usage and compositional features, while Fourier spectral analysis distinguishes a subset of long SCFRs exhibiting codon-scale periodicity from those dominated by repeat-driven structure. Together, our findings demonstrate that uninterrupted length alone does not define coding potential and provide a multidimensional basis for identifying genomic regions most plausibly positioned to contribute to *de novo* gene emergence in primates.

## Ethics

This study did not require approval from a human ethics committee or an animal welfare board.

## Data accessibility

Scripts and supplementary data are available at https://github.com/AswinSSoman/SCFR.git.

## Author’s contributions

NV: Conceptualisation, Funding acquisition, Recourses, Project administration, Supervision, Project administration, Writing-review and editing; ASS: Formal analysis, Investigation, Visualisation, Validation, Writing-original draft; GS: Formal analysis, visualisation; AD: Visualisation, Writing-original draft; GSP: Visualisation, Writing-Supplementary; CS: Visualisation, Writing-Supplementary; DB: Visualisation.

All authors gave final approval for publication and agreed to be accountable for the work reported.

## Acknowledgement

The Department of Biotechnology, Ministry of Science and Technology, India (Grant no. BT/11/IYBA/2018/03) and Science and Engineering Research Board (Grant no. ECR/2017/001430) provided funds used to generate primary sequencing data published in this article and computational resources (i.e., Har Gobind Khorana Computational Biology cluster) used. The authors acknowledge the use of artificial intelligence (AI) tools solely for paraphrasing purposes during the preparation of this manuscript.

