## Supplementary Figures for "The Landscape of Stop Codon-Free Regions in Primates: A Reservoir of Proto-Genes"

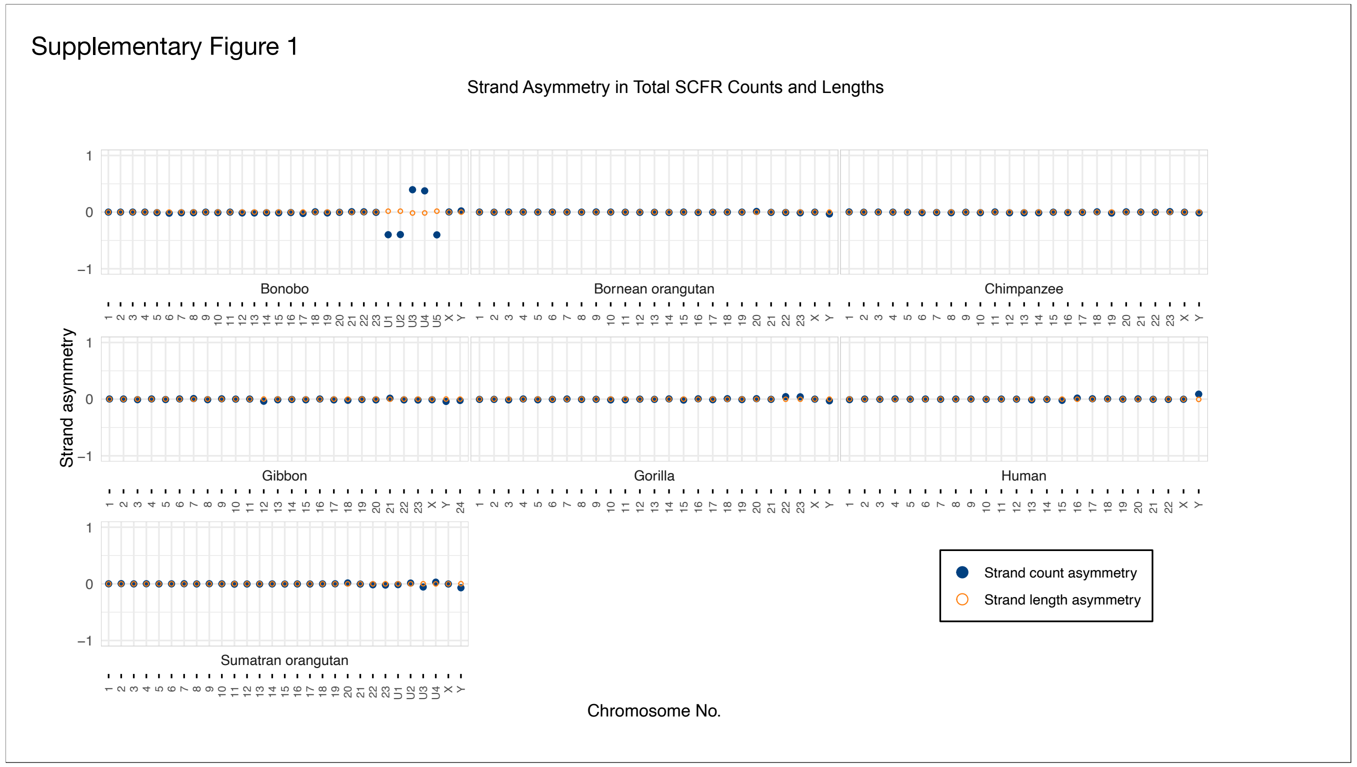


**Supplementary Figure 1: Strand asymmetry in Total SCFR Counts and Lengths.**
This plot displays strand asymmetry for all chromosomes across the seven primate species. The x-axis denotes chromosome identifiers for each species, while the y-axis represents strand asymmetry values ranging from −1 to 1. Blue filled points indicate asymmetry based on SCFR counts, and open yellow circles indicate asymmetry based on the total lengths of SCFRs per strand.

Supplementary Figure 2

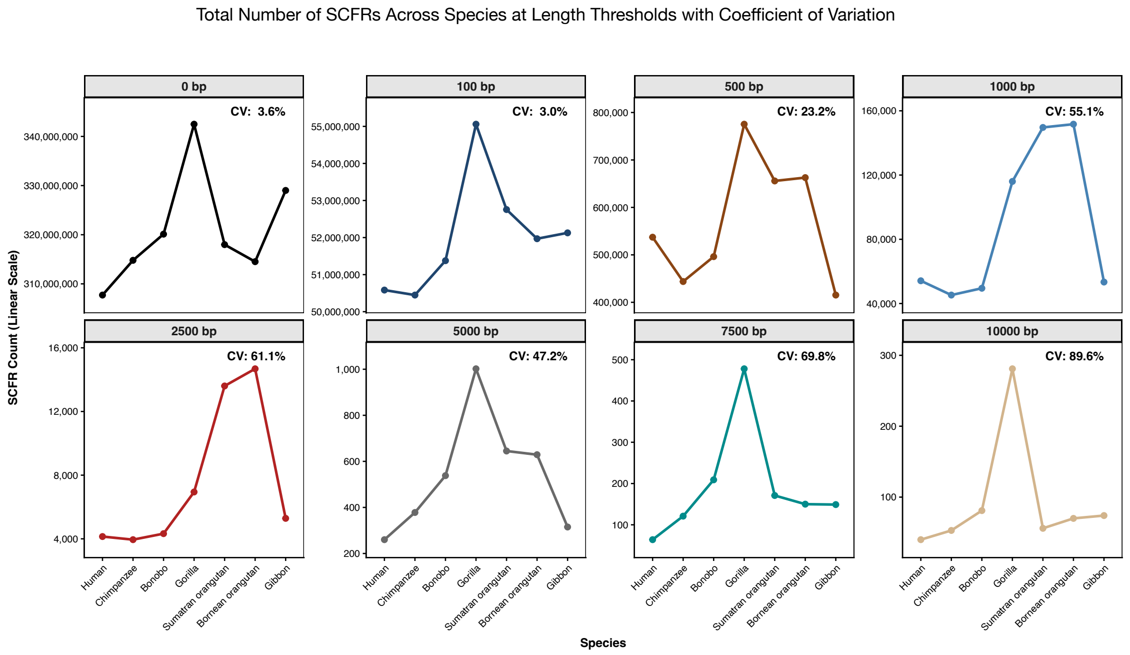


**Supplementary Figure 2: Total Number of SCFRs Across Species at Length Thresholds with Coefficient of Variation.**
This line plot illustrates the total number of unfiltered SCFRs in each species across increasing minimum length thresholds, shown in separate panels. The coefficient of variation (CV) is reported for each threshold to highlight variability in SCFR abundance among species.

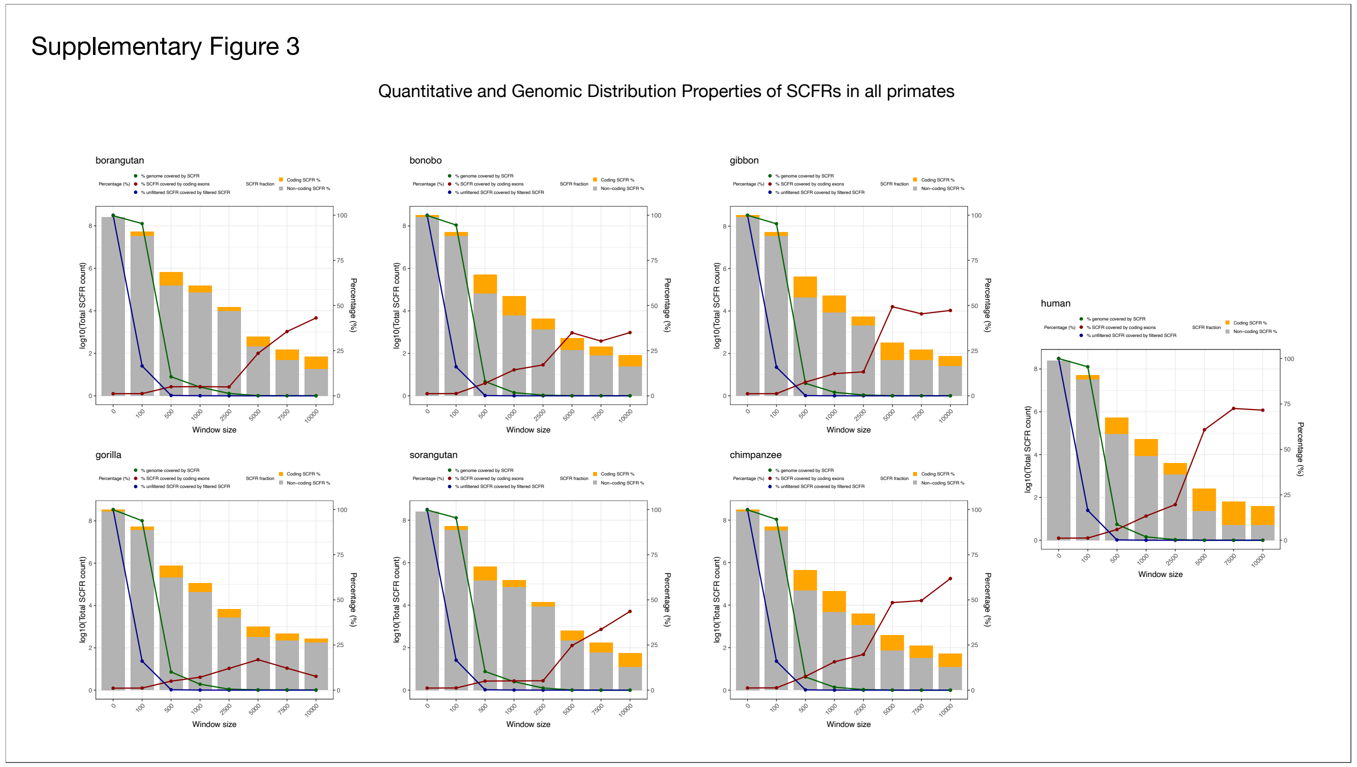


**Supplementary Figure 3: Quantitative and Genomic Distribution Properties of SCFRs in all primates.**
The X-axis represents minimum SCFR length cutoffs (bp), while the left Y-axis displays the total number of SCFRs on a $\log_{10}$ scale, and the right Y-axis shows percentage covered. The bars depict the SCFR count at each threshold, with stacked segments indicating the proportions intersecting coding exons (orange) and non-coding genomic regions (grey). The lines represent percentage covered in terms of length: green denotes the percentage of the genome covered by SCFRs, red indicates the percentage of SCFR bases overlapping coding exons, and blue shows the percentage of unfiltered SCFRs (0-bp threshold) retained after applying each length filter.

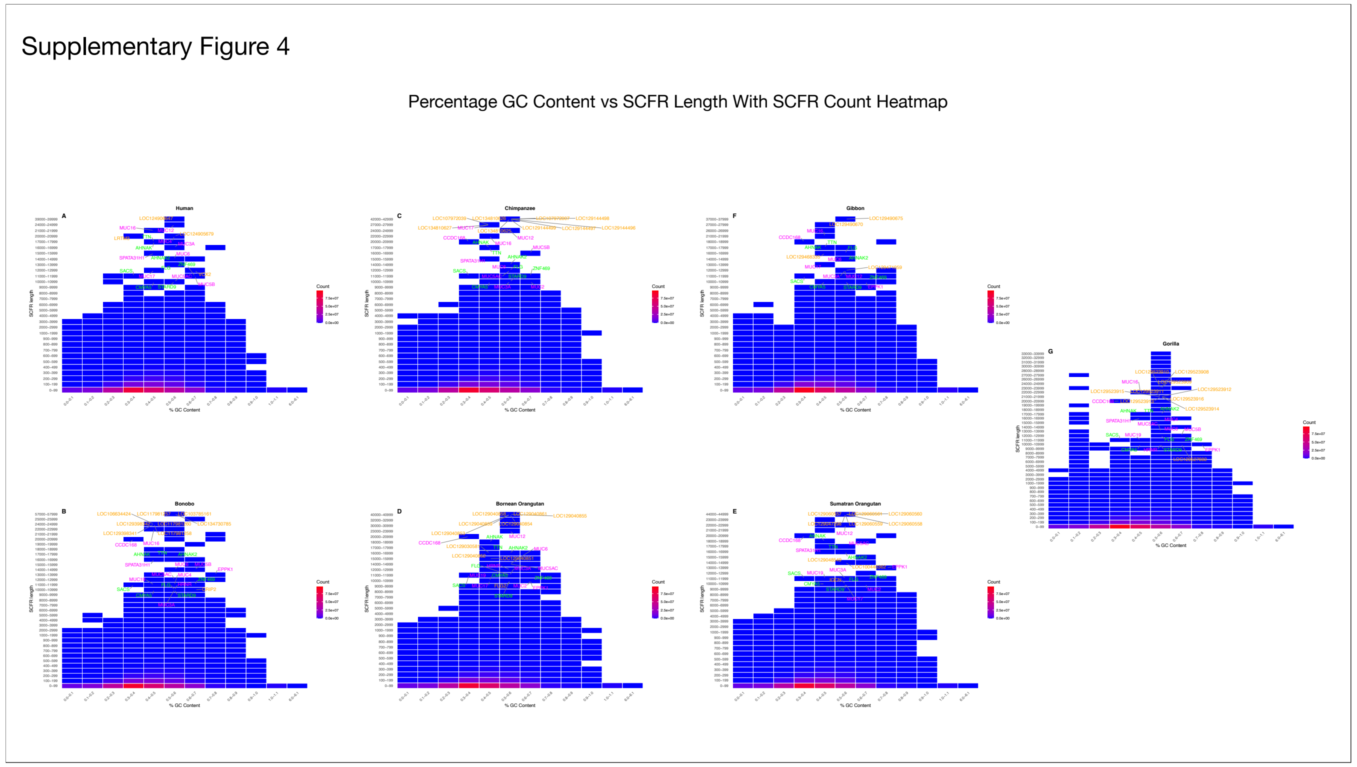


**Supplementary Figure 4: Percentage GC Content vs SCFR Length With SCFR Count Heatmap.**
This heatmap displays the total number of SCFRs in the seven primate genomes across two dimensions: SCFR sequence length (y-axis; binned in 100 bp increments below 1 kb and 1 kb increments above 1 kb) and % GC content (x-axis; binned in 0.1 fractional units). The colour scale represents SCFR frequency per bin, with warmer colours indicating higher abundance. Gene names labelled adjacent to long-SCFR bins denote protein-coding genes that intersect with SCFRs longer than 10 kb. The colours green, magenta, and orange indicate genes that appear in all species, at least in two species, and in a unique species, respectively.

Supplementary Figure 5

Percentage AT Content vs SCFR Length With SCFR Count Heatmap

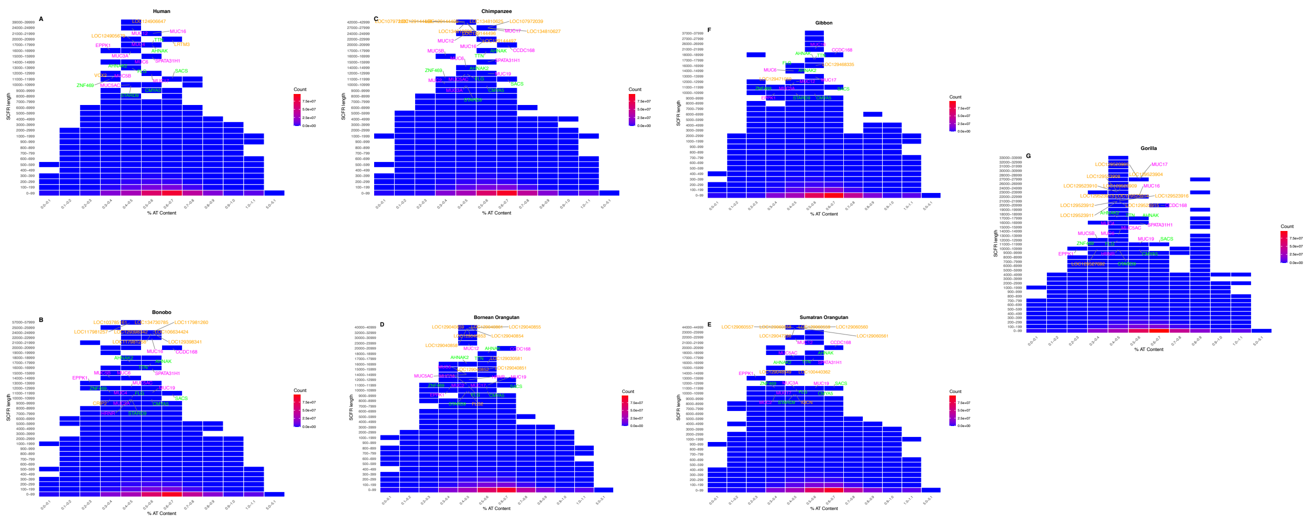


**Supplementary Figure 5: Percentage AT Content vs SCFR Length With SCFR Count Heatmap.**
This heatmap displays the total number of SCFRs in the seven primate genomes across two dimensions: SCFR sequence length (y-axis; binned in 100 bp increments below 1 kb and 1 kb increments above 1 kb) and % AT content (x-axis; binned in 0.1 fractional units). The colour scale represents SCFR frequency per bin, with warmer colours indicating higher abundance. Gene names labelled adjacent to long-SCFR bins denote protein-coding genes that intersect with SCFRs longer than 10 kb. The colours green, magenta, and orange indicate genes that appear in all species, at least in two species, and in a unique species, respectively.

### Supplementary Figure 6

Length Distribution of Intergenic Region and Gene Deserts

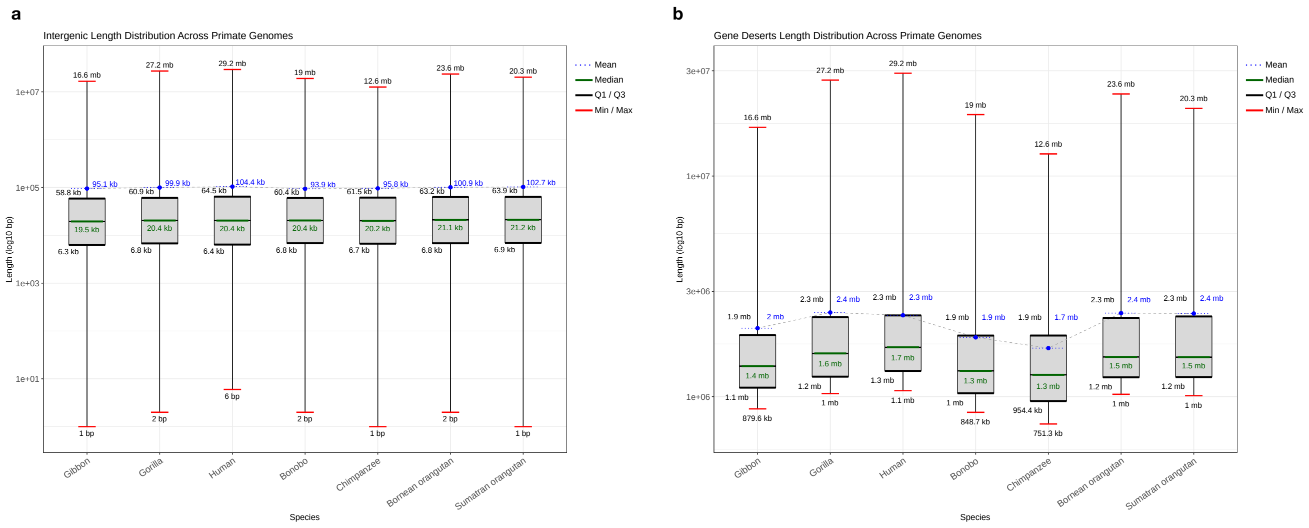


**Supplementary Figure 6: Length Distribution of Intergenic Region and Gene Deserts across seven primates:**

(a) Intergenic regions: Boxplots show the distribution of total intergenic region lengths across species (x-axis), using the base-10 logarithm of length on the y-axis. For each species, minimum, maximum, mean, median, first quartile, and third quartile values are indicated.

(b) (a) Gene deserts: Boxplots show the total length distribution of identified gene deserts per species (x-axis), on a $\log_{10}$-transformed length scale (y-axis). Each box displays the minimum, maximum, mean, median, first quartile, and third quartile for that species.

Supplementary Figure 7

Length Distribution of the Overlap Between SCFR (>=5kb) and Gene Deserts

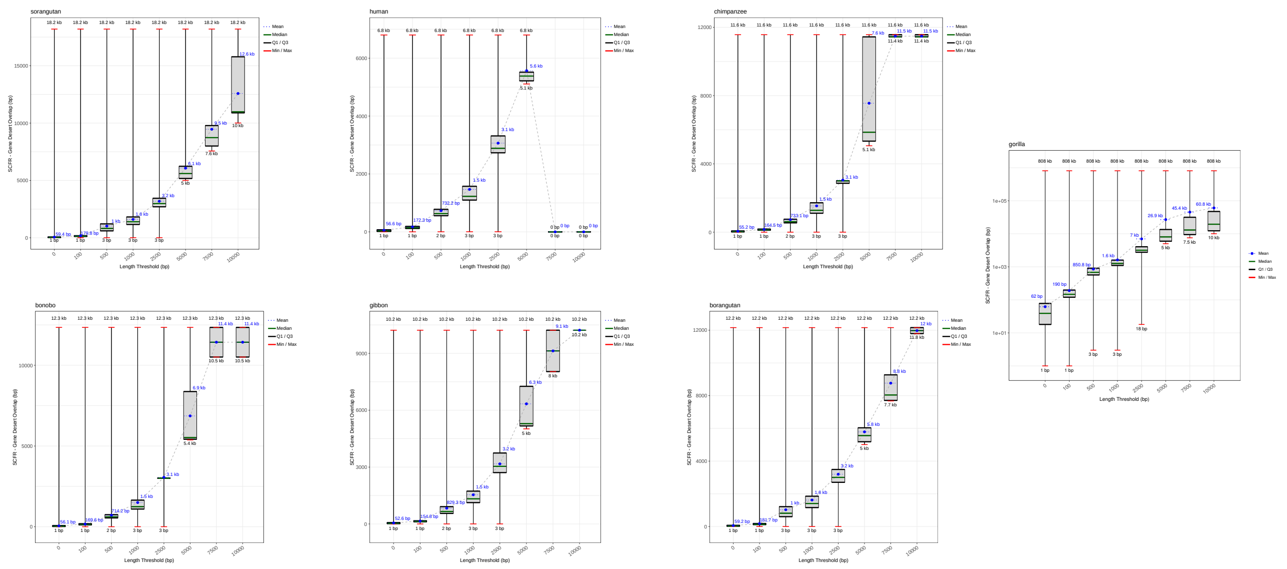

**Supplementary Figure 7: Length Distribution of the Overlap Between SCFR (>=5kb) and Gene Deserts.**
This figure shows how the genomic overlap between SCFRs and gene deserts varies across increasing SCFR length thresholds for each species. The x-axis represents minimum SCFR lengths, and the y-axis reports their overlap with gene deserts in base pairs, with the minimum, maximum, median, quartiles, and mean overlap indicated for each threshold. SCFRs ≥5 kb consistently show complete containment within gene deserts, with all statistics converging at full overlap values. In contrast, SCFRs at thresholds of 2.5 kb and below display minimal or negligible overlap, with ranges extending from 0 to on**ly 18 bp.**

Supplementary Figure 8

Percentage of shadows and exitrons

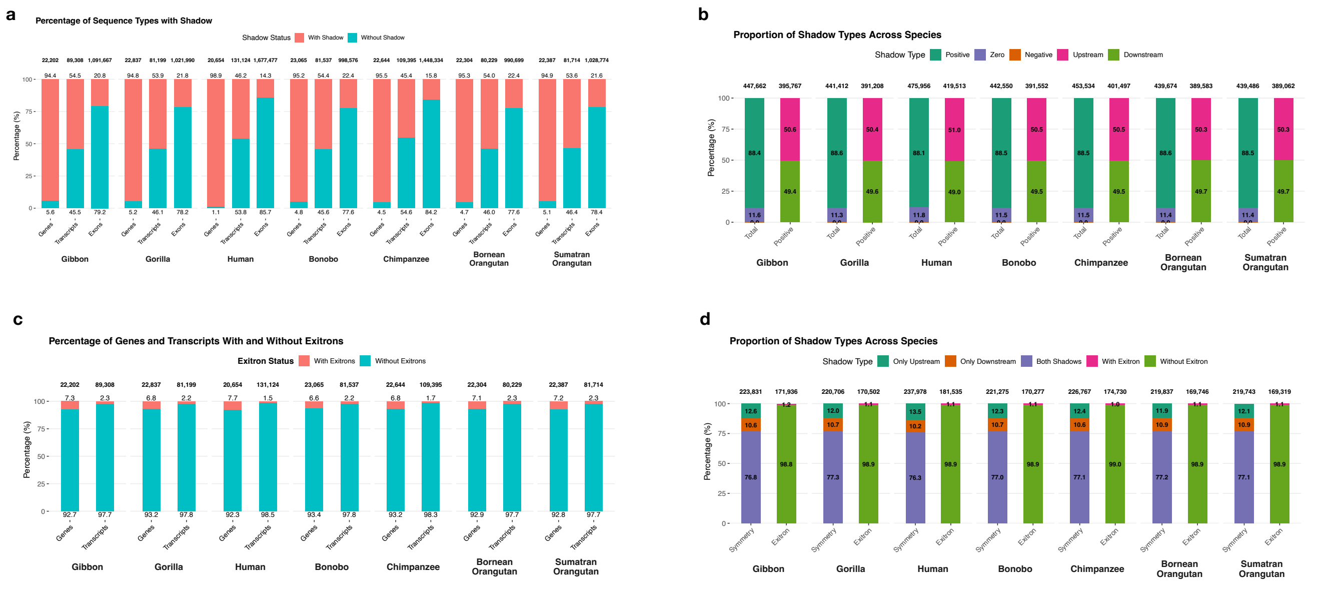


**Supplementary Figure 8:**

**(a) Percentages of genomic regions with and without shadows**

The x-axis is two-layered, with the lower layer representing the species and the upper layer representing three genomic regions (genes, transcripts and exons). The y-axis is the percentage of genomic regions with and without shadows. The red stacks represent region with shadow, while the teal stacks represent region without shadow. The percentages are annotated below and above the respective bars. The total number of genes, transcripts and exons identified in the particular species are annotated above the respective bars for that genomic region.

**(b) Percentages of shadow types across species**

The x-axis is arranged in two tiers, with species shown in the lower layer and shadow categories (overall and positive) in the upper layer. The y-axis indicates the percentage of shadows. For each species, the left bar shows shadow polarity: positive shadows (dark green), zero shadows (purple), and negative shadows (orange). The right bar depicts shadow position, with downstream shadows shown in light green and upstream shadows in pink. The total counts of overall shadows and positive shadows are indicated above each bar, while the percentage contribution of each shadow type is annotated within the corresponding stacked segments.

**(c) Percentage of genes and transcripts with and without exitrons**

The x-axis is arranged in two tiers with the lower tier showing species and upper tier showing genomic regions (genes and transcripts). The y-axis indicates the percentage of region with and without exitrons. The red stacks represent regions with exitrons while teal stacks represent regions without exitrons. The percentages are annotated above and below the respective bars. The total number of genes and transcripts for the respective species are annotated above the respective bars.

**(d) Percentage of shadow pair types across species**

The x-axis is shown in two tiers, with species on the lower tier and shadow pair symmetry and exitron status on upper tier. The y-axis represents bar-specific percentages: the left stacked bar shows the proportion of SCFR-exon pairs with each shadow configuration (blue: both upstream and downstream shadow; orange: only downstream shadow; green: only upstream shadow), while the right stacked bar shows the proportion of SCFRs without (light green) and with (pink) exitrons. Percentages are annotated within the stacks, and total counts of SCFR-exon pairs and SCFRs are annotated above the left and right bars respectively.

Supplementary Figure 9

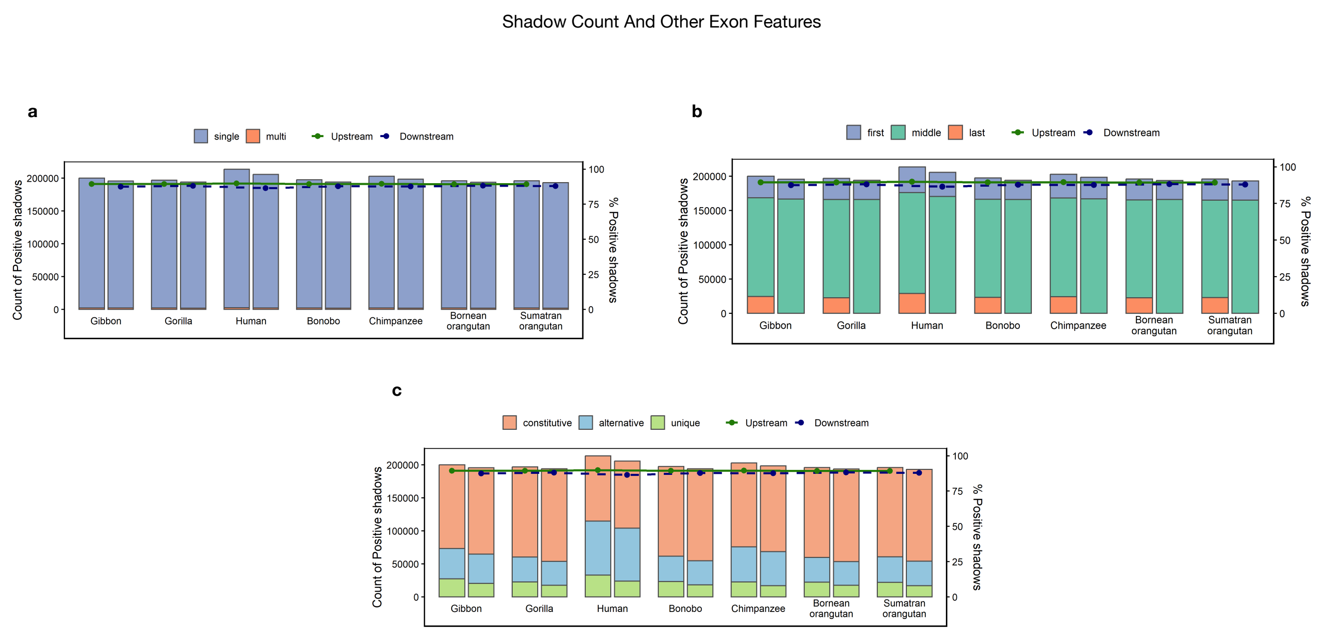


**Supplementary Figure 9:**

**(a) Positive shadow distribution across single- and multi-exon SCFR–exon pairs**

The x-axis represents species. The left y-axis shows total number of positive shadows, while the right y-axis shows percentage of positive shadows among all shadows. Bars marked by a green dot on the solid green line indicate upstream shadows, while bars marked with a blue dot on the dotted blue line indicate downstream shadow. Each bar is stacked, with the blue segment representing single-exon SCFR-exon pairs and the orange segment representing multi-exon pairs.

**(b) Positive shadow distribution across first, middle and last exons**

The x-axis represents species. The left y-axis shows total number of positive shadows, while the right y-axis shows percentage of positive shadows among all shadows. Bars marked by a green dot on the solid green line indicate upstream shadows, while bars marked with a blue dot on the dotted blue line indicate downstream shadow. Each bar is stacked, with the blue portions representing first exons, green portions representing middle exons and orange portions representing last exons.

**(c) Positive shadow distribution across constitutively, alternatively and uniquely spliced exons**

The x-axis represents species. The left y-axis shows total number of positive shadows, while the right y-axis shows percentage of positive shadows among all shadows. Bars marked by a green dot on the solid green line indicate upstream shadows, while bars marked with a blue dot on the dotted blue line indicate downstream shadow. Each bar is stacked, with the orange portion representing constitutively spliced, blue portions representing alternatively spliced and green portion representing uniquely spliced exons.

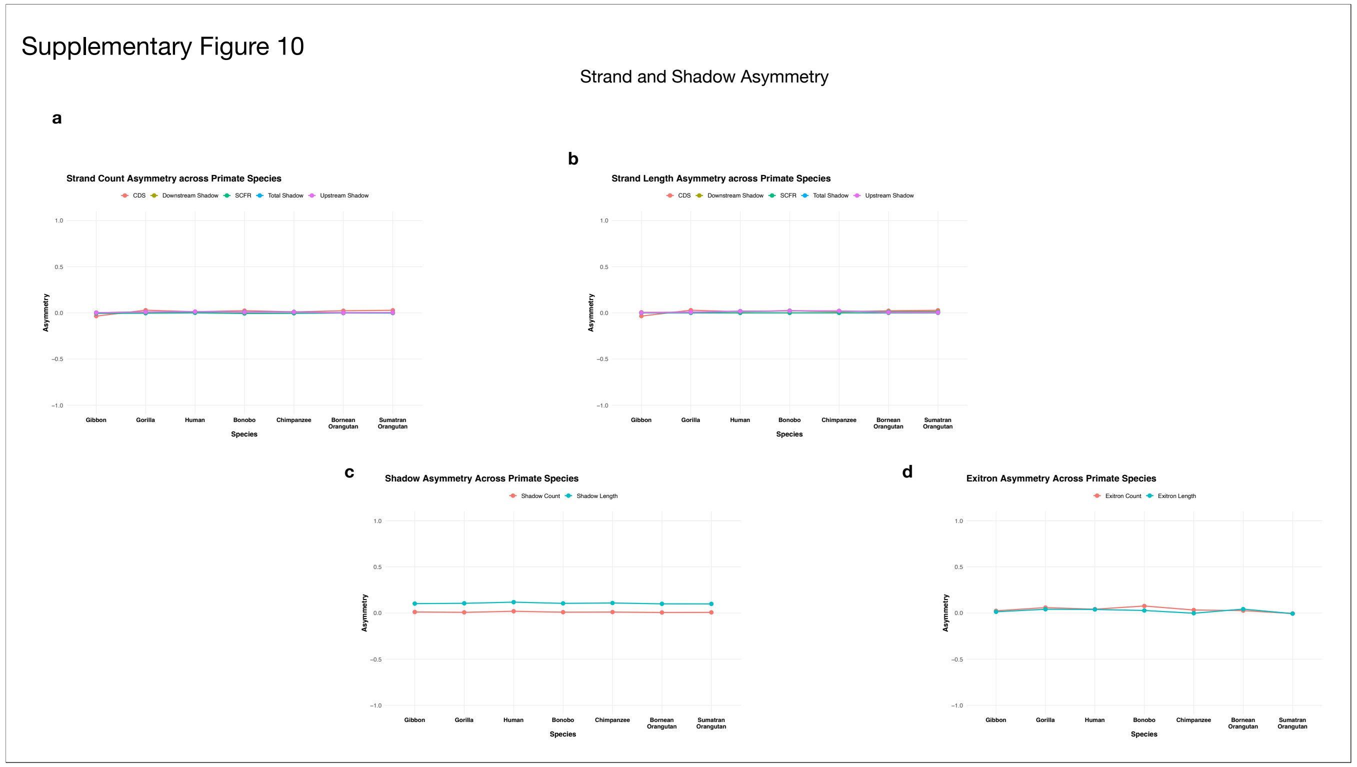


**Supplementary Figure 10:**

**(a) Strand Count Asymmetry across Primate species**

The x-axis represents species, and the y-axis represents strand count asymmetry values ranging from −1 to 1. Pink, olive green, green, blue, and light pink lines represent asymmetry in CDS, downstream shadows, SCFRs, overall shadows, and upstream shadows, respectively.

**(b) Strand Length Asymmetry across Primate Species**

The x-axis represents species, and the y-axis represents strand length asymmetry values ranging from −1 to 1. Pink, olive green, green, blue, and light pink lines represent asymmetry in CDS, downstream shadows, SCFRs, overall shadows, and upstream shadows, respectively.

**(c) Shadow Length and Count Asymmetry across Primate Species**

The x-axis represents species, and the y-axis represents asymmetry values ranging from −1 to 1. The orange and turquoise lines indicate asymmetry in shadow count and length respectively.

**(d) Exitron Length and Count Asymmetry across Primate Species**

The x-axis represents species, and the y-axis represents asymmetry values ranging from −1 to 1. The orange and turquoise lines indicate asymmetry in exitron count and length respectively.

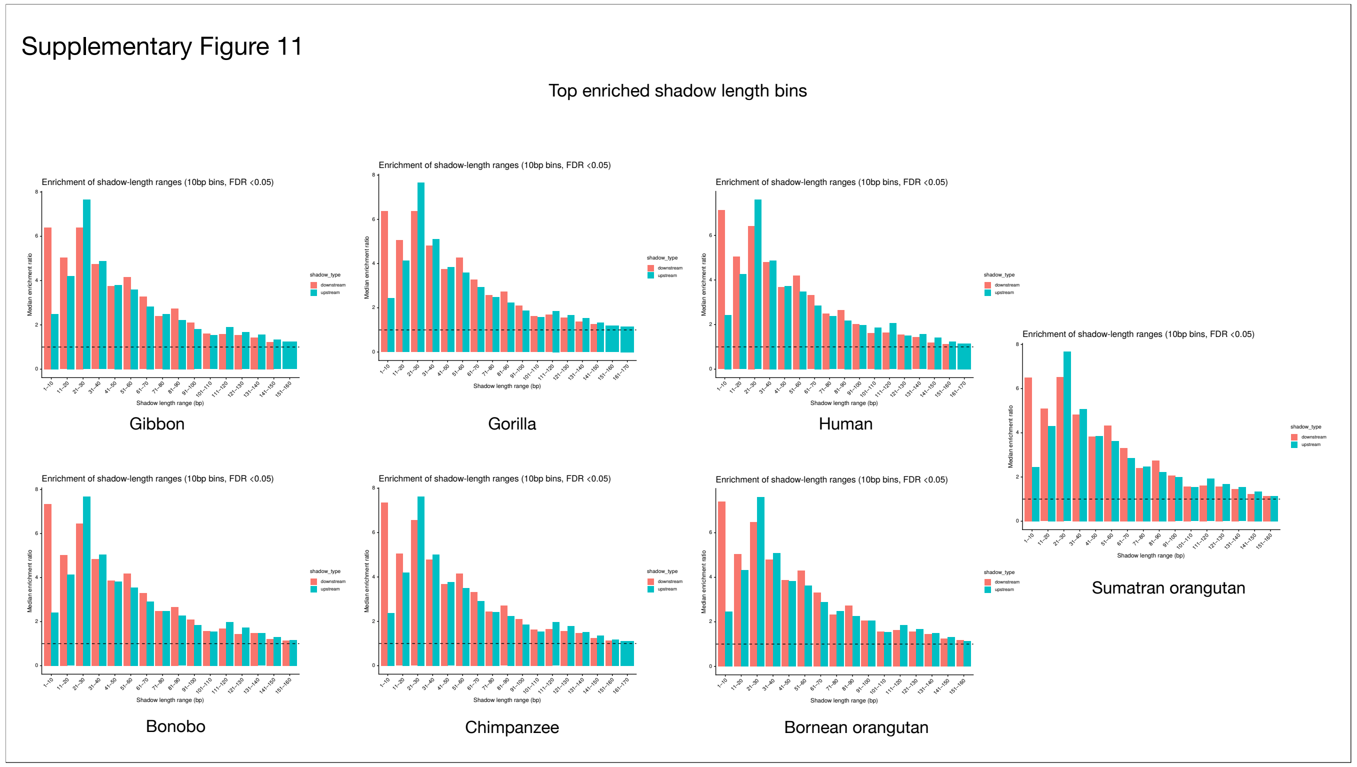


**Supplementary Figure 11:**

**Shadow Length Enrichment in Upstream and Downstream Regions Across Seven Primate Species**

Grouped bar plots depict the enrichment of shadow lengths across all seven primate species, evaluated separately for upstream and downstream shadows. The x-axis represents shadow length ranges, and the y-axis shows the median enrichment ratio relative to a uniform expectation. Green bars indicate enrichment in upstream shadows, while red bars indicate enrichment in downstream shadows.

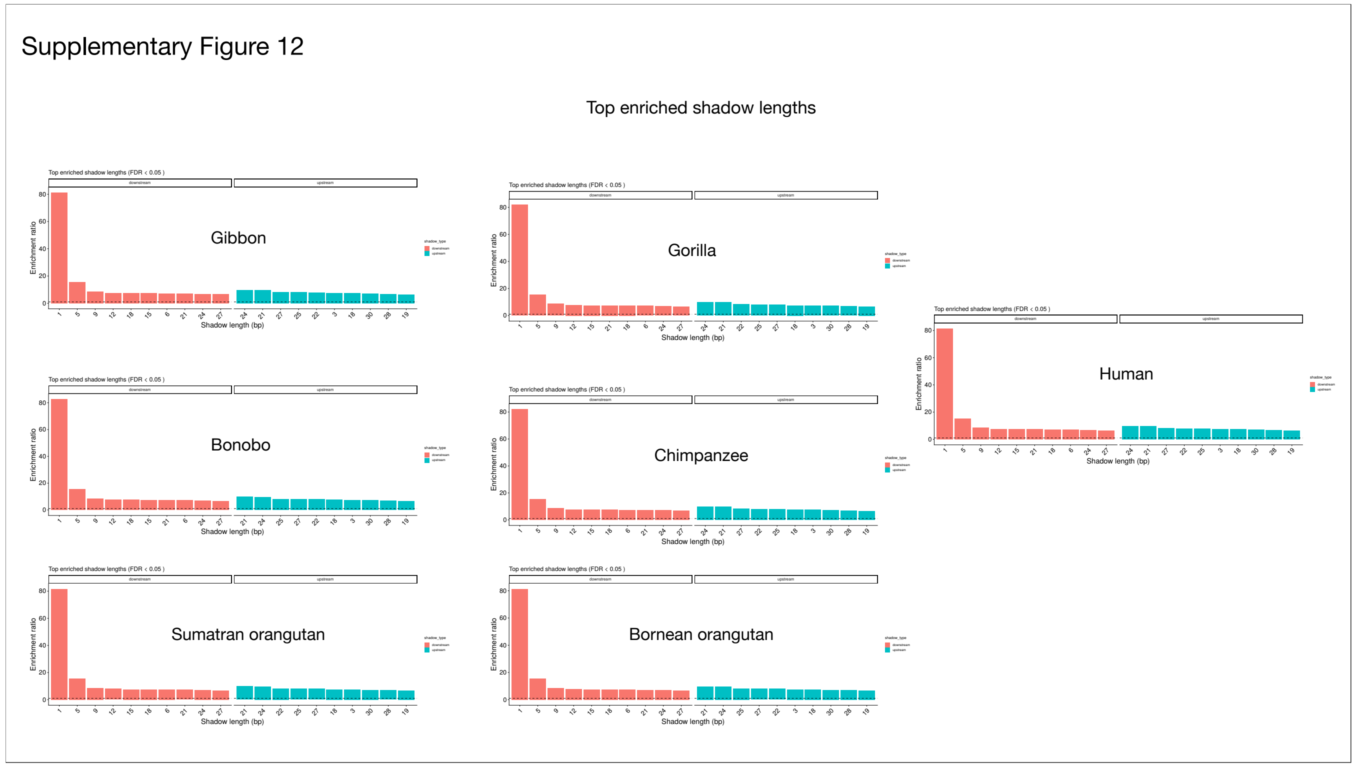


**Supplementary Figure 12:**
The plot shows the top 10 enriched shadow length values across primate species. In each panel, the x-axis represents enriched shadow length values and the y-axis represents the enrichment ratio. Orange bars correspond to downstream shadows, while turquoise bars correspond to upstream shadows.

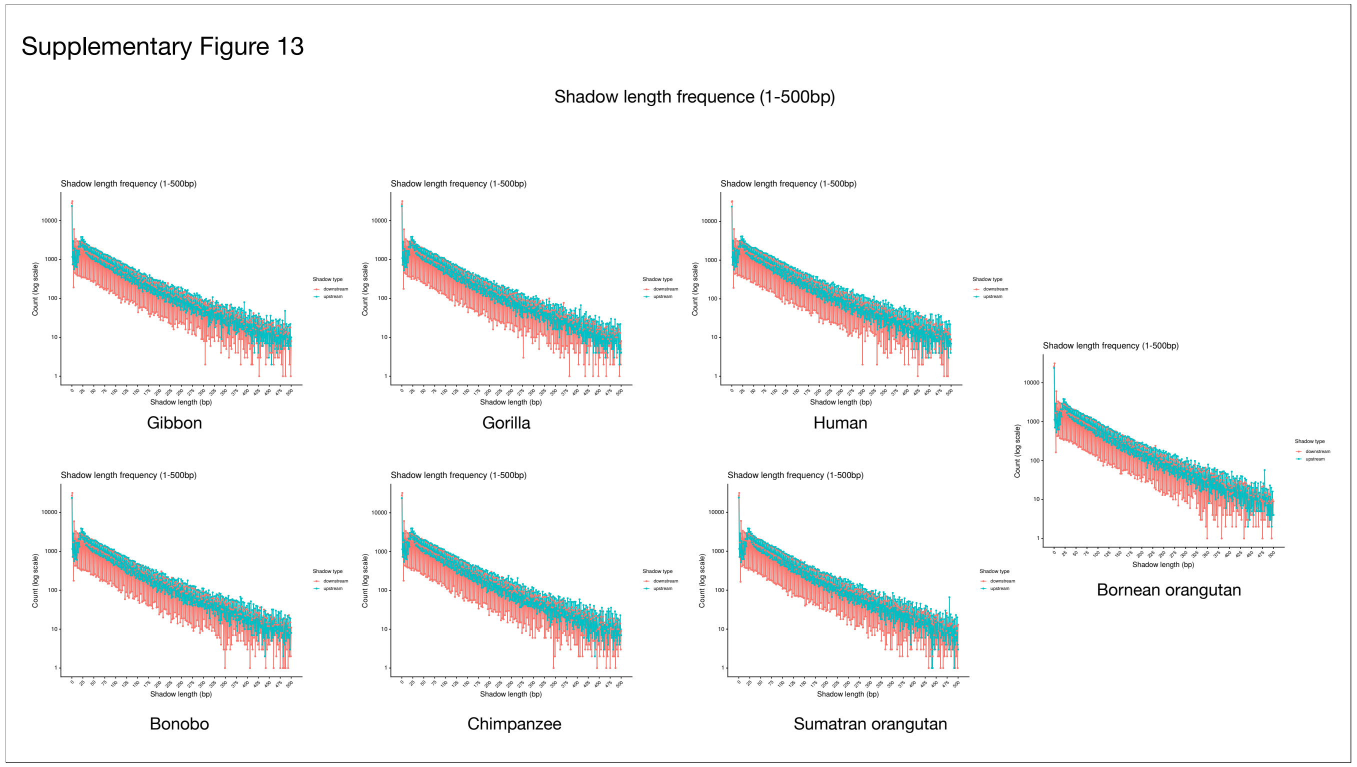


**Supplementary Figure 13:**
The plot shows the frequency distribution of shadow length values (≤500 bp) in primate species. The x-axis represents shadow length, and the y-axis represents the log10-transformed count of shadows at each length, with original counts annotated. Orange and turquoise lines represent upstream and downstream shadow length frequencies, respectively. Each highlighted dot on the lines indicates the frequency at the corresponding shadow length.

### Supplementary Figure 14

Gene Set Enrichment Analysis

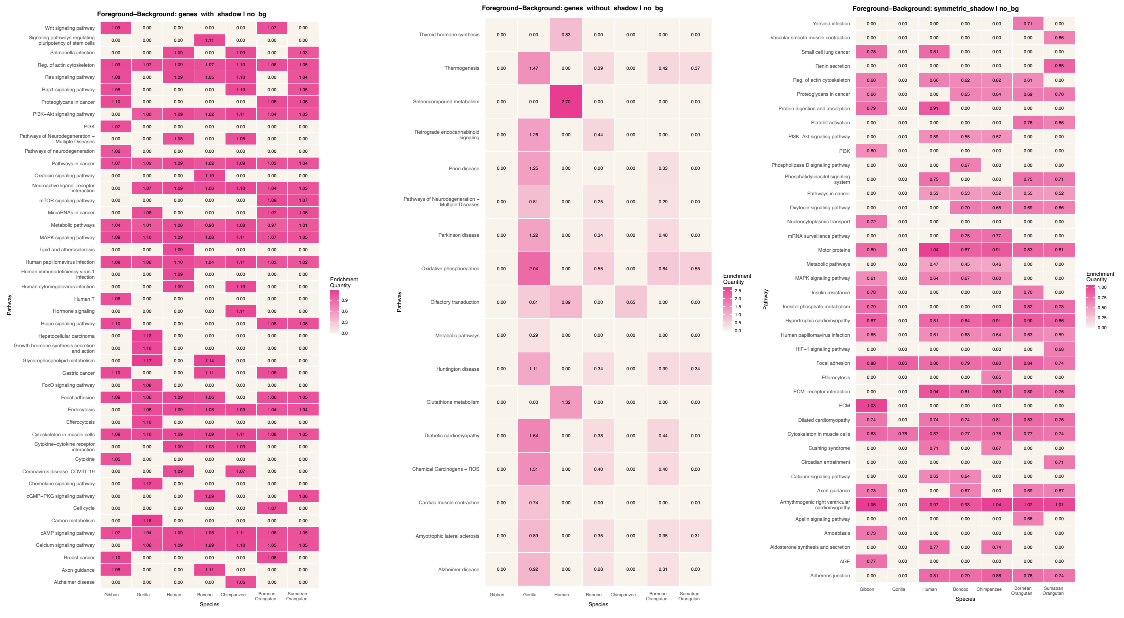

(a) Genes with shadow (b) Genes without shadow (c) Genes with symmetric shadow

**Supplementary Figure 14:**

Species are shown on the x-axis and biological pathways on the y-axis. Color intensity, ranging from off-white to pink, indicates enrichment level, with off-white denoting no enrichment and darker pink shades indicating higher enrichment.

(a) Gene set enrichment analysis using all genes with shadows and the default background.

(b) Gene set enrichment analysis using genes with no shadow and the default background

(c) Gene set enrichment analysis using genes with symmetric shadows and no background.

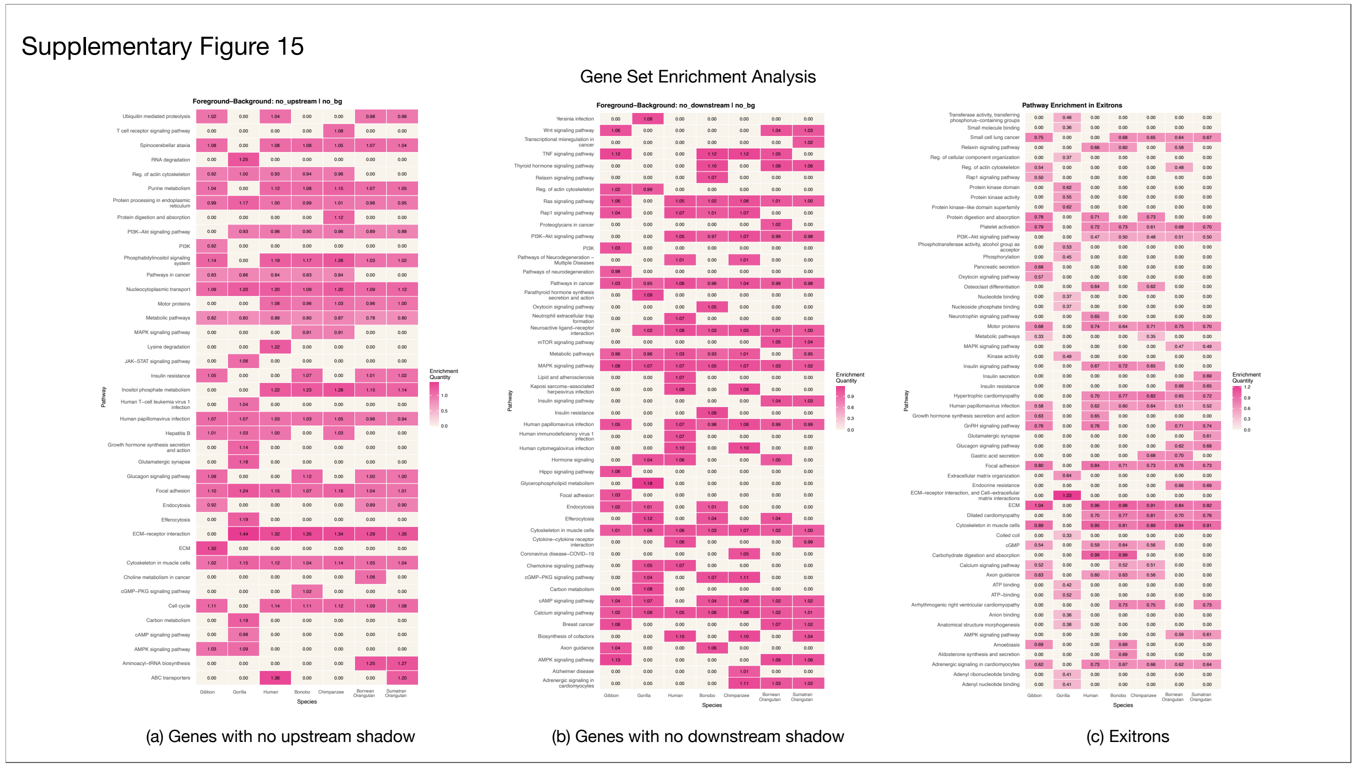


**Supplementary Figure 15:**
Species are shown on the x-axis and biological pathways on the y-axis. Color intensity, ranging from off-white to pink, indicates enrichment level, with off-white denoting no enrichment and darker pink shades indicating higher enrichment.
(a) Gene set enrichment analysis using all genes with no upstream shadow and default background
(b) Gene set enrichment analysis using all genes with no downstream shadow and default background
(c) Gene set enrichment analysis using all genes with exitrons and default background

Supplementary Figure 16

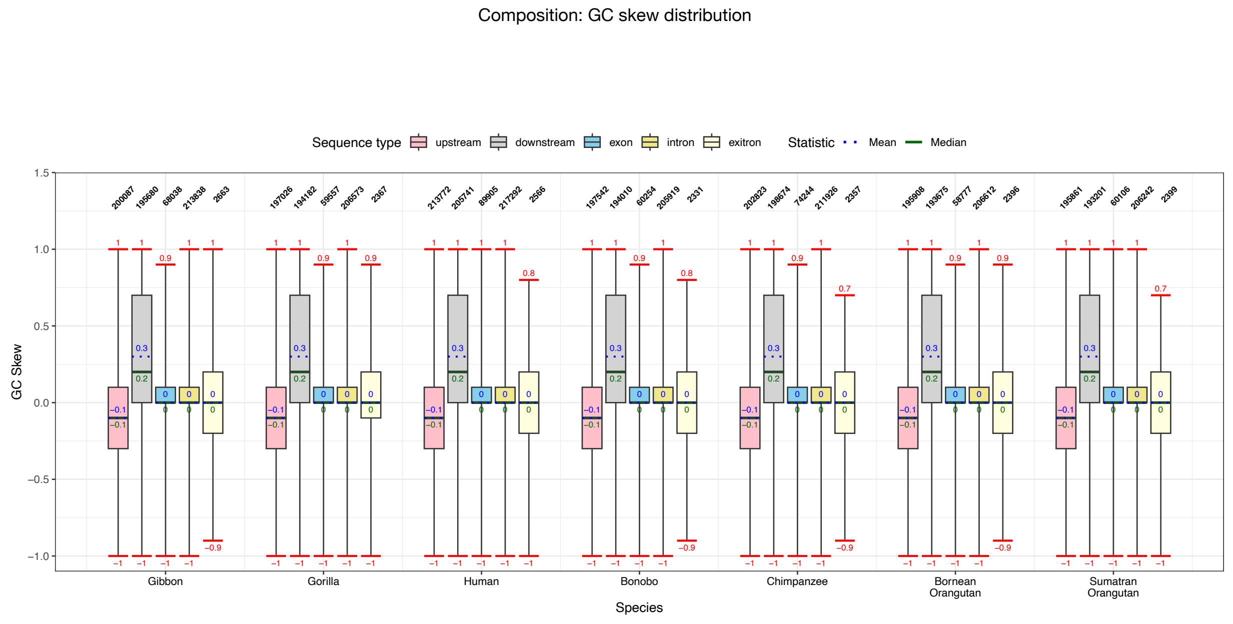


**Supplementary Figure 16:**
The x-axis indicates species and the y-axis indicates GC skew. For each species, boxplots show the skew distribution for five sequence types: upstream shadows (pink), downstream shadows (light grey), exons (blue), introns (yellow), and exitrons (cream). Whisker caps, indicated in red, mark the minimum and maximum values. Solid green and dotted blue lines denote the median and mean, respectively. Maximum and mean skew values are annotated above the corresponding lines, and minimum and mean values below. Total sequence counts for each boxplot are indicated above the boxes.

Supplementary Figure 17

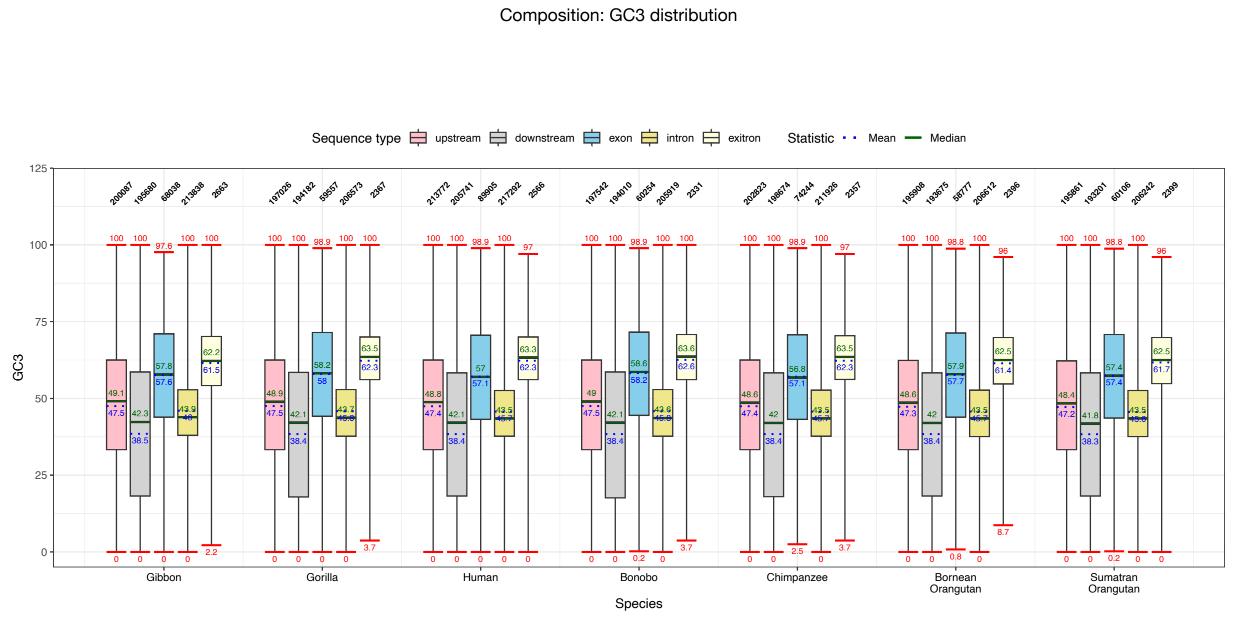


**Supplementary Figure 17:**
The x-axis indicates species and the y-axis indicates GC3. For each species, boxplots show the GC3 skew distribution for five sequence types: upstream shadows (pink), downstream shadows (light grey), exons (blue), introns (yellow), and exitrons (cream). Whisker caps, indicated in red, mark the minimum and maximum values. Solid green and dotted blue lines denote the median and mean, respectively. Maximum and mean GC3 values are annotated above the corresponding lines, and minimum and mean values below. Total sequence counts for each boxplot are indicated above the boxes.

Supplementary Figure 18

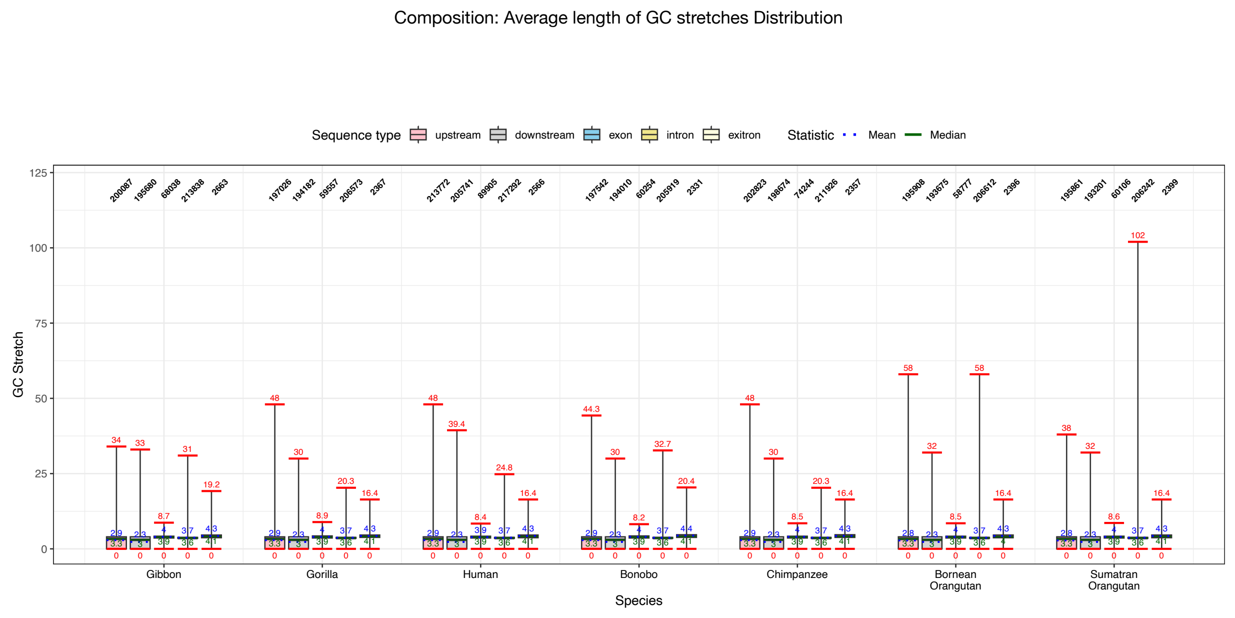


**Supplementary Figure 18:**
The x-axis indicates species and the y-axis indicates GC stretch. For each species, boxplots show the stretch distribution for five sequence types: upstream shadows (pink), downstream shadows (light grey), exons (blue), introns (yellow), and exitrons (cream). Whisker caps, indicated in red, mark the minimum and maximum values. Solid green and dotted blue lines denote the median and mean, respectively. Maximum and mean stretch values are annotated above the corresponding lines, and minimum and mean values below. Total sequence counts for each boxplot are indicated above the boxes.

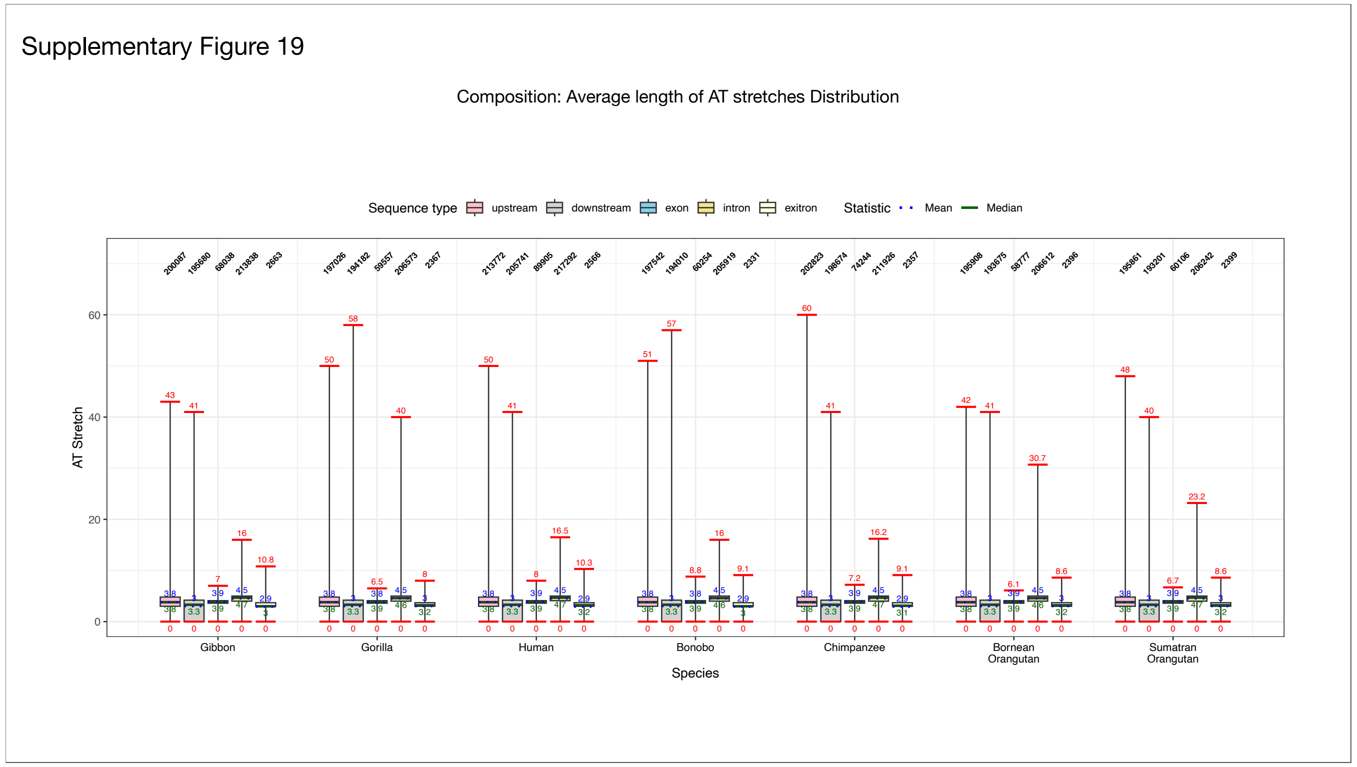


**Supplementary Figure 19:**
The x-axis indicates species and the y-axis indicates AT stretch. For each species, boxplots show the stretch distribution for five sequence types: upstream shadows (pink), downstream shadows (light grey), exons (blue), introns (yellow), and exitrons (cream). Whisker caps, indicated in red, mark the minimum and maximum values. Solid green and dotted blue lines denote the median and mean, respectively. Maximum and mean stretch values are annotated above the corresponding lines, and minimum and mean values below. Total sequence counts for each boxplot are indicated above the boxes.

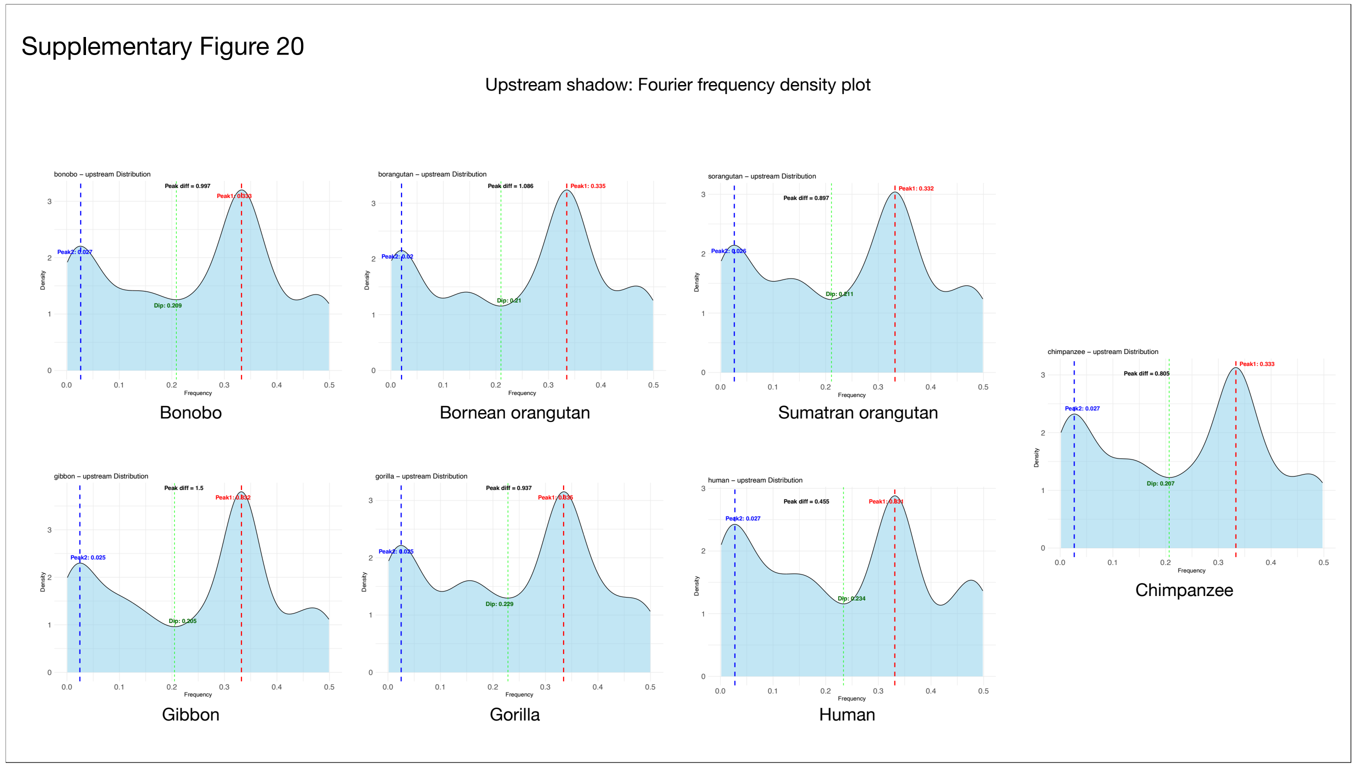


**Supplementary Figure 20:**

**Fourier frequency distribution of upstream shadows**

The plots show kernel density estimates (KDE) of frequency distributions of upstream exon shadows derived from discrete Fourier analysis across seven primate species. The x-axis represents spectral frequencies, and the y-axis indicates density. Red and blue dotted vertical lines mark the two highest spectral peaks, while the green line denotes the minimum (dip) between them. The annotated peak difference represents the distance between the lowest and highest frequencies observed.

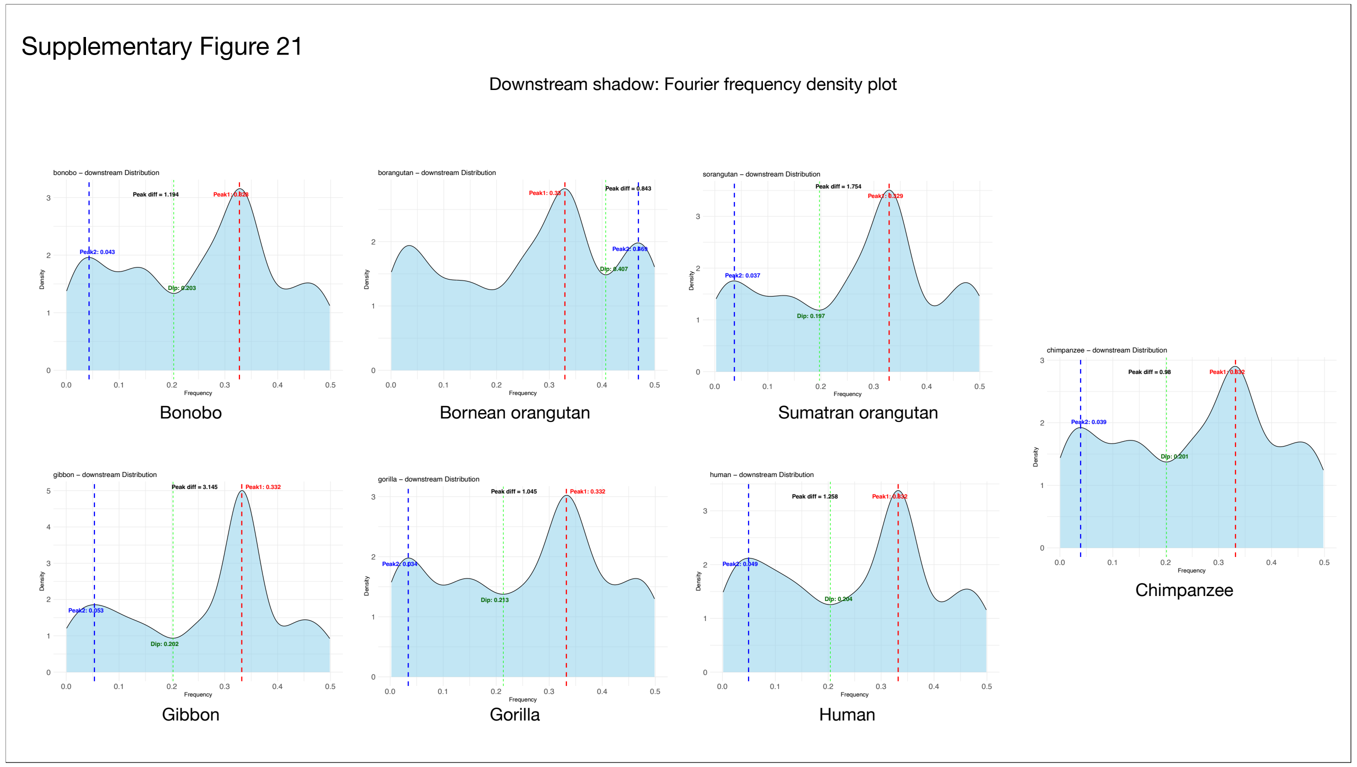


**Supplementary Figure 21:**

**Fourier frequency distribution of downstream shadows**

The plots show kernel density estimates (KDE) of frequency distributions of downstream exon shadows derived from discrete Fourier analysis across seven primate species. The x-axis represents spectral frequencies, and the y-axis indicates density. Red and blue dotted vertical lines mark the two highest spectral peaks, while the green line denotes the minimum (dip) between them. The annotated peak difference represents the distance between the lowest and highest frequencies observed.

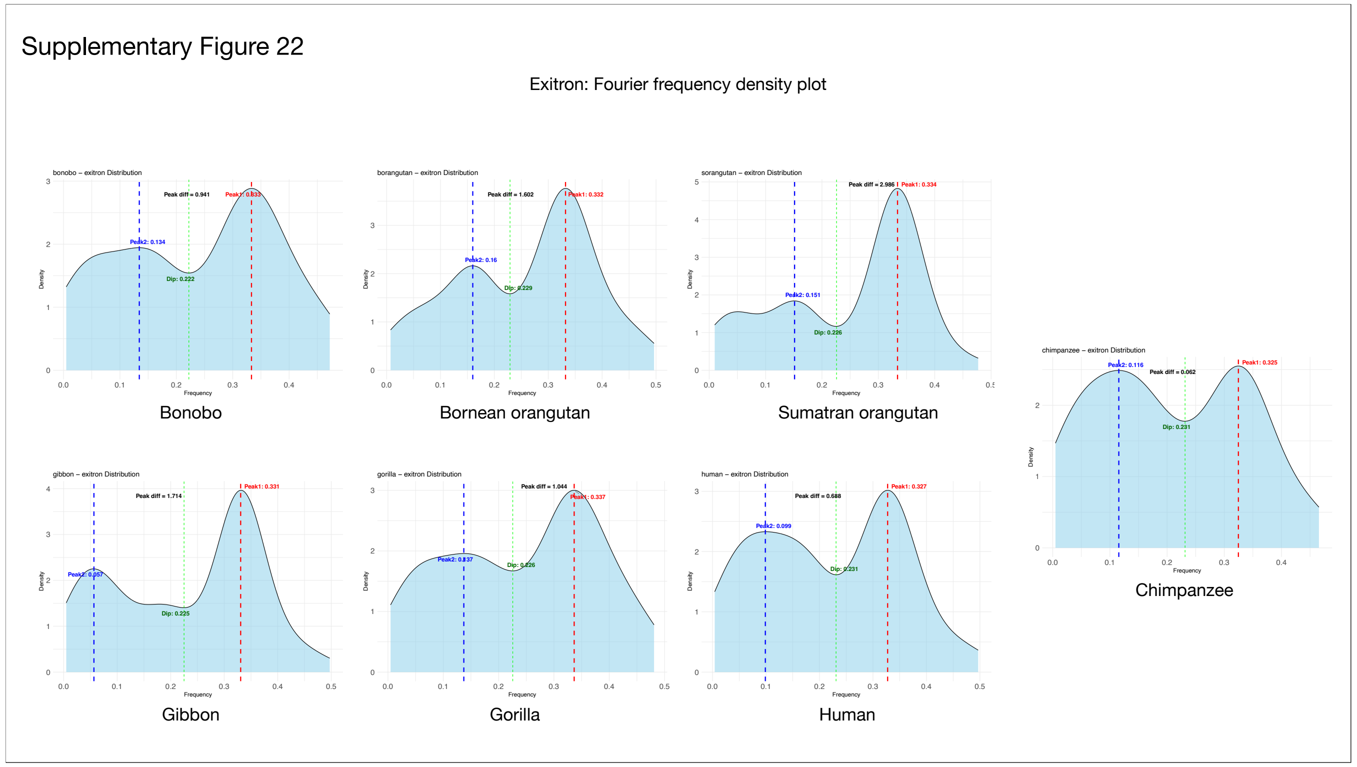


**Supplementary Figure 22:**

**Fourier frequency distribution of exitrons**

The plots show kernel density estimates (KDE) of frequency distributions of exitrons derived from discrete Fourier analysis across seven primate species. The x-axis represents spectral frequencies, and the y-axis indicates density. Red and blue dotted vertical lines mark the two highest spectral peaks, while the green line denotes the minimum (dip) between them. The annotated peak difference represents the distance between the lowest and highest frequencies observed.

Supplementary Figure 23

PCA + k-means at different length thresholds

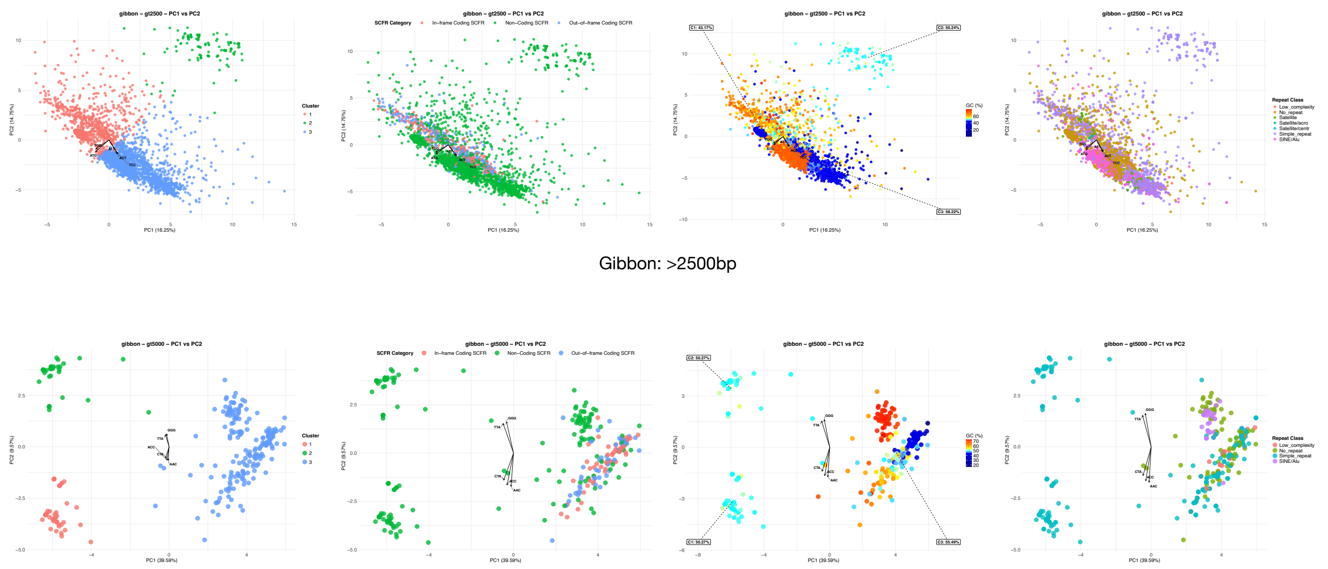


Gibbon: >2500bp

Gibbon: >5000bp

**Supplementary Figure 23:**

**PCA + KMeans at different length thresholds (Gibbon: >2500 bp and >5000 bp)**

Principal component analysis (PCA) was performed on SCFRs filtered by length, followed by K-means clustering. The x- and y-axes represent PC1 and PC2 scores, respectively.

**(a) Distribution by cluster:**

SCFRs are colored according to their assigned K-means cluster.

**(b) Distribution by SCFR category:**

SCFRs are colored by functional categories: red and blue denote in-frame and out-of-frame coding SCFRs, respectively and green denotes non-coding SCFRs.

**(c) Distribution by GC Content (%):**

SCFRs are colored according to GC content, with warmer colors indicating higher GC content and cooler colors indicating lower GC content.

**(d) Distribution by Repeat Class:**

SCFRs are colored based on their annotated repeat class.

Supplementary Figure 24

PCA + k-means at different length thresholds

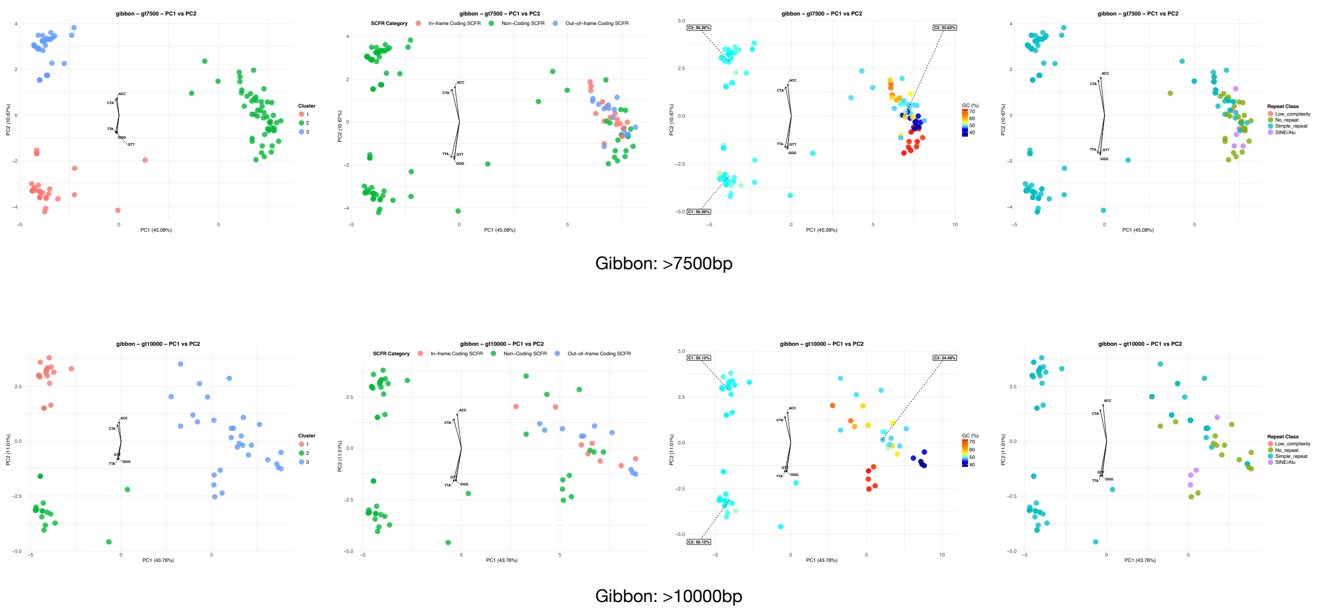


**Supplementary Figure 24:**

**PCA + KMeans at different length thresholds (Gibbon: >7500 bp and >10000 bp)**

Principal component analysis (PCA) was performed on SCFRs filtered by length, followed by K-means clustering. The x- and y-axes represent PC1 and PC2 scores, respectively.

**(a) Distribution by cluster:**

SCFRs are colored according to their assigned K-means cluster.

**(b) Distribution by SCFR category:**

SCFRs are colored by functional categories: red and blue denote in-frame and out-of-frame coding SCFRs, respectively and green denotes non-coding SCFRs.

**(c) Distribution by GC Content (%):**

SCFRs are colored according to GC content, with warmer colors indicating higher GC content and cooler colors indicating lower GC content.

**(d) Distribution by Repeat Class:**

SCFRs are colored based on their annotated repeat class.

Supplementary Figure 25

PCA + k-means at different length thresholds

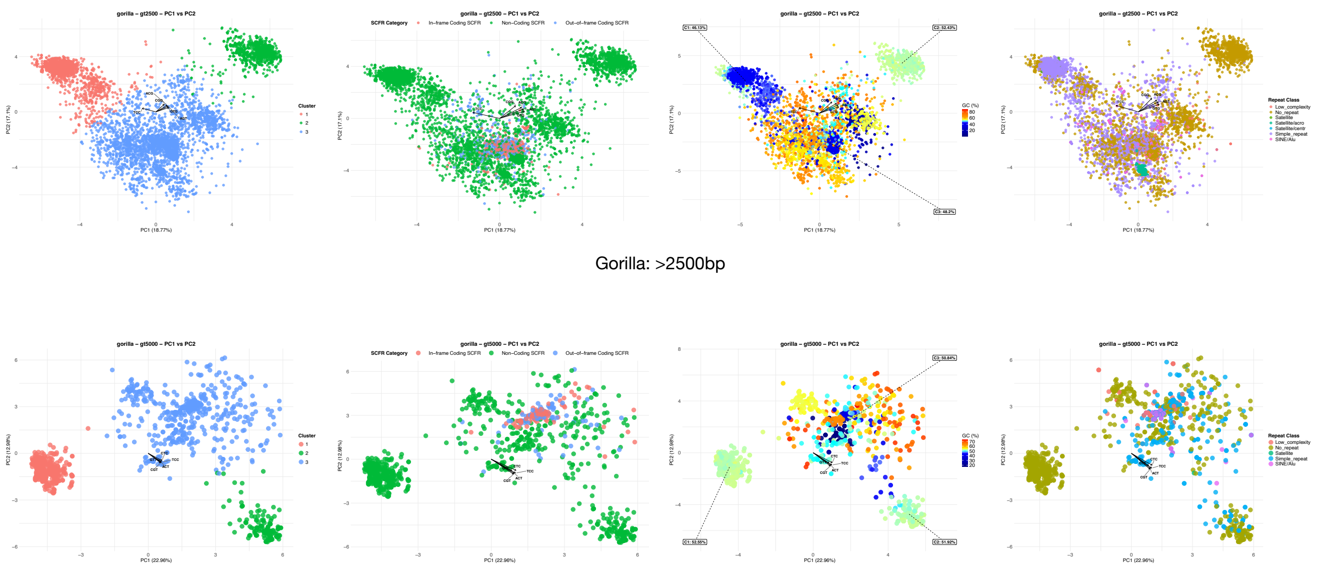


**Supplementary Figure 25:**

**PCA + KMeans at different length thresholds (Gorilla: >2500 bp and >5000 bp)**

Principal component analysis (PCA) was performed on SCFRs filtered by length, followed by K-means clustering. The x- and y-axes represent PC1 and PC2 scores, respectively.

**(a) Distribution by cluster:**

SCFRs are colored according to their assigned K-means cluster.

**(b) Distribution by SCFR category:**

SCFRs are colored by functional categories: red and blue denote in-frame and out-of-frame coding SCFRs, respectively and green denotes non-coding SCFRs.

**(c) Distribution by GC Content (%):**

SCFRs are colored according to GC content, with warmer colors indicating higher GC content and cooler colors indicating lower GC content.

**(d) Distribution by Repeat Class:**

SCFRs are colored based on their annotated repeat class.

Supplementary Figure 26

PCA + k-means at different length thresholds

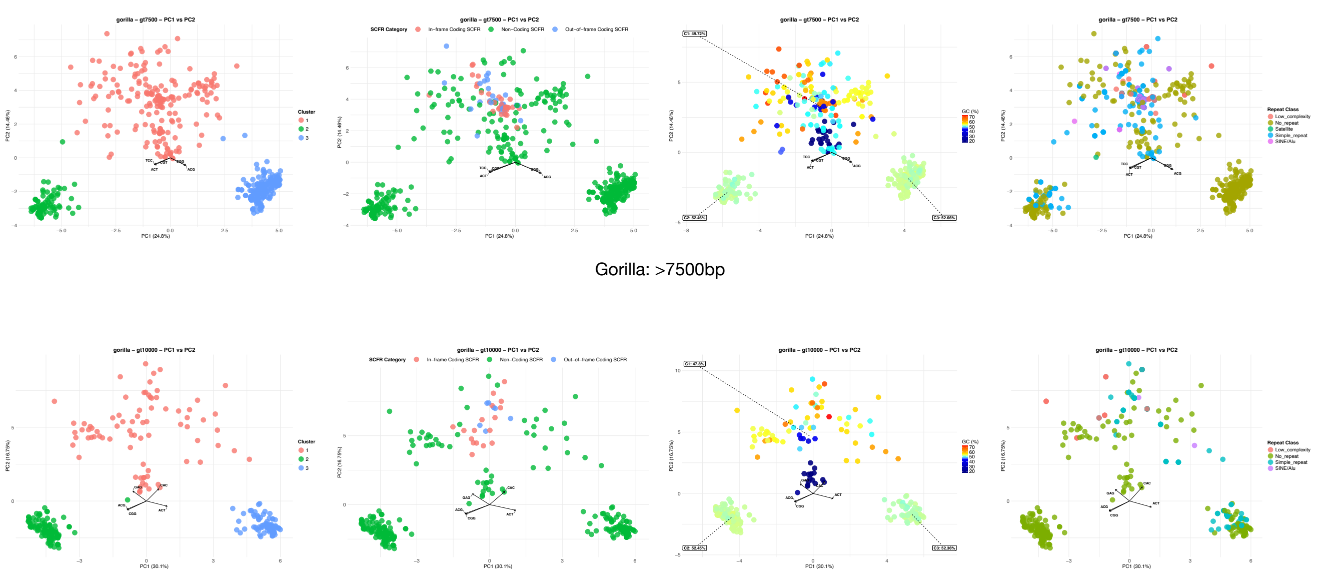


**Supplementary Figure 26:**
**PCA + KMeans at different length thresholds (Gorilla: >7500 bp and >10000 bp)**
Principal component analysis (PCA) was performed on SCFRs filtered by length, followed by K-means clustering. The x- and y-axes represent PC1 and PC2 scores, respectively.
**(a) Distribution by cluster:**
SCFRs are colored according to their assigned K-means cluster.
**(b) Distribution by SCFR category:**
SCFRs are colored by functional categories: red and blue denote in-frame and out-of-frame coding SCFRs, respectively and green denotes non-coding SCFRs.
**(c) Distribution by GC Content (%):**
SCFRs are colored according to GC content, with warmer colors indicating higher GC content and cooler colors indicating lower GC content.
**(d) Distribution by Repeat Class:**
SCFRs are colored based on their annotated repeat class.

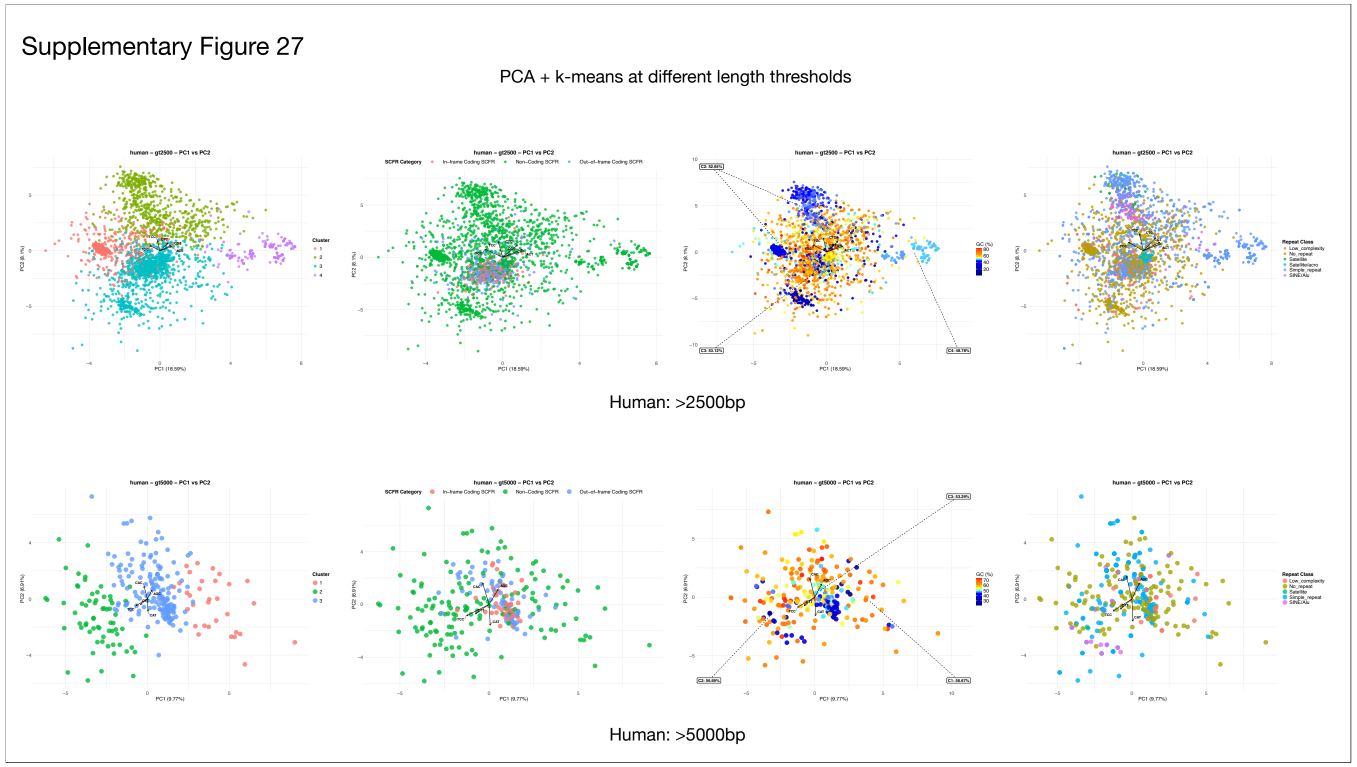


**Supplementary Figure 27:**
**PCA + KMeans at different length thresholds (Human: >2500 bp and >5000 bp)**
Principal component analysis (PCA) was performed on SCFRs filtered by length, followed by K-means clustering. The x- and y-axes represent PC1 and PC2 scores, respectively.
**(a) Distribution by cluster:**
SCFRs are colored according to their assigned K-means cluster.
**(b) Distribution by SCFR category:**
SCFRs are colored by functional categories: red and blue denote in-frame and out-of-frame coding SCFRs, respectively and green denotes non-coding SCFRs.
**(c) Distribution by GC Content (%):**
SCFRs are colored according to GC content, with warmer colors indicating higher GC content and cooler colors indicating lower GC content.
**(d) Distribution by Repeat Class:**
SCFRs are colored based on their annotated repeat class.

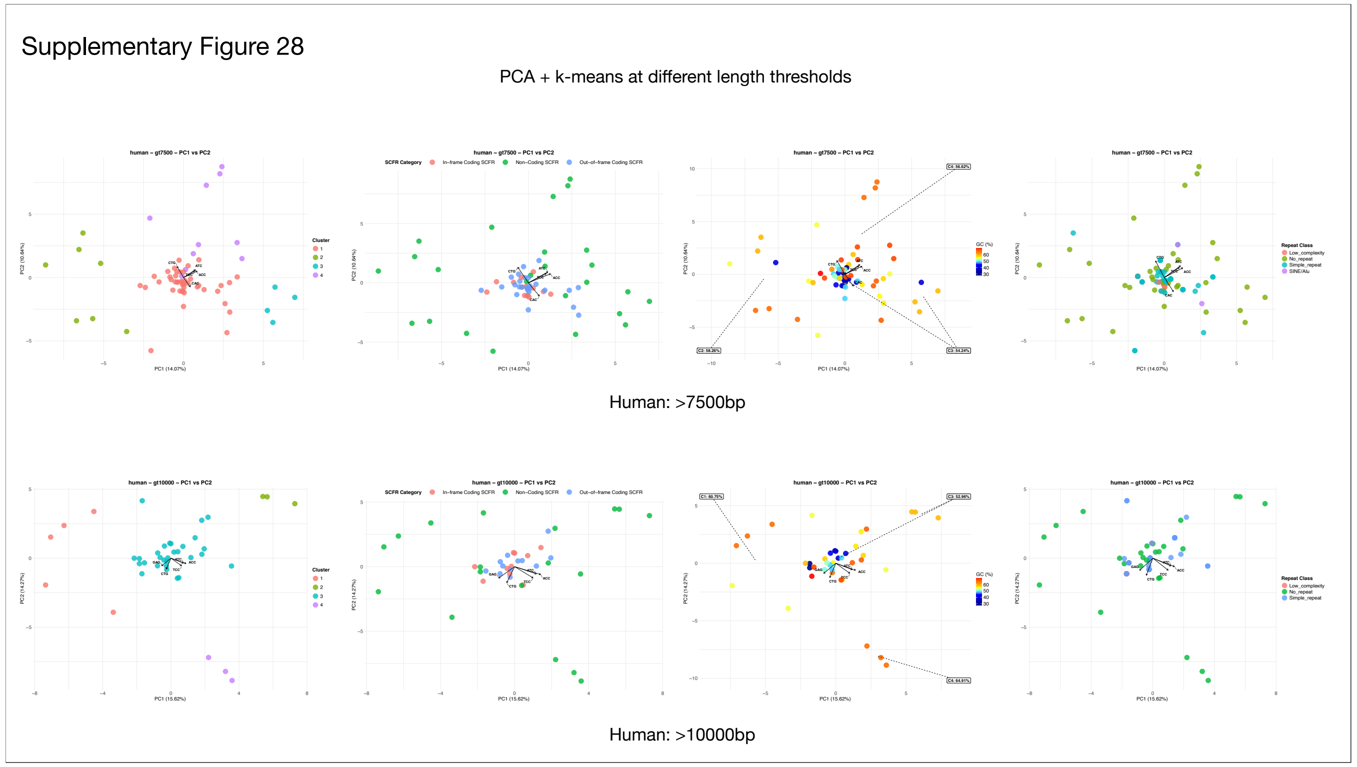


**Supplementary Figure 28:**
**PCA + KMeans at different length thresholds (Human: >7500 bp and >10000 bp)**
Principal component analysis (PCA) was performed on SCFRs filtered by length, followed by K-means clustering. The x- and y-axes represent PC1 and PC2 scores, respectively.
**(a) Distribution by cluster:**
SCFRs are colored according to their assigned K-means cluster.
**(b) Distribution by SCFR category:**
SCFRs are colored by functional categories: red and blue denote in-frame and out-of-frame coding SCFRs, respectively and green denotes non-coding SCFRs.
**(c) Distribution by GC Content (%):**
SCFRs are colored according to GC content, with warmer colors indicating higher GC content and cooler colors indicating lower GC content.
**(d) Distribution by Repeat Class:**
SCFRs are colored based on their annotated repeat class.

Supplementary Figure 29

PCA + k-means at different length thresholds

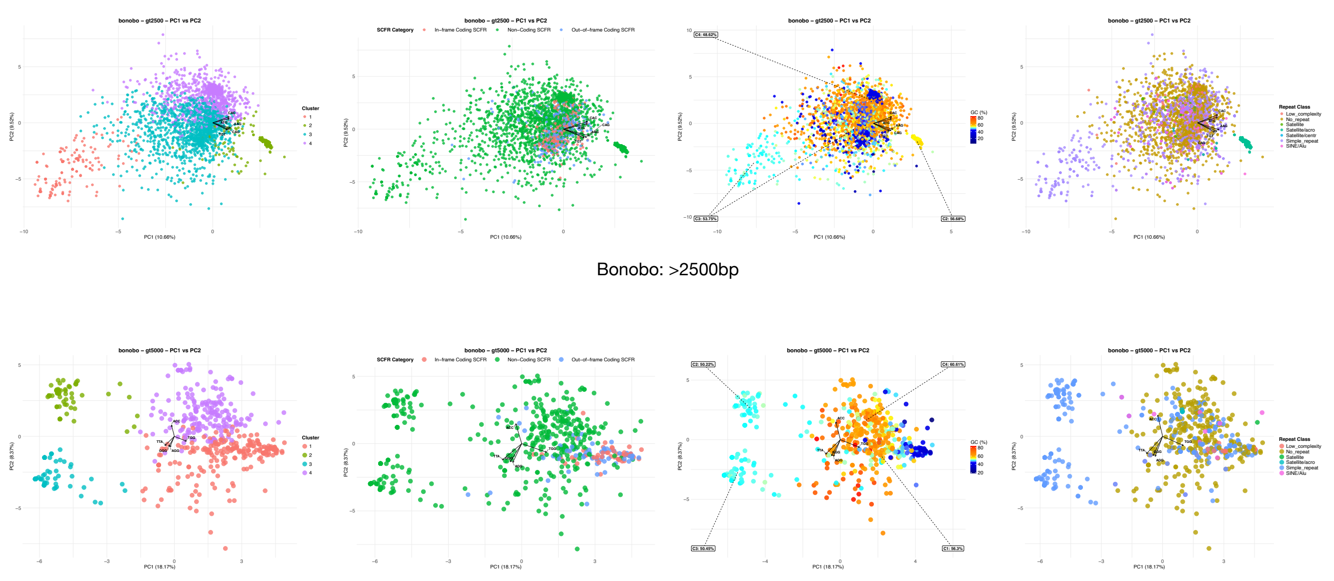


**Supplementary Figure 29:**

**PCA + KMeans at different length thresholds (Bonobo: >2500 bp and >5000 bp)**

Principal component analysis (PCA) was performed on SCFRs filtered by length, followed by K-means clustering. The x- and y-axes represent PC1 and PC2 scores, respectively.

**(a) Distribution by cluster:**

SCFRs are colored according to their assigned K-means cluster.

**(b) Distribution by SCFR category:**

SCFRs are colored by functional categories: red and blue denote in-frame and out-of-frame coding SCFRs, respectively and green denotes non-coding SCFRs.

**(c) Distribution by GC Content (%):**

SCFRs are colored according to GC content, with warmer colors indicating higher GC content and cooler colors indicating lower GC content.

**(d) Distribution by Repeat Class:**

SCFRs are colored based on their annotated repeat class.

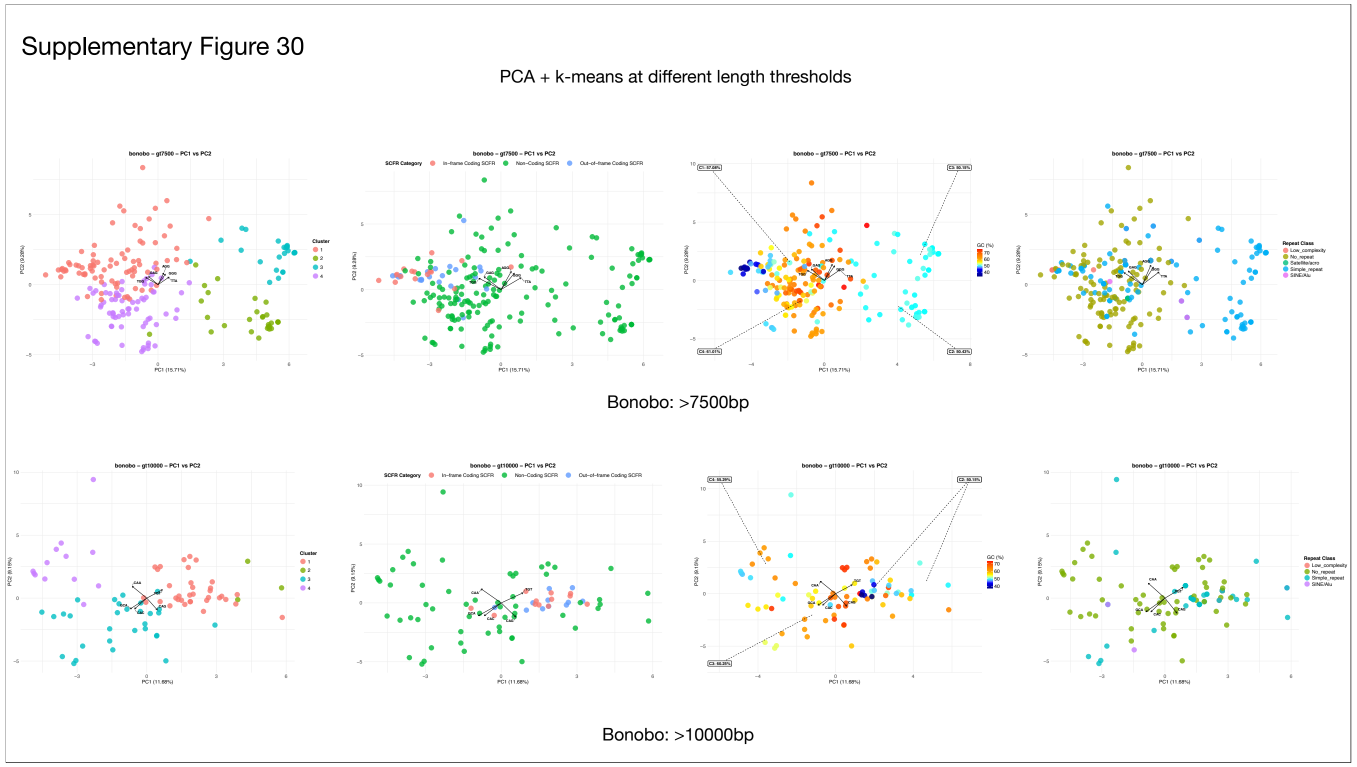


**Supplementary Figure 30:**

**PCA + KMeans at different length thresholds (Bonobo: >7500 bp and >10000 bp)**

Principal component analysis (PCA) was performed on SCFRs filtered by length, followed by K-means clustering. The x- and y-axes represent PC1 and PC2 scores, respectively.

**(a) Distribution by cluster:**

SCFRs are colored according to their assigned K-means cluster.

**(b) Distribution by SCFR category:**

SCFRs are colored by functional categories: red and blue denote in-frame and out-of-frame coding SCFRs, respectively and green denotes non-coding SCFRs.

**(c) Distribution by GC Content (%):**

SCFRs are colored according to GC content, with warmer colors indicating higher GC content and cooler colors indicating lower GC content.

**(d) Distribution by Repeat Class:**

SCFRs are colored based on their annotated repeat class.

Supplementary Figure 31

PCA + k-means at different length thresholds

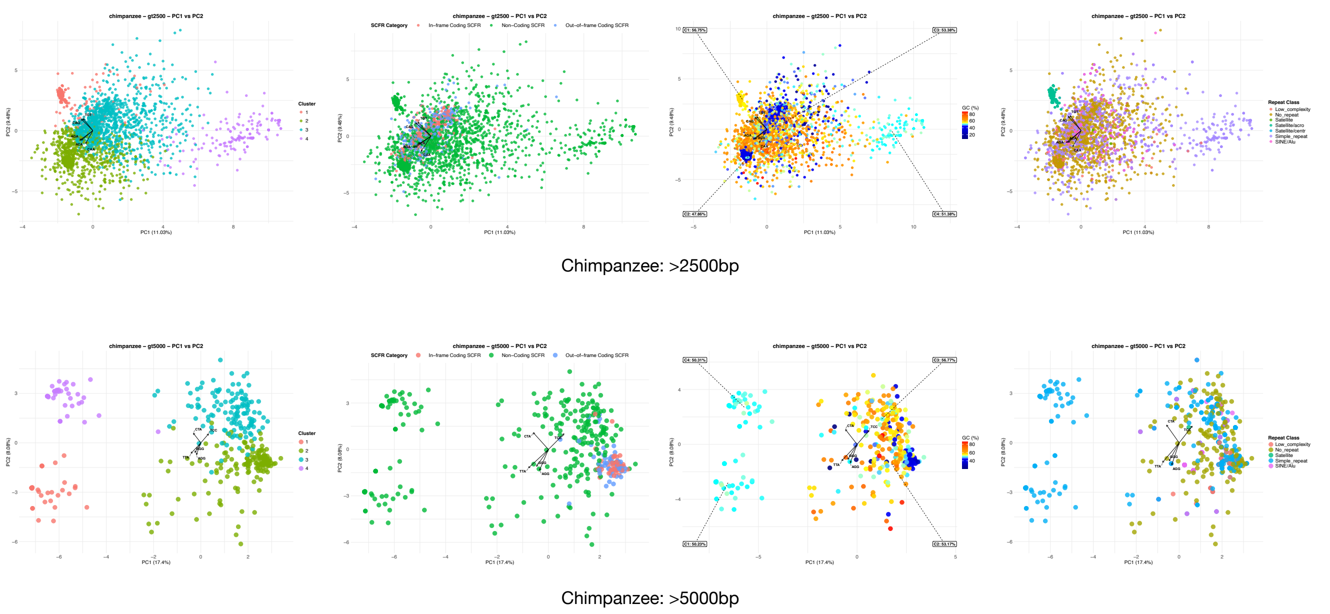


**Supplementary Figure 31:**

**PCA + KMeans at different length thresholds (Chimpanzee: >2500 bp and >5000 bp)**

Principal component analysis (PCA) was performed on SCFRs filtered by length, followed by K-means clustering. The x- and y-axes represent PC1 and PC2 scores, respectively.

**(a) Distribution by cluster:**

SCFRs are colored according to their assigned K-means cluster.

**(b) Distribution by SCFR category:**

SCFRs are colored by functional categories: red and blue denote in-frame and out-of-frame coding SCFRs, respectively and green denotes non-coding SCFRs.

**(c) Distribution by GC Content (%):**

SCFRs are colored according to GC content, with warmer colors indicating higher GC content and cooler colors indicating lower GC content.

**(d) Distribution by Repeat Class:**

SCFRs are colored based on their annotated repeat class.

### Supplementary Figure 32

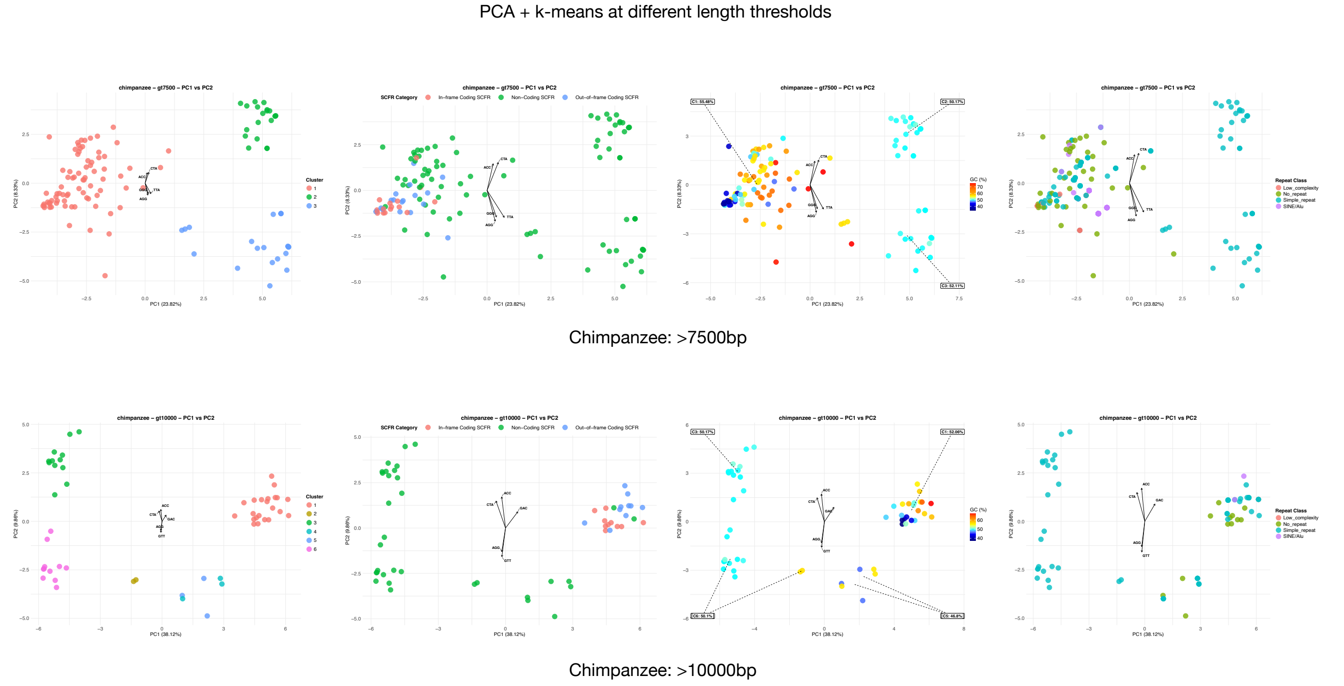


**Supplementary Figure 32:**

**PCA + KMeans at different length thresholds (Chimpanzee: >7500 bp and >10000 bp)**

Principal component analysis (PCA) was performed on SCFRs filtered by length, followed by K-means clustering. The x- and y-axes represent PC1 and PC2 scores, respectively.

**(a) Distribution by cluster:**

SCFRs are colored according to their assigned K-means cluster.

**(b) Distribution by SCFR category:**

SCFRs are colored by functional categories: red and blue denote in-frame and out-of-frame coding SCFRs, respectively and green denotes non-coding SCFRs.

**(c) Distribution by GC Content (%):**

SCFRs are colored according to GC content, with warmer colors indicating higher GC content and cooler colors indicating lower GC content.

**(d) Distribution by Repeat Class:**

SCFRs are colored based on their annotated repeat class.

Supplementary Figure 33

PCA + k-means at different length thresholds

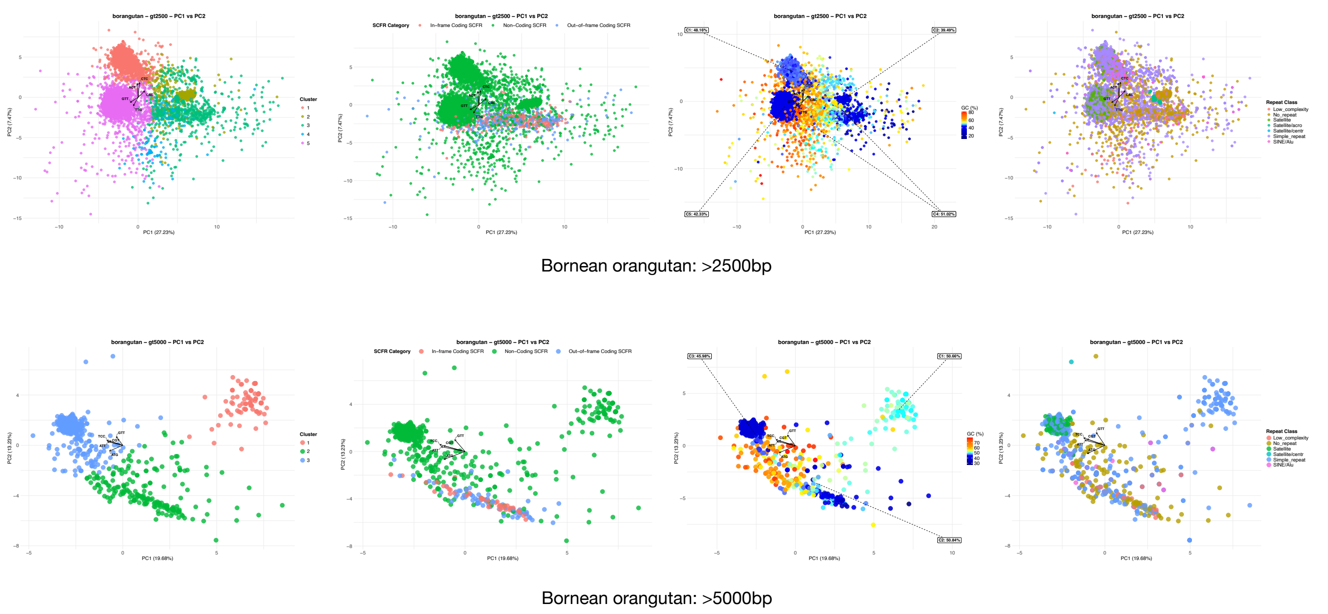


**Supplementary Figure 33:**

**PCA + KMeans at different length thresholds (Bornean orangutan: >2500 bp and >5000 bp)**

Principal component analysis (PCA) was performed on SCFRs filtered by length, followed by K-means clustering. The x- and y-axes represent PC1 and PC2 scores, respectively.

**(a) Distribution by cluster:**

SCFRs are colored according to their assigned K-means cluster.

**(b) Distribution by SCFR category:**

SCFRs are colored by functional categories: red and blue denote in-frame and out-of-frame coding SCFRs, respectively and green denotes non-coding SCFRs.

**(c) Distribution by GC Content (%):**

SCFRs are colored according to GC content, with warmer colors indicating higher GC content and cooler colors indicating lower GC content.

**(d) Distribution by Repeat Class:**

SCFRs are colored based on their annotated repeat class.

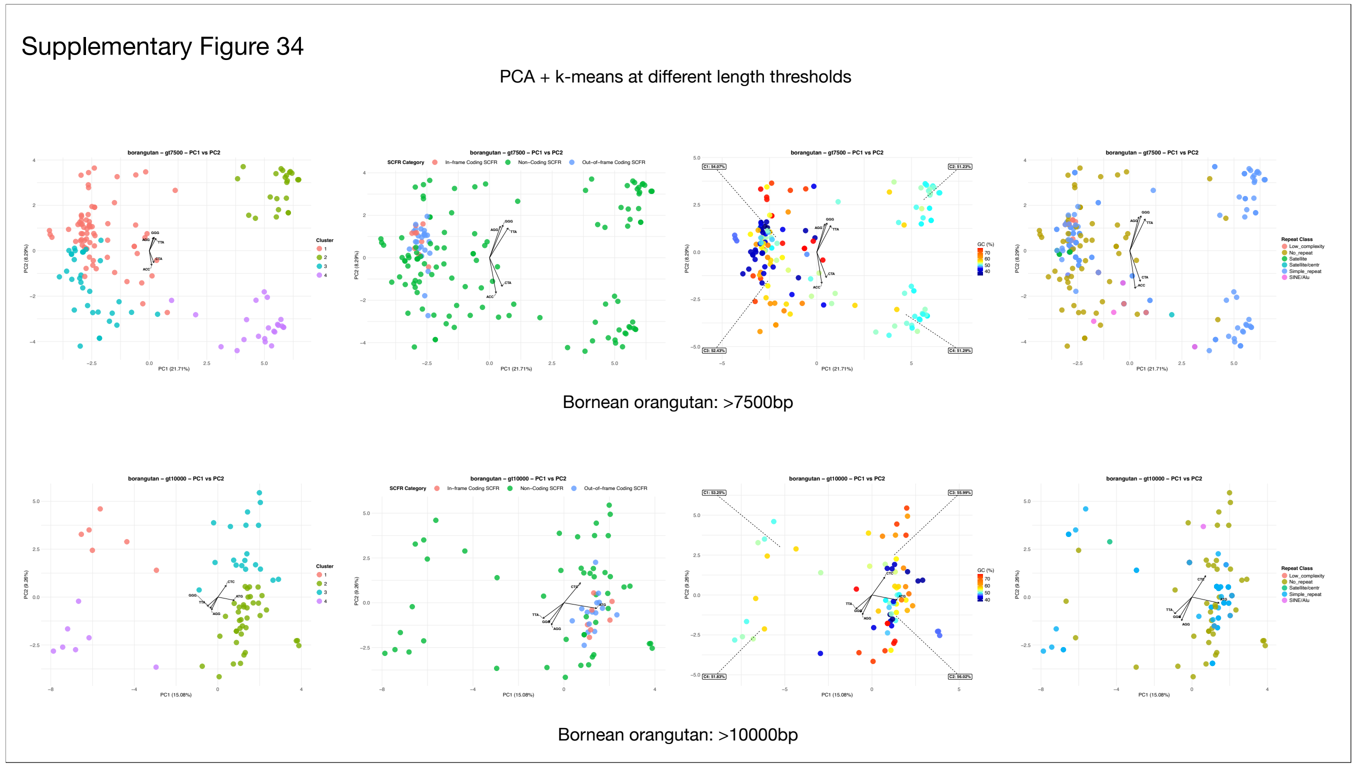


**Supplementary Figure 34:**

**PCA + KMeans at different length thresholds (Bornean orangutan: >7500 bp and >10000 bp)**

Principal component analysis (PCA) was performed on SCFRs filtered by length, followed by K-means clustering. The x- and y-axes represent PC1 and PC2 scores, respectively.

**(a) Distribution by cluster:**

SCFRs are colored according to their assigned K-means cluster.

**(b) Distribution by SCFR category:**

SCFRs are colored by functional categories: red and blue denote in-frame and out-of-frame coding SCFRs, respectively and green denotes non-coding SCFRs.

**(c) Distribution by GC Content (%):**

SCFRs are colored according to GC content, with warmer colors indicating higher GC content and cooler colors indicating lower GC content.

**(d) Distribution by Repeat Class:**

SCFRs are colored based on their annotated repeat class.

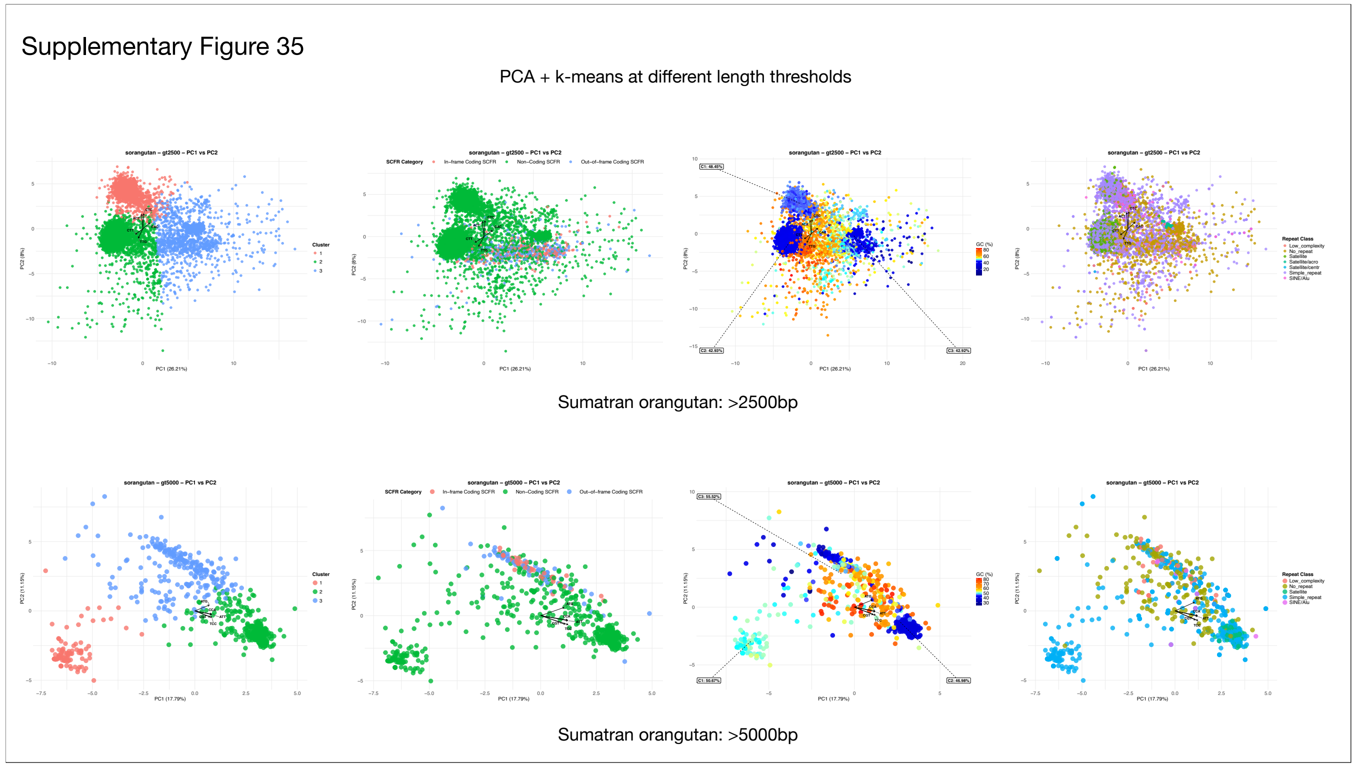


**Supplementary Figure 35:**

**PCA + KMeans at different length thresholds (Sumatran orangutan: >2500 bp and >5000 bp)**

Principal component analysis (PCA) was performed on SCFRs filtered by length, followed by K-means clustering. The x- and y-axes represent PC1 and PC2 scores, respectively.

**(a) Distribution by cluster:**

SCFRs are colored according to their assigned K-means cluster.

**(b) Distribution by SCFR category:**

SCFRs are colored by functional categories: red and blue denote in-frame and out-of-frame coding SCFRs, respectively and green denotes non-coding SCFRs.

**(c) Distribution by GC Content (%):**

SCFRs are colored according to GC content, with warmer colors indicating higher GC content and cooler colors indicating lower GC content.

**(d) Distribution by Repeat Class:**

SCFRs are colored based on their annotated repeat class.

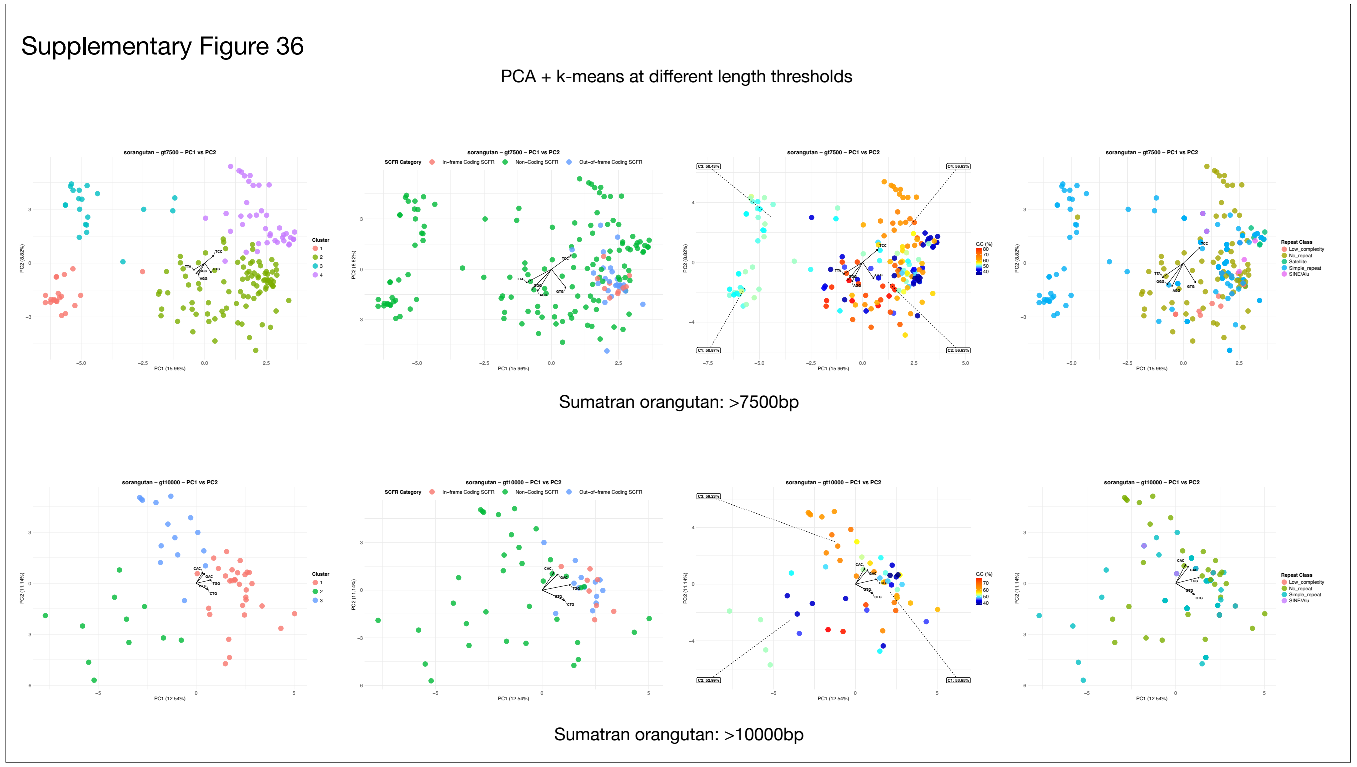


**Supplementary Figure 36:**

**PCA + KMeans at different length thresholds (Sumatran orangutan: >7500 bp and >10000 bp)**

Principal component analysis (PCA) was performed on SCFRs filtered by length, followed by K-means clustering. The x- and y-axes represent PC1 and PC2 scores, respectively.

**(a) Distribution by cluster:**

SCFRs are colored according to their assigned K-means cluster.

**(b) Distribution by SCFR category:**

SCFRs are colored by functional categories: red and blue denote in-frame and out-of-frame coding SCFRs, respectively and green denotes non-coding SCFRs.

**(c) Distribution by GC Content (%):**

SCFRs are colored according to GC content, with warmer colors indicating higher GC content and cooler colors indicating lower GC content.

**(d) Distribution by Repeat Class:**

SCFRs are colored based on their annotated repeat class.

Supplementary Figure 37

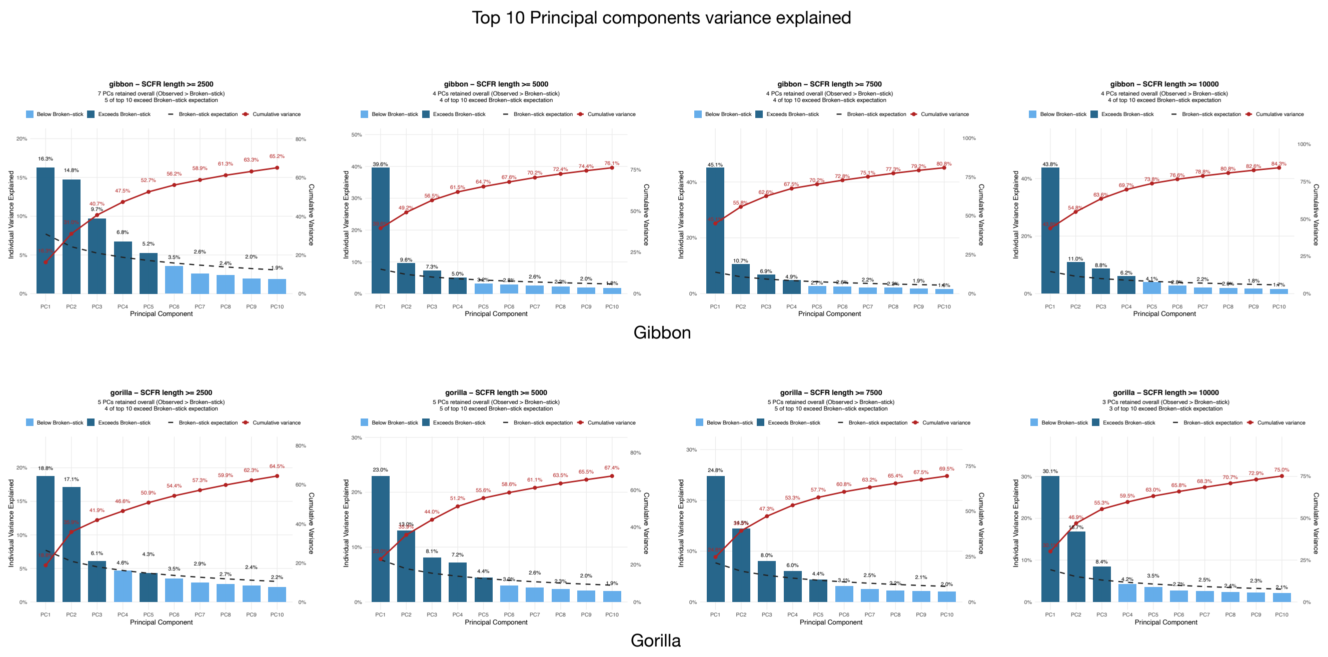


**Supplementary Figure 37:**

**Variance explained by the Top 10 Principal Components**

The plots show the variance explained by the top 10 principal components (PCs). The X-axis denotes the PCs, while the Y-axis indicates the percentage of variance explained by each. Bars in dark blue exceed the expectations from the Broken-stick test, whereas light blue bars fall below it. The black dotted line represents the Broken-stick expectation for each PC, and the red line with circles shows the cumulative variance explained. The total number of retained PCs and the count of top 10 PCs exceeding the Broken-stick expectation are annotated above each plot. For each species, separate figures are provided for SCFRs with minimum lengths of 2,500, 5,000, 7,500, and 10,000.

(a) Variance explained by Top 10 Principal Components – Gibbon

(b) Variance explained by Top 10 Principal Components – Gorilla

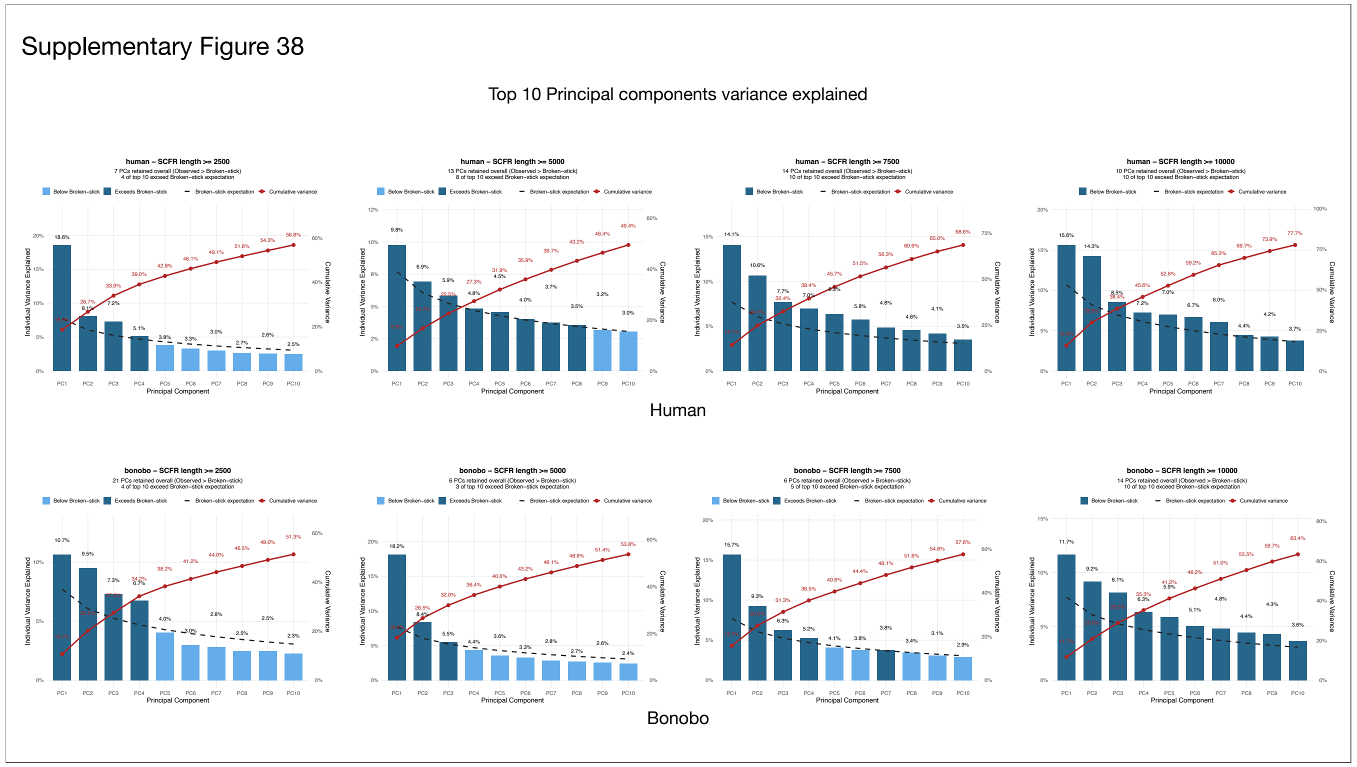


**Supplementary Figure 38:**

**Variance explained by the Top 10 Principal Components**

The plots show the variance explained by the top 10 principal components (PCs). The X-axis denotes the PCs, while the Y-axis indicates the percentage of variance explained by each. Bars in dark blue exceed the expectations from the Broken-stick test, whereas light blue bars fall below it. The black dotted line represents the Broken-stick expectation for each PC, and the red line with circles shows the cumulative variance explained. The total number of retained PCs and the count of top 10 PCs exceeding the Broken-stick expectation are annotated above each plot. For each species, separate figures are provided for SCFRs with minimum lengths of 2,500, 5,000, 7,500, and 10,000.

(a) Variance explained by Top 10 Principal Components – Human

(b) Variance explained by Top 10 Principal Components – Bonobo

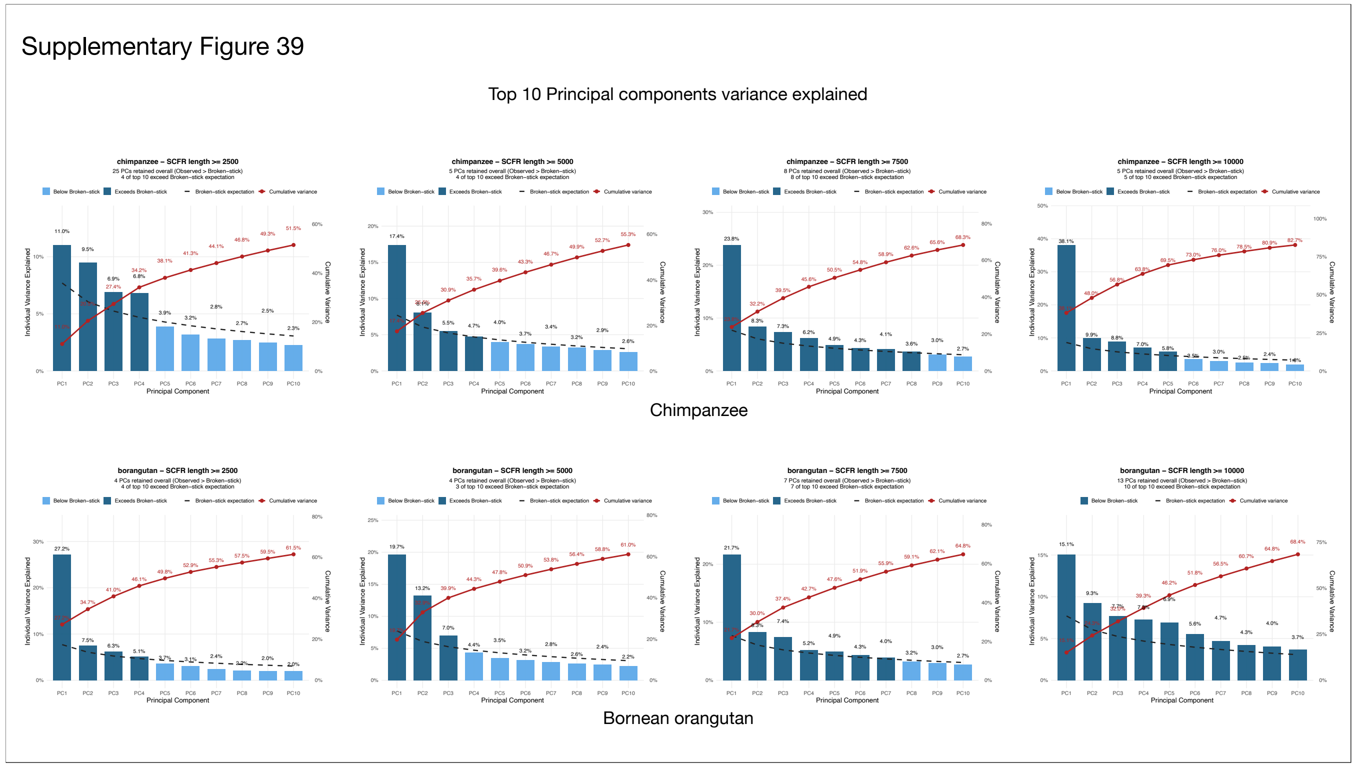


**Supplementary Figure 39:**

**Variance explained by the Top 10 Principal Components**

The plots show the variance explained by the top 10 principal components (PCs). The X-axis denotes the PCs, while the Y-axis indicates the percentage of variance explained by each. Bars in dark blue exceed the expectations from the Broken-stick test, whereas light blue bars fall below it. The black dotted line represents the Broken-stick expectation for each PC, and the red line with circles shows the cumulative variance explained. The total number of retained PCs and the count of top 10 PCs exceeding the Broken-stick expectation are annotated above each plot. For each species, separate figures are provided for SCFRs with minimum lengths of 2,500, 5,000, 7,500, and 10,000.

(a) Variance explained by Top 10 Principal Components – Chimpanzee

(b) Variance explained by Top 10 Principal Components – Bornean Orangutan

Supplementary Figure 40

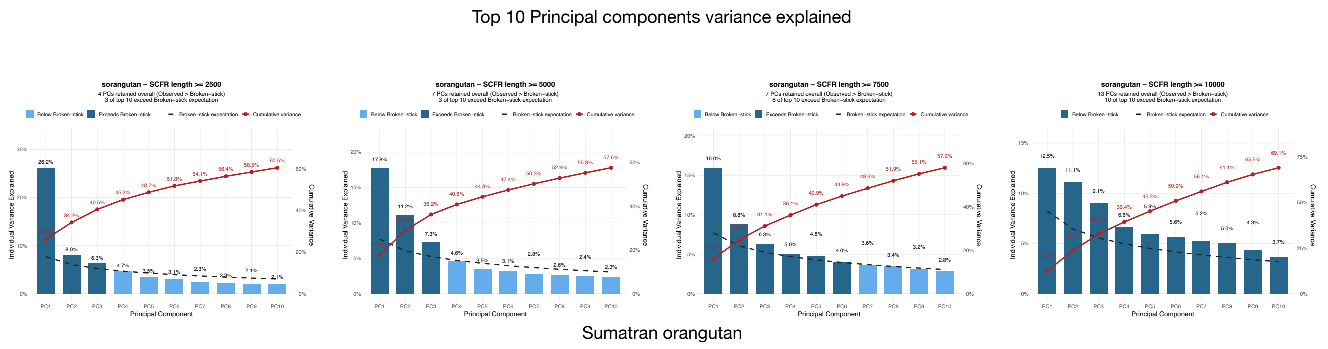


**Supplementary Figure 40:**

**Variance explained by the Top 10 Principal Components**

The plots show the variance explained by the top 10 principal components (PCs). The X-axis denotes the PCs, while the Y-axis indicates the percentage of variance explained by each. Bars in dark blue exceed the expectations from the Broken-stick test, whereas light blue bars fall below it. The black dotted line represents the Broken-stick expectation for each PC, and the red line with circles shows the cumulative variance explained. The total number of retained PCs and the count of top 10 PCs exceeding the Broken-stick expectation are annotated above each plot. For each species, separate figures are provided for SCFRs with minimum lengths of 2,500, 5,000, 7,500, and 10,000.

(a) Variance explained by Top 10 Principal Components – Sumatran Orangutan

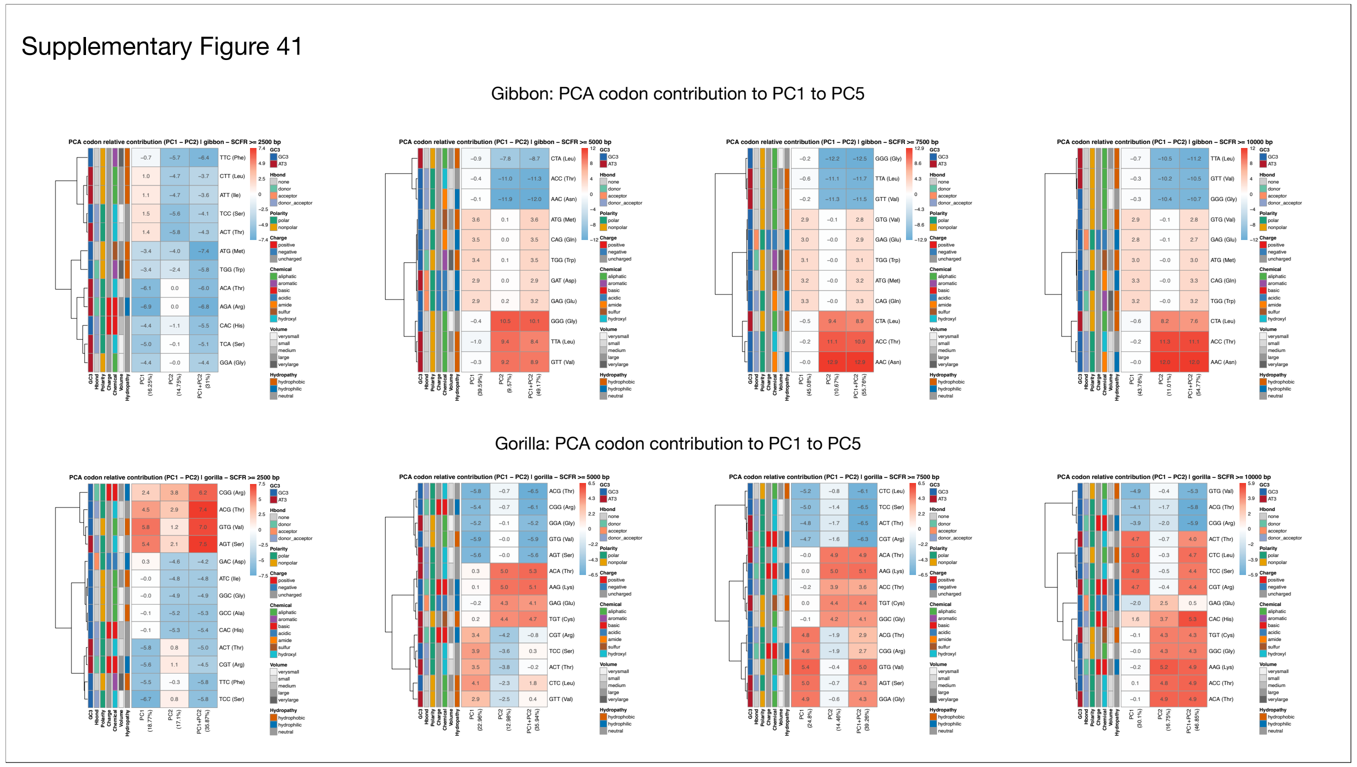


**Supplementary Figure 41:**

**Relative Codon Contributions by Top 2 PCs**

The plots show the relative contribution of each codon to PC1, PC2, and the combined PC1+PC2. The X-axis represents the PCs and their explained variance, while the Y-axis lists codons with their corresponding amino acids. Adjacent bars indicate amino acid properties, including GC3 content, hydrogen bonding, polarity, charge, chemical class, volume, and hydropathy. Cell colors in the heatmap reflect contributions to variance, with cooler tones indicating negative contributions and warmer tones positive contributions. For each species, separate figures are provided for SCFRs with minimum lengths of 2,500, 5,000, 7,500, and 10,000.

(a) Relative Codon Contribution – Top 2 PCs – Gibbon

(b) Relative Codon Contribution – Top 2 PCs – Gorilla

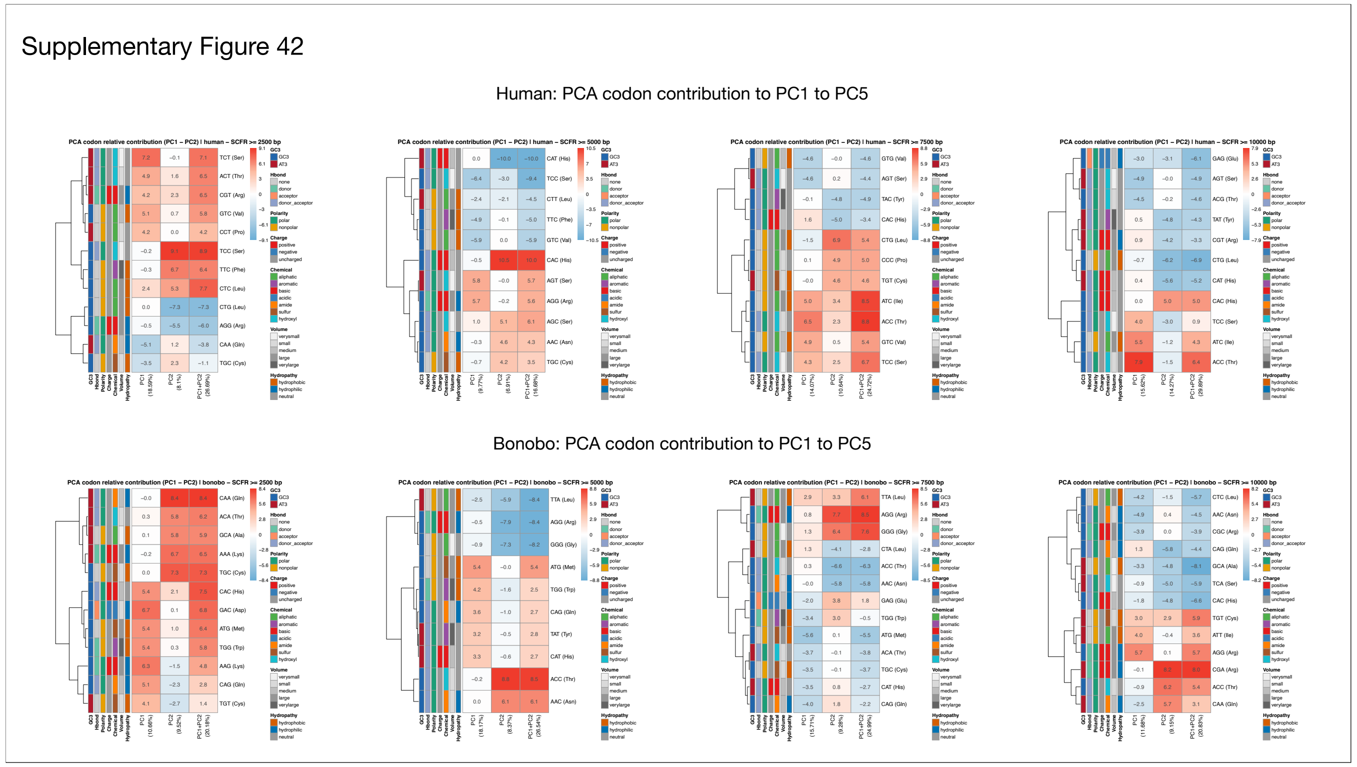


**Supplementary Figure 42:**

**Relative Codon Contributions by Top 2 PCs**

The plots show the relative contribution of each codon to PC1, PC2, and the combined PC1+PC2. The X-axis represents the PCs and their explained variance, while the Y-axis lists codons with their corresponding amino acids. Adjacent bars indicate amino acid properties, including GC3 content, hydrogen bonding, polarity, charge, chemical class, volume, and hydropathy. Cell colors in the heatmap reflect contributions to variance, with cooler tones indicating negative contributions and warmer tones positive contributions. For each species, separate figures are provided for SCFRs with minimum lengths of 2,500, 5,000, 7,500, and 10,000.

(a) Relative Codon Contribution – Top 2 PCs – Human

(b) Relative Codon Contribution – Top 2 PCs – Bonobo

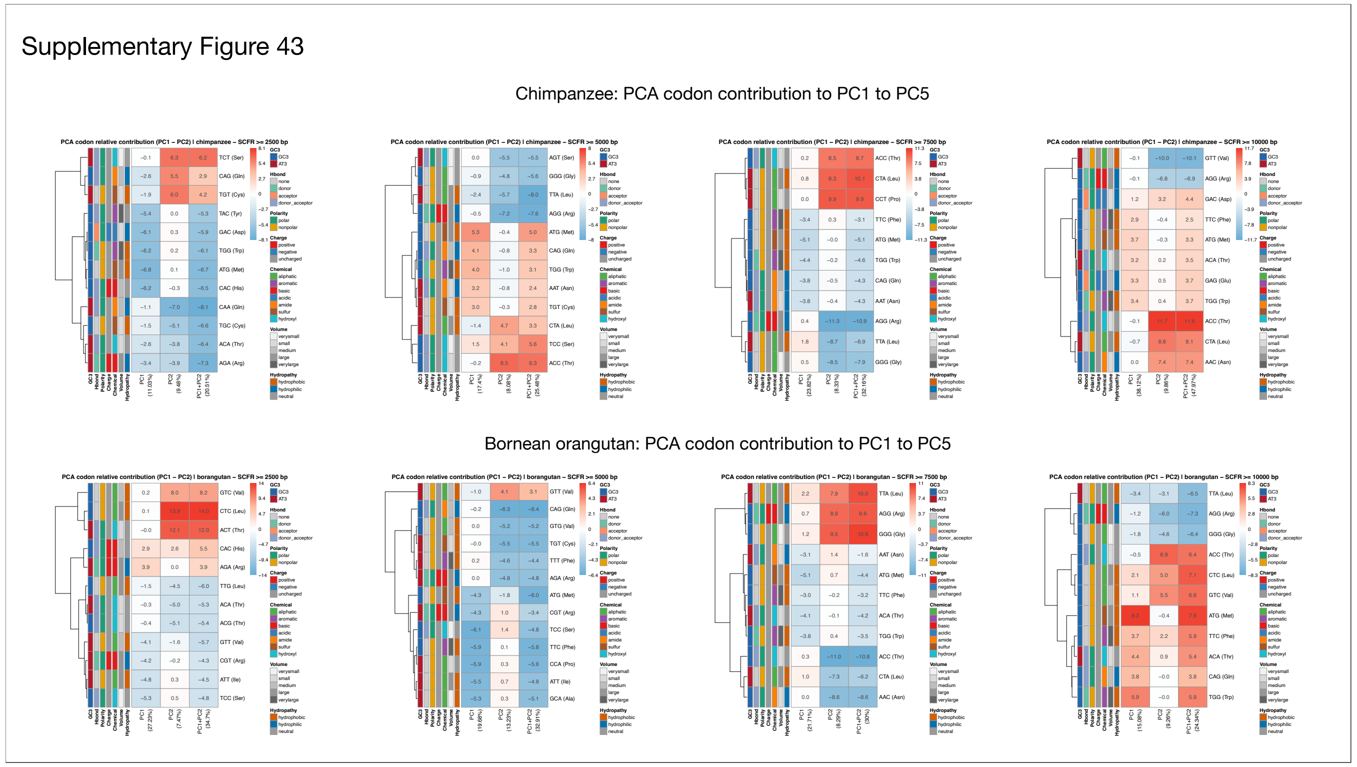


**Supplementary Figure 43:**

**Relative Codon Contributions by Top 2 PCs**

The plots show the relative contribution of each codon to PC1, PC2, and the combined PC1+PC2. The X-axis represents the PCs and their explained variance, while the Y-axis lists codons with their corresponding amino acids. Adjacent bars indicate amino acid properties, including GC3 content, hydrogen bonding, polarity, charge, chemical class, volume, and hydropathy. Cell colors in the heatmap reflect contributions to variance, with cooler tones indicating negative contributions and warmer tones positive contributions. For each species, separate figures are provided for SCFRs with minimum lengths of 2,500, 5,000, 7,500, and 10,000.

(a) Relative Codon Contribution – Top 2 PCs – Chimpanzee

(b) Relative Codon Contribution – Top 2 PCs – Bornean Orangutan

Supplementary Figure 44

Sumatran orangutan: PCA codon contribution to PC1 to PC5

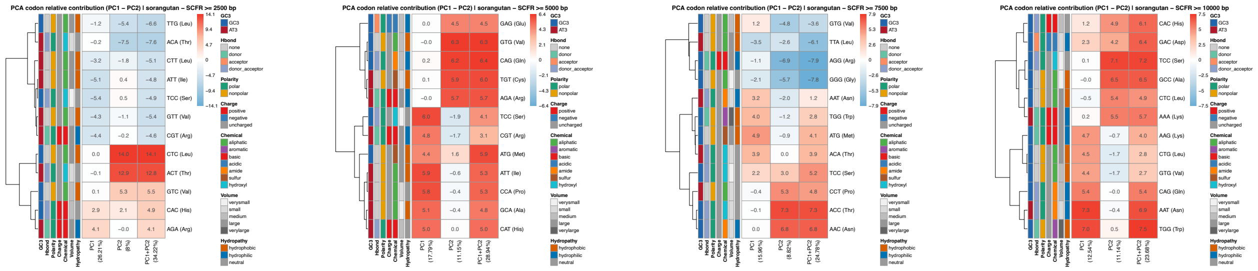

**Supplementary Figure 44:**

**Relative Codon Contributions by Top 2 PCs**

The plots show the relative contribution of each codon to PC1, PC2, and the combined PC1+PC2. The X-axis represents the PCs and their explained variance, while the Y-axis lists codons with their corresponding amino acids. Adjacent bars indicate amino acid properties, including GC3 content, hydrogen bonding, polarity, charge, chemical class, volume, and hydropathy. Cell colors in the heatmap reflect contributions to variance, with cooler tones indicating negative contributions and warmer tones positive contributions. For each species, separate figures are provided for SCFRs with minimum lengths of 2,500, 5,000, 7,500, and 10,000.

(a) Relative Codon Contribution – Top 2 PCs – Sumatran Orangutan

Supplementary Figure 45

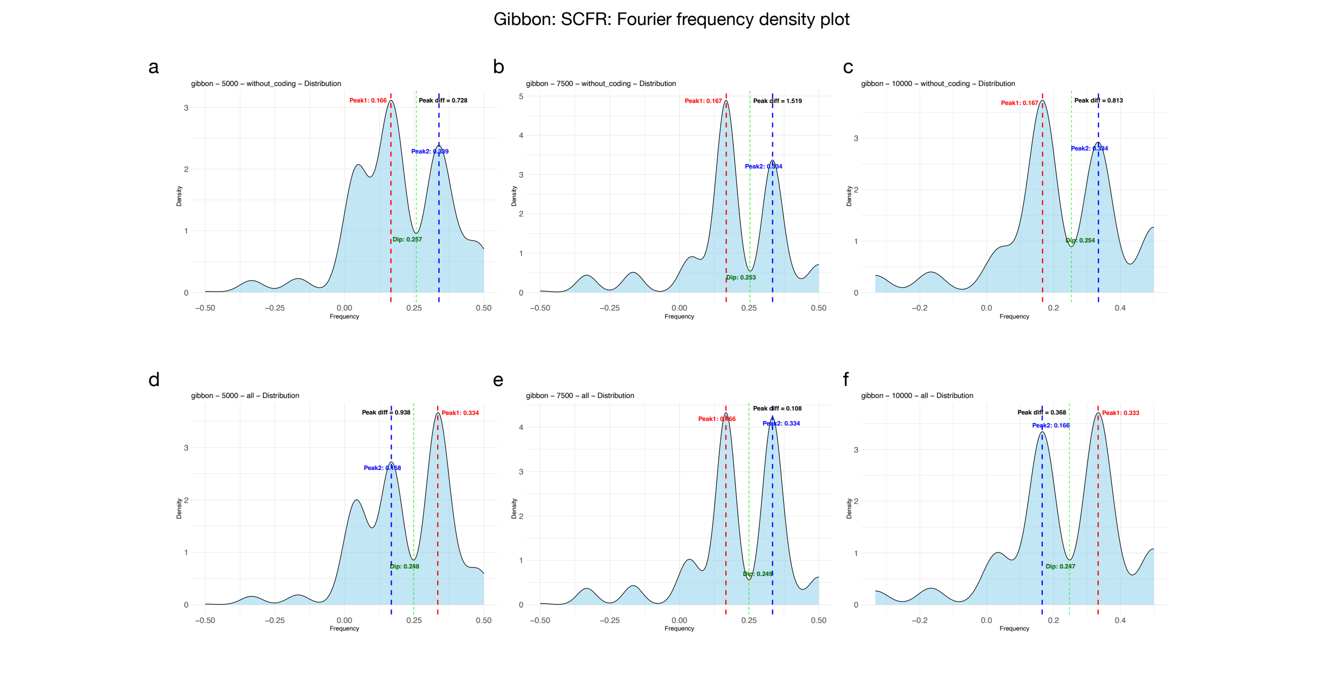


**Supplementary Figure 45:**

**Fourier frequency density plots of SCFRs at different length thresholds and categories in Gibbon.**

Kernel density plots show the distribution of frequencies derived from Fourier spectra of SCFRs. The x-axis represents frequency and the y-axis represents density. Red and blue dotted vertical lines indicate the two highest spectral peaks, while the green line marks the intervening minimum (dip). The annotated peak difference denotes the distance between the lowest and highest frequencies observed.

(a) Non-coding SCFRs with a minimum length of 5000 bp.
(b) Non-coding SCFRs with a minimum length of 7500 bp.
(c) Non-coding SCFRs with a minimum length of 10000 bp.
(d) Combined coding and non-coding SCFRs with a minimum length of 5000 bp.
(e) Combined coding and non-coding SCFRs with a minimum length of 7500 bp.
(f) Combined coding and non-coding SCFRs with a minimum length of 10000 bp.

Supplementary Figure 46

Gorilla: SCFR: Fourier frequency density plot

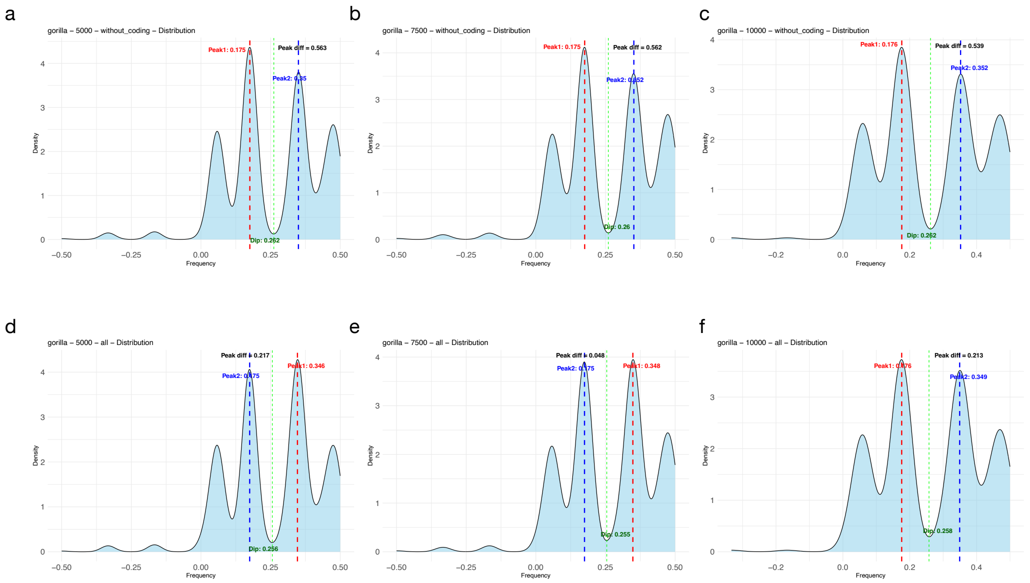


**Supplementary Figure 46:**

**Fourier frequency density plots of SCFRs at different length thresholds and categories in Gorilla.**

Kernel density plots show the distribution of frequencies derived from Fourier spectra of SCFRs. The x-axis represents frequency and the y-axis represents density. Red and blue dotted vertical lines indicate the two highest spectral peaks, while the green line marks the intervening minimum (dip). The annotated peak difference denotes the distance between the lowest and highest frequencies observed.

(a) Non-coding SCFRs with a minimum length of 5000 bp.
(b) Non-coding SCFRs with a minimum length of 7500 bp.
(c) Non-coding SCFRs with a minimum length of 10000 bp.
(d) Combined coding and non-coding SCFRs with a minimum length of 5000 bp.
(e) Combined coding and non-coding SCFRs with a minimum length of 7500 bp.
(f) Combined coding and non-coding SCFRs with a minimum length of 10000 bp.

Supplementary Figure 47

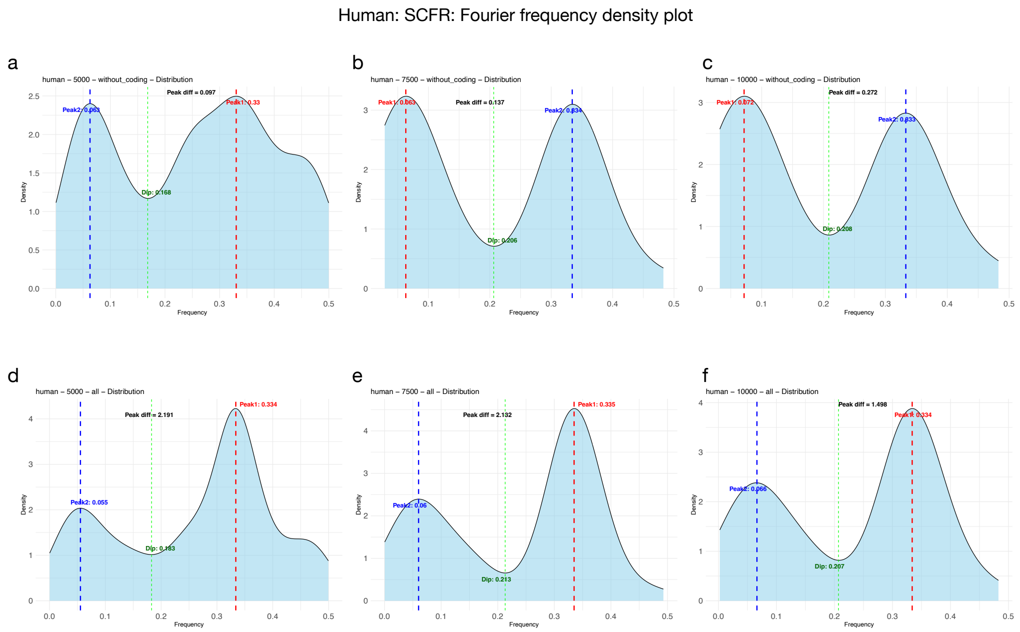


**Supplementary Figure 47:**

**Fourier frequency density plots of SCFRs at different length thresholds and categories in Humans.**

Kernel density plots show the distribution of frequencies derived from Fourier spectra of SCFRs. The x-axis represents frequency and the y-axis represents density. Red and blue dotted vertical lines indicate the two highest spectral peaks, while the green line marks the intervening minimum (dip). The annotated peak difference denotes the distance between the lowest and highest frequencies observed.

(a) Non-coding SCFRs with a minimum length of 5000 bp.
(b) Non-coding SCFRs with a minimum length of 7500 bp.
(c) Non-coding SCFRs with a minimum length of 10000 bp.
(d) Combined coding and non-coding SCFRs with a minimum length of 5000 bp.
(e) Combined coding and non-coding SCFRs with a minimum length of 7500 bp.
(f) Combined coding and non-coding SCFRs with a minimum length of 10000 bp.

Supplementary Figure 48

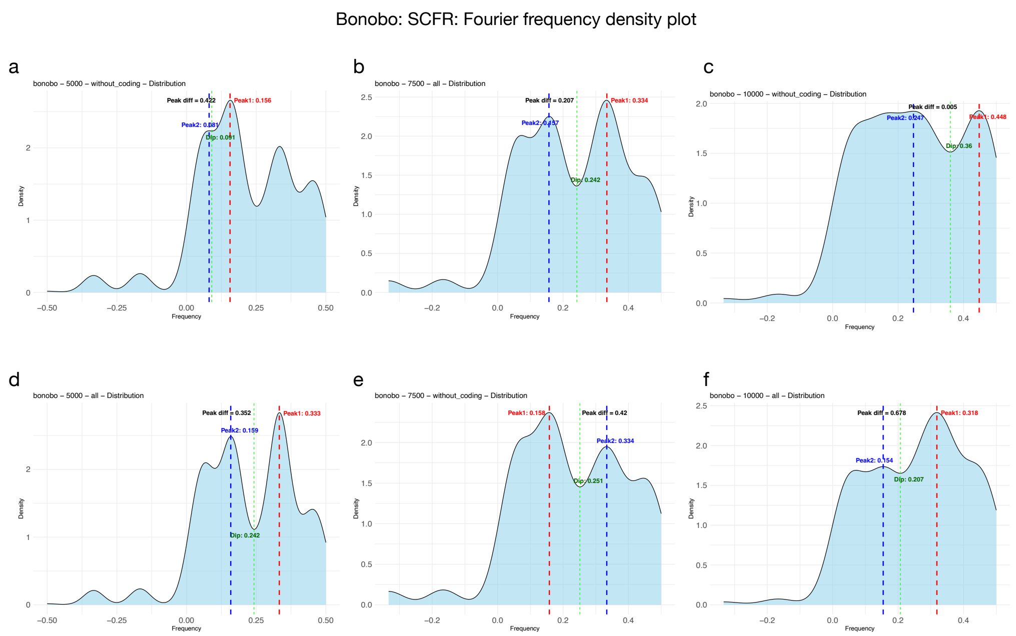


**Supplementary Figure 48:**

**Fourier frequency density plots of SCFRs at different length thresholds and categories in Bonobo.**

Kernel density plots show the distribution of frequencies derived from Fourier spectra of SCFRs. The x-axis represents frequency and the y-axis represents density. Red and blue dotted vertical lines indicate the two highest spectral peaks, while the green line marks the intervening minimum (dip). The annotated peak difference denotes the distance between the lowest and highest frequencies observed.

(a) Non-coding SCFRs with a minimum length of 5000 bp.
(b) Non-coding SCFRs with a minimum length of 7500 bp.
(c) Non-coding SCFRs with a minimum length of 10000 bp.
(d) Combined coding and non-coding SCFRs with a minimum length of 5000 bp.
(e) Combined coding and non-coding SCFRs with a minimum length of 7500 bp.
(f) Combined coding and non-coding SCFRs with a minimum length of 10000 bp.

Supplementary Figure 49

Chimpanzee: SCFR: Fourier frequency density plot

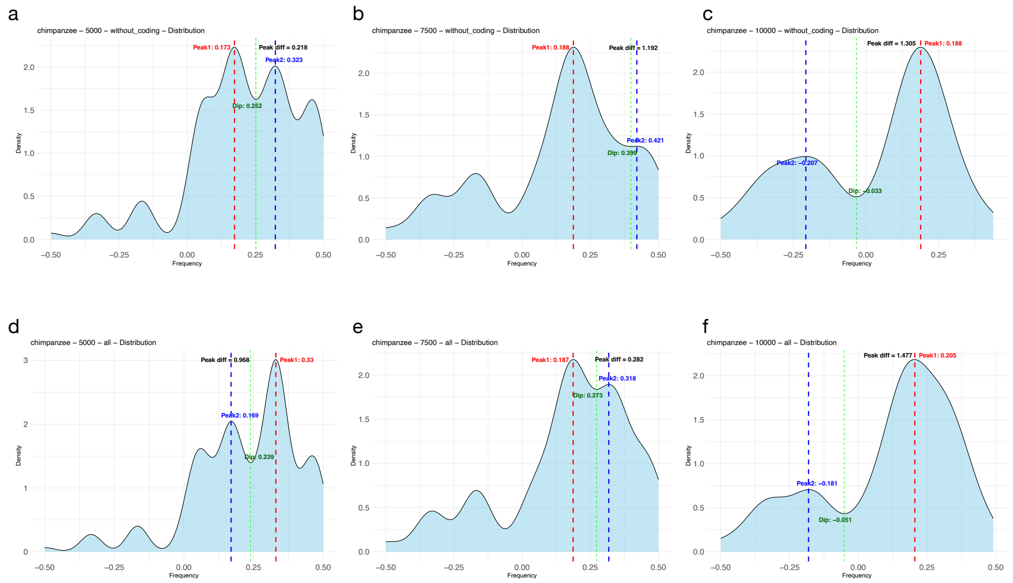


**Supplementary Figure 49:**

**Fourier frequency density plots of SCFRs at different length thresholds and categories in Chimpanzee.**

Kernel density plots show the distribution of frequencies derived from Fourier spectra of SCFRs. The x-axis represents frequency and the y-axis represents density. Red and blue dotted vertical lines indicate the two highest spectral peaks, while the green line marks the intervening minimum (dip). The annotated peak difference denotes the distance between the lowest and highest frequencies observed.

(a) Non-coding SCFRs with a minimum length of 5000 bp.
(b) Non-coding SCFRs with a minimum length of 7500 bp.
(c) Non-coding SCFRs with a minimum length of 10000 bp.
(d) Combined coding and non-coding SCFRs with a minimum length of 5000 bp.
(e) Combined coding and non-coding SCFRs with a minimum length of 7500 bp.
(f) Combined coding and non-coding SCFRs with a minimum length of 10000 bp.

Supplementary Figure 50

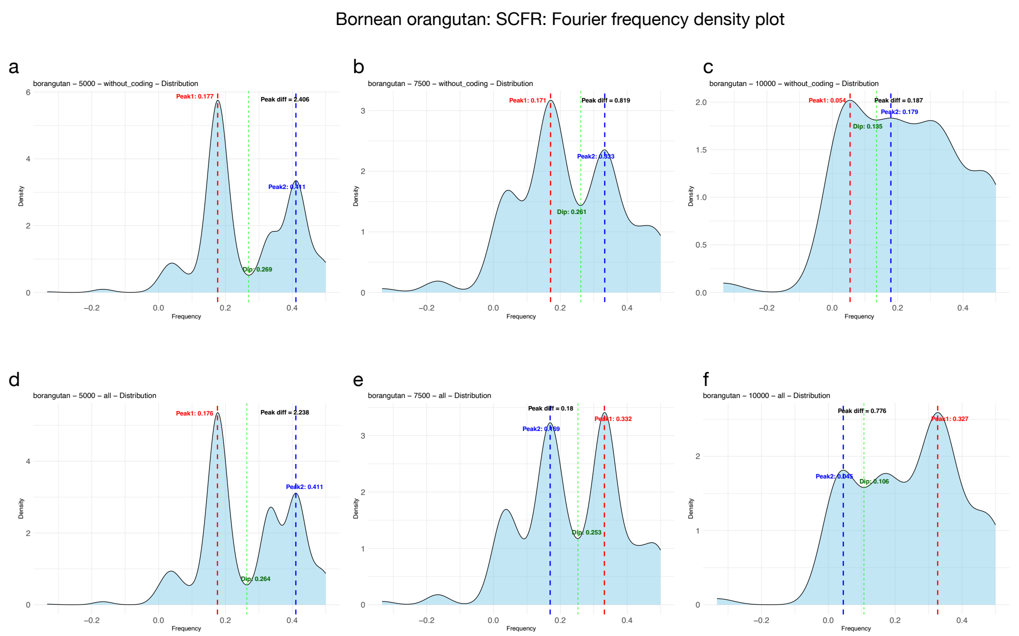


**Supplementary Figure 50:**

**Fourier frequency density plots of SCFRs at different length thresholds and categories in Bornean orangutan.**

Kernel density plots show the distribution of frequencies derived from Fourier spectra of SCFRs. The x-axis represents frequency and the y-axis represents density. Red and blue dotted vertical lines indicate the two highest spectral peaks, while the green line marks the intervening minimum (dip). The annotated peak difference denotes the distance between the lowest and highest frequencies observed.

(a) Non-coding SCFRs with a minimum length of 5000 bp.
(b) Non-coding SCFRs with a minimum length of 7500 bp.
(c) Non-coding SCFRs with a minimum length of 10000 bp.
(d) Combined coding and non-coding SCFRs with a minimum length of 5000 bp.
(e) Combined coding and non-coding SCFRs with a minimum length of 7500 bp.
(f) Combined coding and non-coding SCFRs with a minimum length of 10000 bp.

Supplementary Figure 51

Sumatran orangutan: SCFR: Fourier frequency density plot

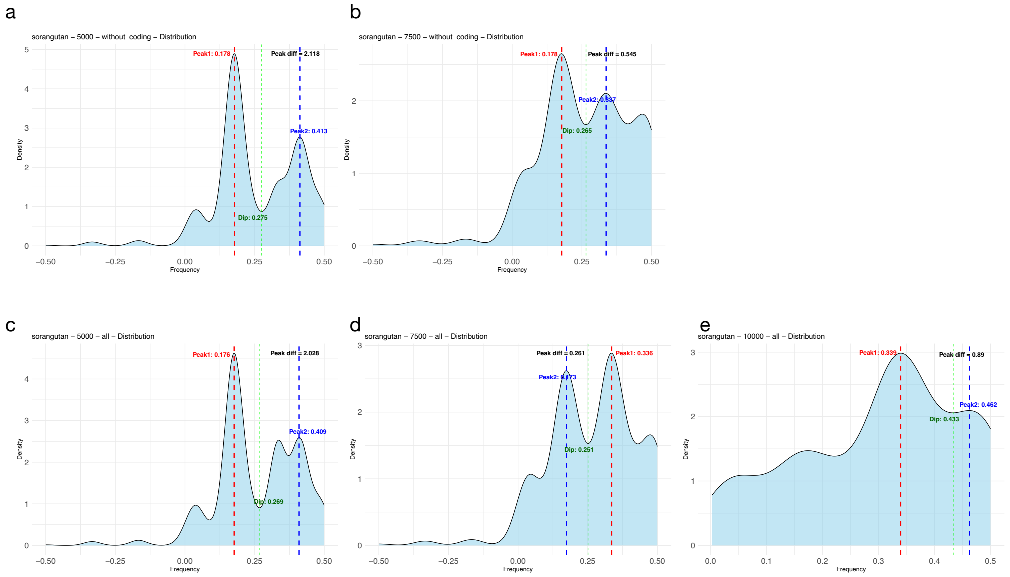


**Supplementary Figure 51:**

**Fourier frequency density plots of SCFRs at different length thresholds and categories in Sumatran orangutan.**

Kernel density plots show the distribution of frequencies derived from Fourier spectra of SCFRs. The x-axis represents frequency and the y-axis represents density. Red and blue dotted vertical lines indicate the two highest spectral peaks, while the green line marks the intervening minimum (dip). The annotated peak difference denotes the distance between the lowest and highest frequencies observed.

(a) Non-coding SCFRs with a minimum length of 5000 bp.

(b) Non-coding SCFRs with a minimum length of 7500 bp.

(c) Combined coding and non-coding SCFRs with a minimum length of 5000 bp.

(d) Combined coding and non-coding SCFRs with a minimum length of 7500 bp.

(e) Combined coding and non-coding SCFRs with a minimum length of 10000 bp.
